# Spatiotemporal proteomic profiling of cellular responses to NLRP3 agonists

**DOI:** 10.1101/2024.04.19.590338

**Authors:** L. Robert Hollingsworth, Priya Veeraraghavan, Joao A. Paulo, J. Wade Harper

## Abstract

Nucleotide-binding domain and leucine-rich repeat pyrin-domain containing protein 3 (NLRP3) is an innate immune sensor that forms an inflammasome in response to various cellular stressors. Gain-of-function mutations in NLRP3 cause autoinflammatory diseases and NLRP3 signalling itself exacerbates the pathogenesis of many other human diseases. Despite considerable therapeutic interest, the primary drivers of NLRP3 activation remain controversial due to the diverse array of signals that are integrated through NLRP3. Here, we mapped subcellular proteome changes to lysosomes, mitochondrion, EEA1-positive endosomes, and Golgi caused by the NLRP3 inflammasome agonists nigericin and CL097. We identified several common disruptions to retrograde trafficking pathways, including COPI and Shiga toxin-related transport, in line with recent studies. We further characterized mouse NLRP3 trafficking throughout its activation using temporal proximity proteomics, which supports a recent model of NLRP3 recruitment to endosomes during inflammasome activation. Collectively, these findings provide additional granularity to our understanding of the molecular events driving NLRP3 activation and serve as a valuable resource for cell biological research. We have made our proteomics data accessible through an open-access Shiny browser to facilitate future research within the community, available at: https://harperlab.connect.hms.harvard.edu/inflame/. We will display anonymous peer review for this manuscript on pubpub.org (https://harperlab.pubpub.org/pub/nlrp3/) rather than a traditional journal. Moreover, we invite community feedback on the pubpub version of this manuscript, and we will address criticisms accordingly.

## Introduction

The mammalian innate immune system comprises a diverse arsenal of intracellular signalling pathways that detect and eliminate existential threats to multicellular life. These pathways are carefully tuned to avoid hyper-activation, as spurious inflammation causes a myriad of diseases and contributes to ageing and neurodegeneration [1]. The intensity and duration of these immune responses fluctuate significantly depending on the specific stimulus that triggers them, the cell types involved and their physiological state, and the molecular pathway(s) activated. These variations can elicit a range of outcomes, such as the activation of inflammatory signalling pathways that ultimately result in cell death. Sensor proteins, responsible for initiating these responses, bind to conserved classes of pathogenic molecules, safeguard specific cellular proteins or pathways, and/or monitor cell state to trigger appropriate defensive countermeasures [2][3].

Several different sensor proteins form a supramolecular signalling complex, known as the canonical inflammasome, in response to danger-associated stimuli [4][5]. Upon recognizing their cognate ligand or cellular perturbation, these sensor proteins self-assemble the inflammasome to activate the cysteine protease caspase-1 either directly or by first recruiting the adaptor protein ASC[6][7]. Active caspase-1 cleaves cytokines to their bioactive forms and releases the N-terminal fragment of gasdermin-D (GSDMD) [8][9][10] [11][12]. Cleaved GSDMD then forms pores in membranes, releasing bioactive cytokines from cells [13][14]. Typically, these pores lead to cell death—referred to as pyroptosis [15]—and the recruitment of NINJ1 to the plasma membrane, which results in a distinct process of cell rupture that releases larger pro-inflammatory molecules from cells [16][17]. Numerous additional signals regulate each intermediate step in the inflammasome pathway to fine-tune inflammation, including at the level of sensor protein activation.

Different inflammasome sensor proteins detect distinct danger-associated molecular patterns in particular tissues relevant to their function [4][18]. Myeloid-lineage cells, for example, express several sensors, including the nucleotide-binding domain and leucine-rich repeat pyrin-domain containing protein 3 (NLRP3). Some cells that can undergo an NLRP3 signalling response first require a transcriptional “priming” stimulus, such as lipopolysaccharide (LPS), that transiently activates the NF-κB transcription factor [19][20]. NF-κB drives transcription of NLRP3 and other proteins in its activation pathway, causing large alterations to the cellular proteome [21]. While transcriptional priming is not required for NLRP3 inflammasome activation in some macrophage and microglia populations [22], it tunes the threshold of cellular stress required for pathway activation and the subsequent quantity of inflammatory cytokines released.

Posttranslational modifications to NLRP3 itself also threshold inflammasome activation. This posttranslational regulation is sometimes termed “licensing”, as licensing factors can include modifications that occur independent of transcriptional priming. For example, such modifications could modulate protein levels (e.g., by (de)ubiquitination) or the pool of NLRP3 more sensitive to activating stimuli (e.g., by phosphorylation, palmitoylation, etc.) [23][24][25]. Such threshold regulation is evident in human diseases resulting from aberrant inflammasome activity. Various disease-causing NLRP3 coding variants induce activation at different expression levels or licensing thresholds [26][27], leading to diverse clinical manifestations[1][28]. Given the importance of thresholding to NLRP3 activation, many modifications could govern NLRP3 activity in specific cell types and/or cell states, thereby playing a role in the pathogenesis of certain human diseases. **The full network of NLRP3 interactors that modulate its activation threshold remains unclear**.

The only licensing interaction frequently associated with NLRP3 activation involves binding the mitotic kinase NEK7 [29][30][31][32]. As cells progress through the cell cycle, NEK7 becomes sequestered [33], potentially diminishing the pool available for NLRP3 licensing and consequently reducing the cell’s capacity for an NLRP3 inflammasome response during mitosis [31]. While the rationale for damping NLRP3 by NEK7 sequestration remains unknown, it might be a protective mechanism against inadvertent inflammasome activation induced by programmatic disruptions in trafficking and disassembly of the Golgi during mitosis [34] [35]. Nonetheless, elevated concentrations of NLRP3 activating compounds, prolonged exposure to them, or particular human cell types circumvent the necessity for NEK7 licensing [36]. Subsequently, once expressed and licensed, NLRP3 maintains an inactive oligomeric state [37][38][39] until it senses one of many activating stimuli [24][40].

One large class of NLRP3 activating stimuli requires potassium efflux from the cell [41][42][43][44] and includes extracellular particulates like uric acid and cholesterol crystals [45][46][47], alum [48], silica [49], and protein aggregates [50][51][52]; extracellular ATP [53]; the potassium ionophore nigericin [53]; and lysosome disrupting agents like LLOMe [48]. Two additional NLRP3 activators, imiquimod (R837) and its derivative CL097, do not require cellular potassium efflux for an inflammasome response. Instead, they inhibit the mitochondrial respiratory complex I and are proposed to cause yet-uncharacterized endolysosomal disruption [54][55][56]. These diverse compounds disrupt cellular homeostasis through incompletely understood mechanisms but likely converge through cellular redistribution of the lipid phosphatidylinositol 4-phosphate (PI4P) and disruptions to intracellular trafficking [57][58][59].

Recent studies demonstrated that various inflammasome agonists disrupt retrograde trafficking and ER-endosome contact sites [58][59]. Contact sites between the ER and organelles where PI4P synthesis occurs, such as Golgi and endosomes [60][61][62], facilitate PI4P exchange to the ER down a concentration gradient [63][64]. Following this exchange, ER-localized Sac1 dephosphorylates PI4P to maintain the lipid gradient, which enables further counter current lipid exchange between membranes [65][66]. Sac1 might also dephosphorylate PI4P present on other organelles due to proximity at ER contact sites [67][68]. Consequently, when cells are unable to effectively transfer and turnover subcellular lipids, they accumulate PI4P, actin comets, and *trans*-Golgi proteins on endosomes [60][65]. **It remains unclear how various classes of NLRP3 agonists orchestrate these perturbations to cell state that ultimately activate the inflammasome**.

Endosomal PI4P enrichment induced by various inflammasome agonists eventually leads to the recruitment of NLRP3 [57][58][59]. NLRP3 associates with membranes peripherally through a polybasic region that binds PI4P, thereby localizing some of the protein to the Golgi and other endomembranes [36][37][57][69]. Posttranslational licensing modifications, such as palmitoylation or phosphorylation, might further enhance affinity between NLRP3 and membrane [36][70][71][72]. Deletion of the PI4P binding region inhibits NLRP3 activation, which can be restored by fusing an orthogonal PI4P binding domain, such as OSBP(PH), SidM, or SidC [73], to NLRP3 [57]. Additionally, suppressing endosomal PI4P accumulation or promoting its dephosphorylation inhibits NLRP3 activation [57][58]. Further supporting endosomes as platforms for inflammasome activation, ceramide-rich vesicles were observed to permeate the filamentous inflammasome framework by cryo-ET and immunofluorescence microscopy [74]. **The precise identity of these PI4P-rich endosomes and how NLRP3 undergoes necessary conformational changes on them remain open questions**.

To better understand how activators of the NLRP3 inflammasome disrupt cellular homeostasis, we generated a proteomic inventory of key organelles implicated in inflammasome activation in the resting state and stimulated with two different classes of inflammasome agonists: the potassium efflux-dependent compound nigericin, and the potassium efflux-independent compound CL097. We find that inflammasome agonists cause large-scale remodelling of the endocytic pathway, consistent with intracellular trafficking disruption [58][59]. The localization changes observed were largely independent of alterations in total protein levels, emphasizing the importance of evaluating spatial dynamics rather than just protein abundance alone.

Subsequently, we mapped how mouse Nlrp3 traffics in response to these compounds with time-resolved proximity biotinylation. We find that Nlrp3 rapidly loses contact with the ER to favour interactions with actin- and endosome-related proteins, supporting a model of inflammasome nucleation from PI4P-rich endosomes following the loss of ER-endosome contact sites [58]. Furthermore, by conducting parallel proximity biotinylation experiments with a PI4P lipid biosensor, we assemble a high-confidence Nlrp3 interactome resource. Our spatiotemporal proteomic inventory provides a hypothesis-generating engine for understanding the perturbations in cell state that are detected by the NLRP3 inflammasome, its subsequent recruitment to endosomes, and broader aspects of cellular trafficking.

## Results

### Inflammasome agonists do not cause global proteome remodelling

We first asked whether different classes of inflammasome agonists cause global proteome remodelling (e.g., stabilization or degradation) by treating HEK293T cells with either nigericin (a potassium ionophore) or CL097 (an NQO2 inhibitor with poorly characterized endolysosomal effects). Few proteins significantly changed in abundance, but notably, CDC25A and CDC25B decreased in response to both inflammasome agonists (Log_2_FC < −1.3; Figure 2A-C). CDC25C loss, however, was much more modest (Log_2_FC < −0.2). These phosphatases coordinate entry into the cell cycle by activating cyclin-dependent kinases and their degradation can prevent cell cycle progression [75]. NEK7 sequestration can damp the NLRP3 inflammasome response during mitosis [31][36], and thus, the cellular stresses induced by NLRP3 agonists might conversely promote cell cycle arrest, which can occur in response to diverse cellular stressors [76]. No additional protein complexes annotated by CORUM [77] or BioPlex [78] were enriched. Moreover, only modest protein accumulation of ER and mitochondrial proteins occurred in aggregate (Figure 2D, Figure 2—Supporting Figure 1).

**Figure 1.**
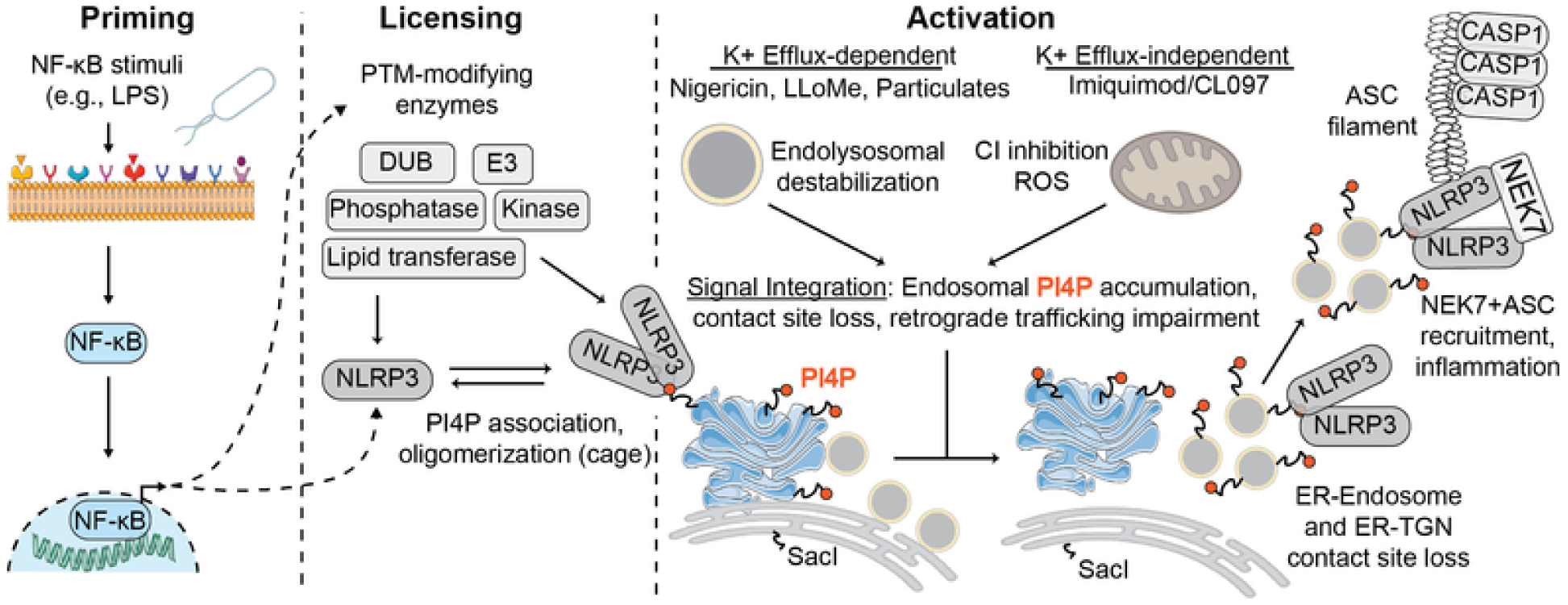
Cellular mechanisms of NLRP3 inflammasome activation, which include transcriptional priming, post-translational licensing, and activation steps (see introduction).

**Figure 2.**
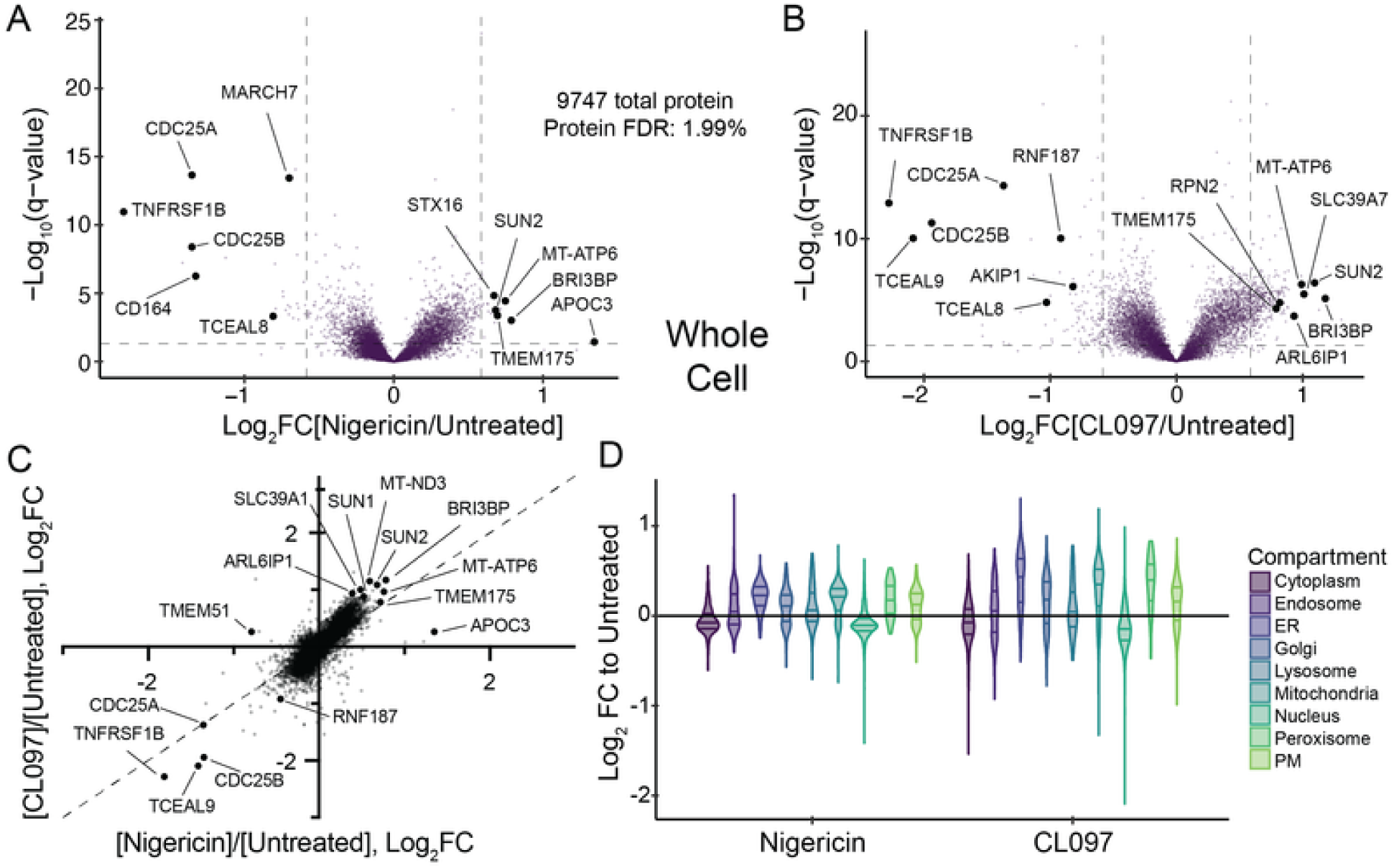
Whole-cell proteome effects of NLRP3 agonists. (A-B) Whole-cell proteomics volcano plot for (A) nigericin (20 μM, 30 min)-treated or (B) CL097 (75 μg/mL, 1 h)-treated versus untreated HEK293T cells. *P* values were calculated from the Student’s t-test (two sided) and adjusted for multiple hypothesis correction using the Benjamini–Hochberg approach (q-value) with MSstats. Data represent n=5 biological replicates (see Figure 2—Supporting Figure 1 for TMTplex design). (C) Comparison of log_2_ fold change (FC) values in response to nigericin (x-axis) or CL097 (y-axis) treatment. Identical changes in both conditions lie on the dashed y-x axis. (D) Violin plots depicting the log_2_FC values across the indicated subcellular compartments for the indicated whole cell proteomics comparison (nigericin/untreated or CL097/untreated)

Inflammasome signalling occurs rapidly, and recent studies associated intracellular trafficking disruptions with many different NLRP3 activating stimuli [57][58][59]. To investigate how inflammasome agonists alter the composition of cellular organelles, we employed organelle immunoprecipitation (organelle-IP) strategies. These approaches rely on protein affinity handles with well-defined subcellular localization, facilitating rapid organelle-IP on magnetic beads [79]. Such affinity handles have been designed and characterized to isolate a range of organelles, including mitochondria [79], peroxisomes [80], synaptic vesicles [81], early/sorting endosomes [82], and Golgi [83]. Recently, researchers at the Chan Zuckerberg Initiative further expanded the organelle-IP toolkit to include many other cellular compartments [84]. By employing these techniques, we sought insights into how organellar dynamics contribute to NLRP3 activation. Importantly, there are two fundamental limitations of our approach: 1) we only assess changes at the protein level, but lipid and metabolite flux likely contribute to NLRP3 activation, and 2) protein handles could bias organelle isolations to particular subpopulations, as recently demonstrated for TMEM192 [85].

### Nigericin induces CASM on lysosomes, and both inflammasome agonists cause subtle mitochondrial proteome changes

Lysosome-damaging agents such as uric acid crystals and LLoMe activate the NLRP3 inflammasome [86]. To uncover general features of lysosomal damage associated with the inflammasome response, we characterized how different NLRP3 agonists remodel the lysosomal proteome using LysoIP [87]. Nigericin and CL097 do not disrupt lysosomal targeting of the LysoTag (TMEM192-3xHA) in HEK cells (HEK293^L^) we previously tagged at the endogenous locus [88] (Figure 3—Supporting Figure 1A). We performed anti-HA immunoprecipitations from untagged control HEK293 cells, untreated HEK293^L^ cells, nigericin-treated HEK293^L^ cells, and CL097-treated HEK293^L^ cells followed by TMT-based quantitative proteomics of the eluent (Figure 3—Supporting Figure 1). Compared to control “untagged” cells, pulldowns with cells expressing the LysoTag enriched for lysosome annotated proteins specifically, validating our approach (Figure 3—Supporting Figures 1B-D).

**Figure 3.**
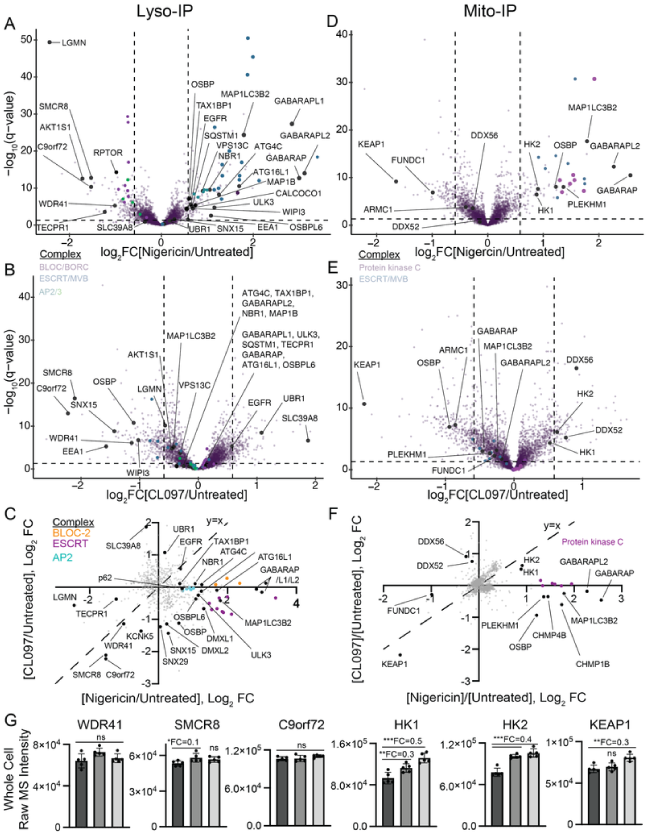
Lysosomal and mitochondrial proteome effects of NLRP3 agonists. (A-B) LysoIP volcano plots for (A) nigericin (20 μM, 30 min)-treated or (B) CL097 (75 μg/mL, 1 h)-treated versus untreated HEK293^L^ cells. Protein complexes are colored as indicated. *P* values were calculated from the Student’s t-test (two sided) and adjusted for multiple hypothesis correction using the Benjamini–Hochberg approach (q-value) with MSstats. Data represent n=3 or 5 biological replicates. For the full TMTplex experimental setup, see Figure 3 —Supporting Figure 1. (C) Comparison of LysoIP log_2_ fold change (FC) values in response to nigericin (x-axis) or CL097 (y-axis) treatment, filtered for proteins localized to lysosomes (any 293^L^/293: q < 0.05, Log_2_ FC > 0.5). Identical changes in both conditions lie on the dashed y-x axis. Protein complexes are colored as indicated. (D-E) MitoIP volcano plots for (A) nigericin (20 μM, 30 min)-treated or (B) CL097 (75 μg/mL, 1 h)-treated versus untreated HEK293^L^ cells. Protein complexes are colored as indicated. *P* values were calculated from the Student’s t-test (two sided) and adjusted for multiple hypothesis correction using the Benjamini–Hochberg approach (q-value) with MSstats. Data represent n= 4 biological replicates. For the full TMTplex experimental setup, see Figure 3—Supporting Figure 3. (F) Comparison of MitoIP log_2_FC values in response to nigericin (x-axis) or CL097 (y-axis) treatment, filtered for proteins localized to mitochondria (any 293T^M^/293T^Control^: q < 0.05, Log_2_ FC > 0.5). Identical changes in both conditions lie on the dashed y-x axis. Protein complexes are colored as indicated. (G) Whole cell proteomics data (see Figure 1 and Figure 1—Supporting Figure 1) for the indicated proteins, either untreated (dark grey), nigericin-treated (medium-grey), or CL097-treated (light grey). Log_2_FC values over the untreated condition are indicated for significant changes. ns, not significant, *q<0.05, **q<0.01, **q<0.001. Error bars represent ± SD of 5 replicates.

Few proteins, GO terms [89], BioPlex interactomes [78], and organelle protein sets demonstrated similar patterns in both treatment conditions compared to untreated cells (Figures 3A-C, Figure 3—Supporting Figure 2). Notably, both treatments caused reduced lysosomal localization of the C9orf72 complex (C9orf72, SMCR8, and WDR41), which localizes to lysosomes and the cytoplasm [90][91]. Mutations in this complex lead to familial amyotrophic lateral sclerosis (ALS) and frontotemporal dementia (FTD) [92], and recent studies demonstrated that genetic ablation of the C9orf72 complex caused sustained NLRP3 inflammasome activation in microglia [93][94]. Previous lysosomal proteomics of damaged lysosomes, induced by either LLoMe or GPN [95][96], only identified one protein of this complex [88][97]. Furthermore, we were unable to identify experimental conditions for C9orf72 immunofluorescence. Therefore, we cannot conclude whether C9orf72 lysosomal depletion is a general characteristic of lysosomal damage or plays a role in lysosomal repair.

Nigericin treatment induces conjugation of ATG8 to single membranes (CASM) on lysosomes [98][99][100]. Indeed, nigericin treatment, but not CL097, increased lysosomal localization of autophagy receptors (TAX1BP1, SQSTM1/p62, NBR1, CALCOCO1), ATG8s (GABARAP, GABARAPL1, GABARAPL2, MAP1LC3B2), autophagy-associated machinery (ATG4C, WIPI3/WDR45B), and ATG16L1, a component required for ATG8 conjugation in both autophagy and CASM (Figure 3A-C) [101][102][103]. Additionally, nigericin treatment led to lysosomal recruitment of the ESCRT machinery (Figure 3C), possibly due to endolysosomal damage [104]. We did not, however, detect enrichment of proteins associated with lysosomal membrane permeation (e.g., OSBPL9/10/11, PI4K2A [105]); instead, three proteins associated with lipid transfer were enriched: OSBP, OSBPL6, and VPS13C. The substantial differences between nigericin and CL097 treatments suggest that the cellular state required for NLRP3 inflammasome activation occurs downstream of or independently from lysosomal damage.

Imiquimod and its derivative CL097 activate the inflammasome in part by disrupting mitochondrial and metabolic homeostasis [55][56]. Additionally, inhibition of the electron transport chain dampens NLRP3 activation by multiple agonists [56], and mitochondria have been proposed as a site of inflammasome activation [106][107][108][109]. We thus profiled how inflammasome agonists remodel the mitochondrial proteome. First, we expressed the MitoTag [79] (3xHA-GFP-OMP25) in HEK293T cells (HEK293T^M^), sorted for stable low expressors by FACS, and validated tag localization by immunofluorescence (Figure 3—Supporting Figure 3A). We then performed MitoIPs followed by quantitative proteomics with untagged HEK293T cells, untreated HEK293T^M^ cells, nigericin-treated HEK293T^M^ cells, and CL097-treated HEK293T^M^ cells (Figure 3—Supporting Figure 3B). Compared to control “untagged” cells, pulldowns with cells expressing the MitoTag enriched for mitochondrial annotated proteins specifically, validating our approach (Figure 3—Supporting Figure 3C). ER and peroxisome-associated proteins were also slightly enriched, in line with previous mitochondrial isolations by MitoIP [110].

Like lysosomes, CL097 and nigericin share few common mitochondrial proteome perturbations (Figure 3D-F, Figure 3—Supporting Figure 4). Additionally, far fewer mitochondrial proteins were perturbed in response to these compounds compared to a previous study that treated macrophages with LLoMe [111], a compound that ruptures lysosomes and activates NLRP3. Nigericin and CL097 both deplete KEAP1, a component of a Cullin E3 ubiquitin ligase complex that targets the transcription factor Nrf2 for degradation. Upon oxidative stress, KEAP1 dissociates from Nrf2, allowing Nrf2 to translocate to the nucleus and stimulate an antioxidative gene expression profile [112]. As Nrf2 depletion can dampen NLRP3 signalling in some contexts [113], and many NLRP3 activating compounds induce cellular ROS [43][49][55][114], we did not investigate further.

**Figure 4.**
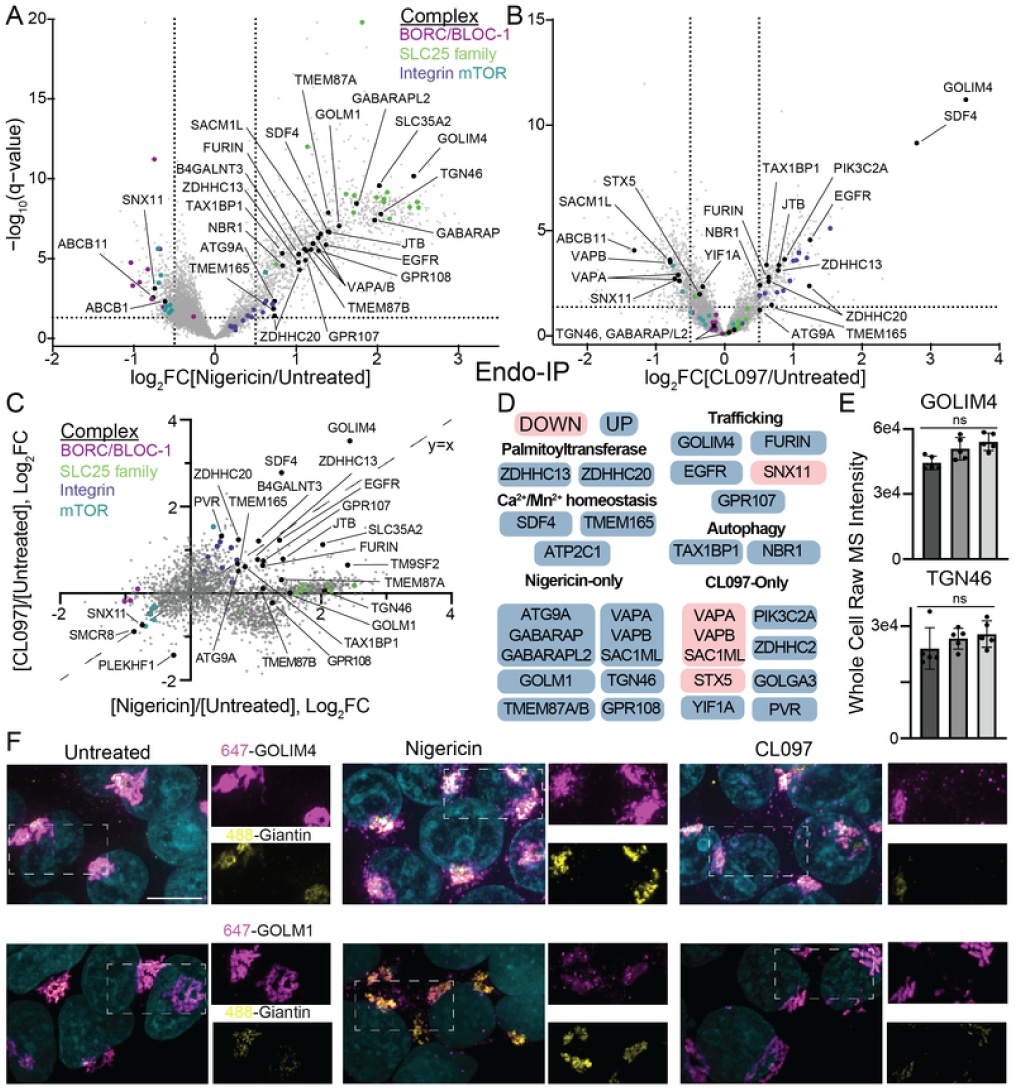
Endosomal proteome effects of NLRP3 agonists. (A-B) EndoIP volcano plots for (A) nigericin (20 μM, 30 min)-treated or (B) CL097 (75 μg/mL, 1 h)-treated versus untreated HEK293T^EG^ cells. Protein complexes are colored as indicated. *P* values were calculated from the Student’s t-test (two sided) and adjusted for multiple hypothesis correction using the Benjamini–Hochberg approach (q-value) with MSstats. Data represent n=3 or 5 biological replicates. For the full TMTplex experimental setup, see Figure 4—Supporting Figure 1. (C) Comparison of EndoIP log_2_ fold change (FC) values in response to nigericin (x-axis) or CL097 (y-axis) treatment, filtered for proteins localized to endosomes (293T^EG^/293T: q < 0.05, Log_2_FC > 0.5). Identical changes in both conditions lie on the dashed y-x axis. Protein complexes are colored as indicated. Annotated proteins change significantly in at least one treatment condition (nigericin or CL097 293T^EG^/293T^EG^: q < 0.05). (D) Summary of proteins that change in response to inflammasome agonist treatments. Selected proteins with q < 0.05, Log_2_ FC > 0.5 are included. (E) Whole cell proteomics data (see Figure 1 and Figure 1—Supporting Figure 1) for the indicated proteins, either untreated (dark grey), nigericin-treated (medium-grey), or CL097-treated (light grey). ns, not significant. Error bars represent ± SD of 5 replicates. (F) Nigericin and CL097 treatment both increase GOLIM4 vesicle staining, whereas only nigericin increases GOLM1 vesicle staining. HEK293T cells were treated with nigericin (20 μM, 30 min) or CL097 (75 μg/mL, 1 h), fixed, and immunostained with the indicated antibody and Hoechst. Maximum intensity projection images (z=8 μM, 29 steps), representative of n > 6 fields of view in at least two independent experiments. Scale bar (top left panel), 10 μm.

Additionally, in contrast to a recent report [106], we find that both hexokinase 1 and 2 (HK1, HK2) accumulate, rather than dissociate from, the mitochondria upon inflammasome stimulation (Figures 3D-E). This effect could be due to different metabolic states between cell types, particularly with NF-κB stimulation in macrophages, so it warrants further study. Only nigericin treatment accumulated proteins that could indicate membrane damage, including ATG8s (GABARAP, GABARAPL2, MAP1LC3B2), CHMP4B, and OSBP (Figure 3F). Beyond ATG8s, there was no additional evidence of a selective mitophagy response to the inflammasome agonists as previously reported [115], possibly due to a lack of PARKIN expression in the HEK293T cell line. Additionally, several subunits of protein kinase C accumulated following nigericin treatment. These localization changes at the lysosome and mitochondria were mainly not reflected in global protein levels, with only modest increases in the levels of HK1 and HK2 in response to inflammasome agonists (Figure 3G).

### Inflammasome agonists induce large changes in endosomal trafficking

NLRP3 co-localizes with EEA1-positive endosomes upon treatment with nigericin or CL097/imiquimod [58] [59]. Consequently, endosomes are a proposed site for NLRP3 trafficking and activation. We recently developed a system to IP intact early/sorting endosomes for quantitative proteomics using 3xFLAG-tagged EEA1 (EndoIP) [82]. Overexpression of FLAG-EEA1 can lead to heterogeneously localized protein ([see negative data](https://harperlab.pubpub.org/pub/nlrp3#n0k1i1xwi40)), so we engineered a 3xFLAG tagged EEA1 at its endogenous locus in HEK293T cells. We first isolated single clones that demonstrated uniform FLAG endosomal localization (HEK293T^E^). Subsequently, we also incorporated the GolgiTag [83] at its endogenous locus (TMEM115-3xHA), creating a single clonal cell line for both EndoIP and GolgiIP (HEK293T^EG^, Figure 4—Supporting Figure 1A). We then confirmed that 3xFLAG-EEA1 remained associated with endosomes after stimulation with nigericin and CL097 (Figure 4—Supporting Figure 1B).

We performed EndoIP followed by quantitative proteomics from untagged HEK293T cells, untreated HEK293T^EG^ cells, nigericin-treated HEK293T^EG^ cells, and CL097-treated HEK293T^EG^ (Figures 4A-B, Figure 4—Supporting Figure 1C). Compared to control cells lacking the EndoTag, EndoIP specifically enriched endosome, lysosome, and plasma membrane annotated proteins that likely traffic through EEA1-positive early/sorting endosomes (Figure 4—Supporting Figure 1D). Principal component and organelle protein set analyses indicate that nigericin caused a broad disruption of endosomal homeostasis (Figure 4—Supporting Figures 1D-G). Nigericin disrupts endolysosomal pH and profoundly increases endosome size [116] [117], which could, in part, explain the large skew in protein distribution upon nigericin treatment (Figure 4C). In contrast, CL097 only caused a modest reduction in ER protein enrichment across all organellar annotations (Figure 4—Supporting Figures 1D-G), which could indicate disrupted ER-endosome contact sites, as recently demonstrated [58].

Nigericin and CL097 treatments induced endosomal enrichment of several proteins that cycle in the endolysosomal system, such as FURIN, EGFR, and GOLIM4 (Figures 4C-D). Additionally, many proteins involved in Shiga toxin and/or ricin trafficking, including GOLIM4, TMEM165, JTB, SLC35C1, SLC35A2, and TM9SF2 [118][119][120][121], were enriched on endosomes in both conditions (Figure 4C). These findings align with recent studies demonstrating that inflammasome agonists disrupt retrograde trafficking pathways, as evidenced by transferrin, EGF, and Shiga toxin trafficking assays [58][59].

Nigericin induced the accumulation of FU7TM family proteins (GPR107, GPR108, TMEM87A, TMEM87B), TGN46 (also known as TGOLN2; TGN38 in mouse), GOLM1 (also known as GP73 or GOLPH2), and GOLIM4 (also known as GOLPH4 or GPP130) within endosomes. TGN46, GOLIM4, and GOLM1 play roles in bypassing lysosomal trafficking, thereby facilitating the retrieval of endosomal cargo back to the Golgi apparatus [122][123]. Among these proteins, only GOLIM4 also exhibited enrichment in EEA1-positive endosomes after CL097 treatment (Figure 4C). Importantly, enrichment was not a result of total proteome changes, as the overall levels of GOLIM4 and TGN46 were unchanged (Figure 4E). Moreover, nigericin distinctly accumulated GOLIM4 and GOLM1 within vesicular structures by immunofluorescence, whereas CL097 induced solely GOLIM4 endosomal enrichment (Figure 4F), further validating these findings.

GOLIM4 becomes trapped in endosomes and ultimately degraded by lysosomes on endolysosomal pH disturbance (e.g., monensin or bafilomycin treatment), manganese treatment, and retro-2 treatment [122][124] [125][126]. Interestingly, these same compounds either activate or potentiate the NLRP3 inflammasome response in specific contexts [58][59][127][128]. The particular disruptions to endosomal trafficking required for NLRP3 activation likely occur upstream or in parallel to GOLIM4 disruption, as GOLIM4 knockout itself did not impact inflammasome activation in THP-1 macrophages ([see negative data](https://harperlab.pubpub.org/pub/nlrp3/release/1#n79240mp24c)). In summary, inflammasome agonists caused large-scale changes in retrograde trafficking pathways, as recently described [58][59], which could be due to their known impact on endolysomal pH [55][117][129].

### Inflammasome agonists cause large-scale Golgi proteome changes

A pool of NLRP3 localizes to the Golgi following transcriptional priming and prior to inflammasome activation [36][57]. Furthermore, disrupting Golgi pH homeostasis either activates or potentiates inflammasome signalling in specific cell types [59][130]. To investigate how Golgi stress might contribute to inflammasome activation, we characterized how nigericin and CL097 remodel the Golgi proteome. We chose the recently developed Golgi-IP approach [83], and found that the GolgiTag (TMEM115-3xHA) remained properly localized following treatment with nigericin or CL097 in our HEK293T^EG^ cells (Figure 5—Supporting Figure 1A).

**Figure 5.**
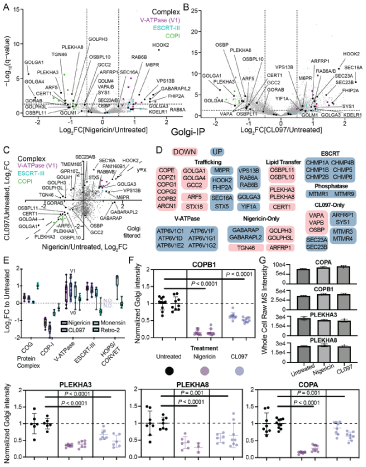
Golgi proteome effects of NLRP3 agonists. (A-B) GolgiP volcano plots for (A) nigericin (20 μM, 30 min)-treated or (B) CL097 (75 μg/mL, 1 h)-treated versus untreated HEK293T^EG^ cells. Protein complexes are colored as indicated. *P* values were calculated from the Student’s t-test (two sided) and adjusted for multiple hypothesis correction using the Benjamini–Hochberg approach (q-value) with MSstats. Data represent n=3 or 4 biological replicates. For the full TMTplex experimental setup, see Figure 5—Supporting Figure 1. (C) Comparison of GolgiIP log_2_ fold change (FC) values in response to nigericin (x-axis) or CL097 (y-axis) treatment, filtered for proteins localized to the Golgi (293T^EG^/293T: q < 0.05, Log_2_ FC > 0.5). Identical changes in both conditions lie on the dashed y-x axis. Protein complexes are colored as indicated. Annotated proteins change significantly in at least one treatment condition (nigericin or CL097 293T^EG^/293T^EG^: q < 0.05). (D) Summary of proteins that change in response to inflammasome agonist treatments. Selected proteins with q < 0.05, Log_2_FC > 0.5 are included. (E) GolgiIP log_2_FC values for individual members of the indicated protein complexes in response to nigericin (20 μM, 30 min), CL097 (75 μg/mL, 1 h), monensin (10 μM, 2 h), or retro-2 (25 μM, 2 h) compared to the untreated condition. Note that fold changes between the nigericin/CL097 and monensin/retro-2 conditions cannot be directly compared due to different TMTplex experiments. V1 subunits of the V-ATPase were consistently those enriched on the Golgi following treatment. For the monensin/retro-2 TMTplex experimental setup and volcano plots, see Figure 5—Supporting Figure 4. (F) CL097 and nigericin deplete Golgi localization of COPB1, COPA1, PLEKHA3, and PLEKHA8. HEK293T cells were treated with nigericin (20 μM, 30 min) or CL097 (75 μg/mL, 1 h), fixed, and immunostained with the indicated antibody, a Golgi marker, and Hoechst. The Golgi marker was used as a mask, and the ratio of the intensities of the indicated protein to the Golgi marker were quantified within the mask using CellProfiler [189]. These values were then normalized to the average untreated condition for a given antibody. Maximum intensity projection images (z=8 μM, 29 steps), representative of n > 6 fields were quantified from two independent experiments. *P* values were calculated using a 2-way ANOVA with Tukey’s multiple comparison test, as indicated. (G) Whole cell proteomics data (see Figure 1 and Figure 1—Supporting Figure 1) for the indicated proteins, either untreated (dark grey), nigericin-treated (medium-grey), or CL097-treated (light grey). All comparisons are not significant.

We performed anti-HA immunoprecipitation from untagged control HEK293T cells, untreated HEK293T^EG^ cells, nigericin-treated HEK293T^EG^ cells, and CL097-treated HEK293T^EG^ cells in biological quadruplicate followed by Tandem Mass Tagging (TMT)-based proteomics of the eluent (Figure 5—Supporting Figures 1B-C). As expected, GolgiIP specifically enriched Golgi-associated proteins compared to control cells lacking the GolgiTag (Figure 5—Supporting Figure 1D). However, depending on the particular annotation list used, *trans-* Golgi (TGN) proteins were either not enriched or enriched at levels similar to bulk Golgi annotations. Therefore, changes in GolgiIP might also reflect movement within the Golgi stack itself (e.g., *cis-* versus *trans*-Golgi) if one compartment IPs more efficiently than the other, but we could not validate either claim.

Principal component and organelle protein set analyses revealed that CL097 induced widespread disruption of Golgi homeostasis across various protein classes, whereas nigericin treatment did not perturb any specific organellar protein set in bulk (Figure 5—Supporting Figure 1E-F). Among proteins enriched by GolgiIP, inflammasome agonist treatment did not cause bulk changes in TGN, Golgi, and ERGIC annotations [84] (Figure 5—Supporting Figure 2A). Thus, in agreement with a recent report, the TGN was largely intact based on proteome analysis [58]. However, several TGN golgins were depleted with nigericin and CL097, including GOLGA1 (Golgin-97), GOLGA4 (p230/Golgin-245), and GCC2 (GCC185) (Figures 5A-D). These proteins tether vesicles at the TGN, promoting endosome-to-TGN retrograde transport [131]. Treatment caused a slight Golgi depletion and accompanying endosomal enrichment of annotated [132] GCC1 cargo (Figure 5—Supporting Figures 2B-C). GOLGA1 cargo was not impacted in bulk (Figure 5—Supporting Figures 2B-C). As previously discussed, these changes could reflect differences in Golgi stack positioning if the IPs bias towards proximal Golgi compartments, but regardless support a model where inflammasome agonists disrupt endosome-to-TGN retrograde transport [58][59].

Several proteins associated with trafficking (e.g., M6PR, RAB6) and lipid transfer at organellar contact sites (e.g., CERT1, OSBPL10, PLEKHA3) were disrupted (Figure 5D). This is consistent with inflammasome agonists disrupting contact sites, lipid distribution, and trafficking [57][58][59]. Nigericin, but not CL097, depleted TGN46 from the Golgi (Figures 5C-D), concomitant with its enrichment in EEA1-positive endosomes (Figures 4C-D). Despite significant enrichment of GOLIM4 in endosomes (Figures 4C-D), depletion from the Golgi was not prominent (Figures 5A-B). This observation could suggest that the steady-state pool of GOLIM4 in the endosomes is small, the pool in the Golgi is large, or a combination of both factors.

We leveraged GO-term analysis and BioPlex interactome data [78] to discover protein pathways/complexes affected by nigericin or CL097 (Figure 5—Supporting Figure 3). Both agonists caused Golgi accumulation of ESCRT-III machinery and V1 (but not V0) subunits of the V-ATPase (Figure 5D). Increased localization of the V-ATPase to the Golgi could indicate that both compounds disrupt Golgi pH. While the effect of nigericin and CL097 on Golgi pH has not been tested directly, it would be consistent with their effects on endolysosomal pH [55][117][129]. Moreover, chemical inhibition of the V-ATPase with bafilomycin A1 or concanamycin A exacerbates but is not sufficient for NLRP3 inflammasome formation in human macrophages [128], and STING-mediated Golgi pH disruption activates NLRP3 in specific contexts [130][133][134].

Nigericin and CL097 treatment depleted the entire COPI complex from the Golgi; nigericin additionally depleted the COPI adaptors [135] GOLPH3 and GOLPH3L (Figures 5A-D). COPI-coated vesicles transport cargo retrogradely between Golgi stacks and from the Golgi to the ER [136]. There was a concomitant enrichment of COPI cargo, but not COPII cargo, on the Golgi in response to CL097 treatment (Figure 5—Supporting Figure 2D). This is consistent with build-up due to impaired COPI-mediated retrograde trafficking of cargo to the ER. We did not detect COPI cargo enrichment following nigericin treatment, but this difference could be due to different timescales of the experimental conditions (30-minute nigericin treatment versus 60-minute CL097 treatment).

We performed an additional GolgiIP experiment with monensin and retro-2, compounds that are insufficient to activate NLRP3 but potentiate CL097/imiquimod inflammasome responses [59] (Figure 5—Supporting Figure 4). Monensin, but not retro-2, accumulated ESCRT-III at the Golgi and depleted COPI (Figure 5E). While the V-ATPase accumulated in all four treatment conditions (Figure 5E), CL097 treatment did not cause Golgi enrichment of ATG8 proteins (e.g., GABARAP, GABARAPL2, MAP1LC3B2; Figure 5C). Therefore, while nigericin, retro-2, and monensin might cause autophagy or CASM at the Golgi, CL097 likely does not. Only retro-2 treatment enriched members of the COG (conserved oligomeric Golgi) complex, which orchestrates retrograde trafficking at the Golgi [137][138]. Interestingly, CL097, but not nigericin, enriched SEC16A, SEC23A, and SEC23B. These proteins reside at ER exit sites and function in COPII biogenesis to maintain the anterograde flow of proteins [139]. Retro-2 blocks certain endosome-TGN retrieval pathways [140] by targeting SEC16A [124], which could indicate similarities in the trafficking disruptions induced by retro-2 and CL097.

We validated the Golgi depletion of COPA, COPB1, PLEKHA3 (FAPP1), and PLEKHA8 (FAPP2) by both nigericin and CL097 using immunofluorescence (Figure 5F and Figure 5—Supporting Figure 5). Additionally, nigericin, but not CL097, affected TGN46 and GPR107 localization, whereas the Golgi localization of TMEM165 was not noticeably affected by either treatment (Figure 5—Supporting Figure 5). Despite the large changes in localization for COPA, COPB1, PLEKHA3, and PLEKHA8, the global levels of these proteins did not significantly change following nigericin and CL097 treatment (Figure 5G). Despite these localization changes, we found that bulk knockouts of either PLEKHA3 or PLEKHA8 in THP-1s, or their clonal knockout in iBMDMs, was not sufficient to affect NLRP3 signalling ([see negative data](https://harperlab.pubpub.org/pub/nlrp3/release/1#n79240mp24c)). In summary, inflammasome agonists induce large-scale changes to proteins involved in retrograde transport, contact site maintenance, and lipid transfer.

### Spatiotemporal trafficking of NLRP3 and PI4P-associated proteins in response to nigericin and CL097

During activation, NLRP3 undergoes a conformational change from an inactive oligomer to a NEK7-associated inflammasome nucleator [32][37][38][39][141]. The active conformation of NLRP3 recruits the adaptor protein ASC (PYCARD) through homotypic interactions facilitated by a pyrin domain (PYD) present on each protein [6][7][142]. Subsequently, ASC condenses into a single, dense filamentous mesh known as the “ASC speck”, mediated by its caspase activation and recruitment domain (CARD). The CARD of ASC also recruits signalling molecules like caspase-1, thereby amplifying the inflammasome response [143][144][145] [146]. Upon activation, NLRP3 and ASC colocalize with PI4P, endosomal markers, and ceramide-containing vesicles, which suggest the inflammasome assembles from endosomes [57][58][59][74]. **However, the precise mechanism by which PI4P-rich endosomes recruit NLRP3, their specific identity, and how they facilitate the necessary conformational changes in NLRP3 to initiate inflammasome assembly remain unclear**.

To monitor how NLRP3 traffics to the inflammasome assembly site, we employed an APEX2-based proximity biotinylation approach [147] (Figure 6A). Initially, we reconstituted *Nlrp3*^-/-^ immortalized bone marrow-derived macrophages (iBMDMs) with various Nlrp3 expression constructs linked to APEX2 (e.g., Nlrp3-APEX-FLAG-IRES-GFP), driven by a portion of the UBC promoter. Using FACS, we isolated stable cell populations wherein Nlrp3 expression levels closely resembled those of LPS-stimulated wild-type (WT) iBMDMs (Figure 6—Supporting Figure 1A-B). We confirmed that these cells exhibited reconstituted inflammasome activity, whereas specific mutations, such as truncation of the PYD, impaired this activity, as evidenced by LDH release and ASC speck formation (Figure 6—Supporting Figures 1C-E).

**Figure 6.**
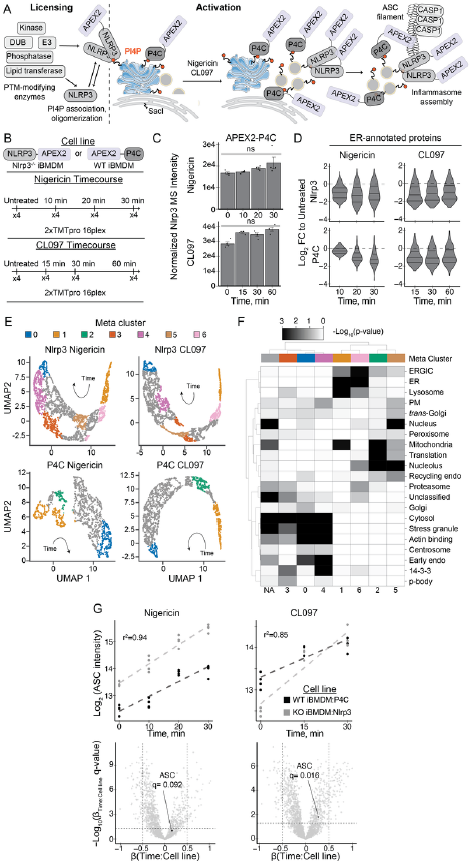
Timecourse proximity labeling of Nlrp3 and P4C during inflammasome activation. (A) Schematic of APEX2-based proximity biotinylation during Nlrp3 activation. While the lentivirally-reconstituted APEX2 fusions are constitutively expressed, LPS leads to proteome changes that could include enzymes that modify Nlrp3. Nearby proteins are indiscriminately biotinylated within a ∼20 nm falloff radius by APEX2, which could capture trafficking events that occur during inflammasome activation. The P4C biosensor binds PI4P and thus serves as a location-specific control. (B) Summary of the separate 4 TMTplex experiments. *Nlrp3*^-/-^ iBMDMs were reconstituted with Nlrp3-APEX2, primed with LPS (1 μg/mL, 4 h), and stimulated with either nigericin (20 μM; 0, 10, 20, 30 min) or CL097 (75 μg/mL; 0, 15, 30, 60 min). WT iBMDMs were reconstituted with APEX2-P4C, primed with LPS (1 μg/mL, 4 h), and stimulated with either nigericin (20 μM; 0, 10, 20, 30 min) or CL097 (75 μg/mL; 0, 15, 30, 60 min). Complete TMTplex experimental designs for each condition are in Figure 6—Supporting Figures 3-6. (C) Nlrp3 MS intensity values for APEX2-P4C timecourse experiments. Nlrp3 proximity to P4C-APEX2 does not change over the course of the experiment. *P* values were calculated from the Student’s t-test (two sided) and adjusted for multiple hypothesis correction using the Benjamini–Hochberg approach (q-value) with MSstats. Data represent n=4 biological replicates. ns, not significant. (D) Violin plots depicting the log_2_ fold change (FC) values for ER proteins detected in each experiment. Notably, proximity to ER decreases rapidly in all four experiments. Complete organellar annotations for each condition are in Figure 6—Supporting Figures 3-6. (E) ANOVA-significant proteins in each APEX2 dataset were clustered with the Leiden algorithm. These clusters were then compared across the four different datasets with the Jaccard index to find like-clusters between datasets (meta-clusters). These meta clusters are colored as annotated on UMAP embeddings of the data. (F) Heatmap of organellar enrichment for proteins in each meta cluster, colored as in (E). ER and lysosomal proteins enrich in early timepoint meta clusters, whereas actin binding proteins enrich in late timepoint clusters. Organellar annotations reported in Hein*, Peng*, Todorova*, McCarthy*, Kim*, and Liu* *et al.* (graph-based annotations) [84]. *P* values were calculated from the Fisher’s exact test for enrichment (one-sided). (G) Normalized intensity values with MSstats were fitted with generalized linear models. The effect of the treatment (β1), APEX2 bait condition (β2), and their interaction (β1β2) over the timecourse were tested. The difference in slopes represents the β1β2 interaction, which corresponds to differences in the behavior of APEX bait condition in response to inflammasome agonists (β1β2 = β_Time:Cell line_). Normalized MS intensity values for Asc over the linear range of the timecourses (CL097 1 h excluded) are shown, in addition to volcano plots of the β1β2 interaction term. *P* values were calculated from the Student’s t-test (two sided) and adjusted for multiple hypothesis correction using the Benjamini–Hochberg approach (q-value).

We also developed control proximity biotinylation constructs for two purposes: 1) to identify proteins that move alongside PI4P and 2) to differentiate between Nlrp3-specific interactors and changes in subcellular localization. This approach was necessary because APEX2 indiscriminately labels proteins within a ∼20 nm falloff radius [147], potentially leading to false positives when proteins are colocalized without specific interaction, particularly if there are significant alterations in cellular localization. To address this, we linked APEX2 to PI4P biosensors with different affinity for lipids, known as P4C and P4M (Figure 6—Supporting Figures 2A-B) [73][148][149]. Consistent with a previous study [68], only the high-affinity P4C probe maintained visible Golgi localization following fixation with PFA (Figure 6—Supporting Figure 2C). As we encountered difficulties validating the localization of the low-affinity APEX2-P4M construct, we proceeded solely with the high-affinity P4C construct.

Next, we optimized the timing for proximity biotinylation to reflect the initial trafficking of Nlrp3 towards the site(s) where the inflammasome assembles. We determined that many reconstituted cells formed Asc specks within 30 minutes of nigericin stimulation (Figure 6—Supporting Figures 1E), which we established as our final time point. We performed separate time-resolved proximity proteomics experiments, employing either CL097 or nigericin stimulation and using either Nlrp3-APEX2 or APEX2-P4C iBMDMs (Figure 6B). This approach has several important limitations: first, we based our final time point on ASC speck formation, but it is plausible that additional cellular Nlrp3 traffics and activates at later time points but does not recruit Asc due to a sink at the first site of inflammasome assembly. Indeed, NLRP3 oligomers traffic asynchronously within and between cells [150], and thus, snapshots along the activation trajectory reflect a moving population average rather than a consistent pulse. For this reason, we used high concentrations of inflammasome agonists that caused >50% maximum LDH release.

Inactive Nlrp3 molecules likely exchange between cellular compartments, including organelles and the cytosol, which could complicate the interpretation of trafficking patterns associated with activation. We thus initially examined whether the PI4P biosensor interacted with endogenous Nlrp3 in wild-type iBMDMs. We observed that the proximity between Nlrp3 and the PI4P biosensor did not significantly change upon stimulation with inflammasome agonists (Figure 6C) despite detecting numerous other alterations throughout the time course (Figure 6—Supporting Figures 3 and 4). These data are consistent with Nlrp3 fluxing with the PI4P gradient like the P4C biosensor and support its use as a location-specific control.

We then investigated whether inflammasome agonists influenced interactions between P4C or Nlrp3 and particular organellar protein groups as a proxy for organellar localization. Such widespread alterations might signify trafficking or disruptions in cellular architecture. Consistently, treatment with inflammasome agonists caused a rapid decrease in proximity between Nlrp3 or P4C and the ER (Figure 6D). These data suggest that Nlrp3 and P4C traffic away from the ER, and/or that contact sites between the ER and PI4P-rich endomembranes, where Nlrp3 resides, become disrupted. This observation aligns with a recent study, which found that inflammasome agonists disrupt ∼50% of ER-TGN and ER-endosome contact sites within five minutes [58].

To further interrogate temporal changes in the Nlrp3 and P4C proximal proteomes, we clustered ANOVA-significant proteins by their fold changes to the untreated condition across the time course of inflammasome activation. These Leiden clusters, visualized on a UMAP embedding of the data, separate groups of proteins that share similar association kinetics with Nlrp3 or P4C. Several of these kinetic clusters are enriched for different organellar proteomes (Figure 6—Supporting Figures 3-6); in particular, clusters with high labelling early in the time course are enriched for the ER, whereas clusters with high labelling late in the time course are enriched for components of the actin cytoskeleton. These findings align with the loss of ER contact and the presence of endosomal actin comets during Nlrp3 inflammasome activation [58].

We then identified meta-groups of clusters across experiments that contained similar proteins, using the Jaccard index to compare Leiden clusters ([Methods](https://harperlab.pubpub.org/pub/nlrp3/release/1#n4hq1ynwo6t)). The meta-groups with the highest cross-experiment similarity are those representing high labelling either at the beginning or at the end of the respective time course (Figure 6E). The meta cluster representing the beginning of each trajectory (cluster 1) was enriched with lysosomal, mitochondrial, and ER proteins, whereas the end state cluster (cluster 0) was enriched with actin binding, stress granule, and cytosolic proteins (Figure 6E-F). For the Nlrp3 bait, treatment with either compound consistently resolved additional groups of proteins that include endosomal intermediates (Figure 6E-F).

Finally, we created linear models for either nigericin or CL097 stimulation to better understand the differences in responses between Nlrp3 and P4C. To do so, we first combined and normalized our timecourse TMTplexes with MSstats [151], eliminated missing values, and modelled the linear range of the data (0-30 min nigericin or 0-30 min CL097). We plotted the interaction term that characterizes variance in treatment response between the two different APEX baits. We observed comparable rates of Asc proximity change between Nlrp3 and the P4C biosensor, as indicated by their interaction terms (Figure 6G). These data are consistent with Asc filament nucleation and expansion occurring within a PI4P-rich environment[57].

### The Nlrp3 proximity network in response to inflammasome agonists

Our prior APEX2 experiments effectively captured the broad subcellular trafficking events that accompany Nlrp3 inflammasome activation and revealed discernible patterns such as the kinetics of Asc association. However, significant alterations in subcellular localization over the timecourse, and therefore changes to the proximity background, obscure the reliable identification of novel Nlrp3 interactors within these datasets. This problem is further exacerbated by the proximity of P4C to endogenous Nlrp3 (and presumably its interactors). Furthermore, direct comparisons between different TMTplexes are constrained by missing protein/peptide values across datasets. To address these limitations, we performed an additional APEX2 experiment wherein we reconstituted *Nlrp3*^-/-^ iBMDMs with either APEX2-P4C or Nlrp3-APEX2 (Figure 7A). We selected single time points falling within a linear response range for both nigericin and CL097 and subsequently analyzed all 18 replicates with TMT-based quantitative proteomics.

**Figure 7.**
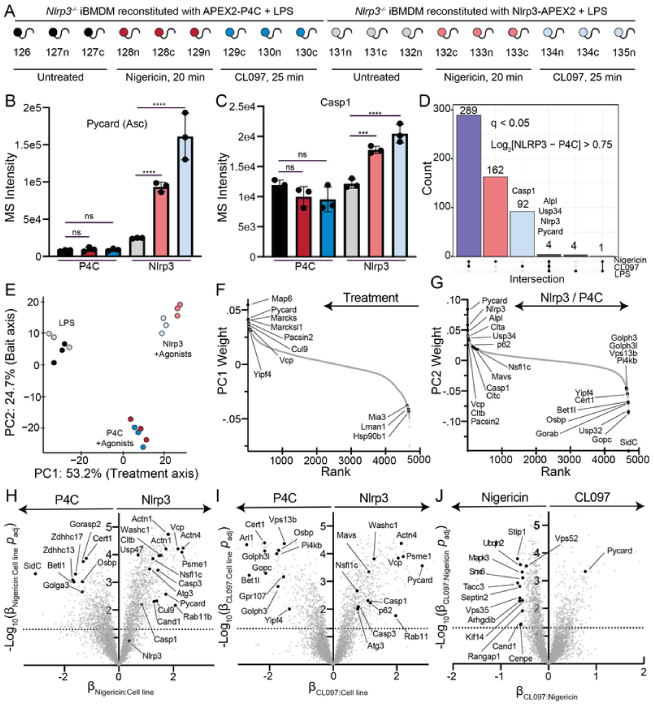
Proximity biotinylation of Nlrp3 and P4C within the same TMTplex. (A) Combined APEX2 experiment design. *Nlrp3^-/-^* iBMDMs expressing either APEX2-P4C (background control) or Nlrp3-APEX2 were primed with LPS (1 μg/mL, 4 h) then treated with nigericin (20 μM, 20 min) or CL097 (75 μg/mL, 25 min) prior to labeling with H_2_O_2_ (10 μM, 1 min). All conditions were also treated with biotin phenol (500 μM, 45 min) prior to labeling. (B-C) Asc and Casp1 proximity to APEX2-P4C does not change over the course of the experiment, whereas it increases with stimulation for Nlrp3-APEX2. (B) Asc and (C) Casp1 MS intensity values, colored as in (A). Error bars represent ± standard deviation of n=4 biological replicates. ns, not significant. (D) UpSet plot of proteins enriched in Nlrp3-APEX2 over APEX2-P4C (any matched comparison of Nlrp3/P4C: q < 0.05, Log_2_FC > 0.75). Intersections show multiple instances of Nlrp3-APEX2 enrichment over the APEX2-P4C background. (E) Principal component analysis (PCA) of the data colored as in (A). PC1 describes treatment with inflammasome agonists, whereas PC2 separates Nlrp3-APEX2 versus APEX2-P4C responses to treatment. (F-G) Ranked contributors to (F) PC1 and (G) PC2. (H-I) Intensity values were fit with generalized linear models. The effect of the treatment (β1), APEX2 bait condition (β2), and their interaction (β1β2) were tested. The difference in slopes represents the β1β2 interaction, which corresponds to differences in the behavior of APEX2 handles (Nlrp3 versus P4C) in response to inflammasome agonists (β1β2 = β_Treatment:Cell line_). Volcano plots of the β1β2 interaction term for (H) nigericin-treatment and (I) CL097-treatment are shown. (J) Intensity values were fit with generalized linear models. The effect of the nigericin treatment (β1), CL097 treatment (β2), and their interaction (β1β2) were tested. The difference in slopes represents the β1β2 interaction, which corresponds to differences in the behavior of Nlrp3 in response to each treatment (β1β2 = β_Nigericin:CL097_). A volcano plot of the β1β2 interaction term is shown. *P* values for (B-C, H-J) were calculated with the Student’s t-test (two sided) and adjusted for multiple hypothesis correction using the Benjamini–Hochberg approach (q-value)

Like previous APEX2 experiments, treating cells with nigericin or CL097 caused a large reduction in the proximity between ER and Nlrp3 or P4C (Figure 7—Supporting Figures 1B-D). However, in contrast to WT iBMDMs reconstituted with APEX2-P4C (Figure 6G), *Nlrp3*^-/-^ iBMDMs reconstituted with APEX2-P4C failed to recruit Asc following nigericin or CL097 stimulation (Figure 7B). These data further support a model where sites of PI4P enrichment serve as specific locations for inflammasome assembly [36][57][58][59] rather than coincidental colocalization following inflammatory cellular stress. Notably, despite a marked increase in proximity to the previously reported mitochondrial interactor Mavs [107], proximity between Nlrp3 and mitochondrial proteins did not surpass the background level observed with P4C in bulk (Figure 7—Supporting Figures 1D-E).

For Nlrp3-APEX2, proximity to Asc and Casp1 increased following stimulation with either agonist, as expected (Figures 7B-C). A recent human NLRP3 interactome study conducted an extensive meta-analysis of additional NLRP3 interactors detected in low and high-throughput studies [152]. From this analysis, we selected several high-confidence interactors validated with biochemical or cellular data. The proximity of several such interactors was higher than the P4C background, including IKKε [153], PP2A [154], Brcc3 [155], and Mavs [107] (Figure 7—Supporting Figure 1E). In total, 552 proteins were proximal to Nlrp3 over background (q value < 0.05, log_2_ fold change > 0.75) in at least one treatment condition (Figure 7D, [Supplementary Data](https://harperlab.pubpub.org/pub/nlrp3/release/1#supplemental-data-resources-and-reagents)).

We subsequently investigated variance between different conditions with PCA and linear modelling. The largest principal component (PC1) accounted for the variance resulting from inflammasome agonist treatment, while the second principal component (PC2) reflected differences between Nlrp3 and P4C background after treatment (Figure 7E). Notably, both bait proteins exhibited minimal differences in the two largest PCs (which collectively explained approximately 80% of the variance between replicates) with LPS stimulation alone. We visualized the specific contributors to PC1 and PC2 (Figures 7F-G). Apart from several known interactors that were dependent on stimulation, such as Mavs, Asc (Pycard), and Casp1, components of the clathrin coat (Clta/b/c) and p97 complexes (Vcp/p97, Nsfl1c) also contributed to this variance. Clathrin coat proteins, actin-related proteins, and p97 complexes were enriched in linear models describing differences in treatment responses between Nlrp3 and P4C (Figures 7H-J).

## Discussion

While certain innate immune pathways directly detect the presence of microbial ligands or their effectors, others embed deep within homeostasis networks to sense broad perturbations to cell state. These sensors integrate diverse cellular cues to respond appropriately to potential threats. Such indirect sensing has advantages and disadvantages, as previously discussed [5]. Notably, numerous pathogens may induce similar host cell states, and these pathogens may struggle to evolve away from eliciting an immunogenic effect while maintaining productive infection. Moreover, the coevolution of innate immune pathways alongside host quality control likely facilitated the exchange of mechanisms and vestigial components between them to enable broad antimicrobial control. However, such coevolution also carries the risk of inflammation arising from cell states induced by sterile damage or stress, contributing to various diseases, from skin disorders to neurodegeneration and ageing. Hence, tight regulation of signal integrators, such as context-dependent thresholding, is crucial for differentiating between sterile programmatic changes in cell state and those induced by existential threats.

The NLRP3 inflammasome, one such signal integrator, accomplishes this through an extensive regulatory network that adjusts the thresholds required for its activation and modulates the intensity of its response. The context dependency for NLRP3 inflammasome signalling is evidenced by the growing number of reported cellular pathways and posttranslational modifications that tune inflammasome signalling [24][25][156]. Additionally, many variations in signalling have been observed among commonly used experimental systems, including BMDMs, THP-1s, BLaER1 macrophages, iMacs, microglia, and primary human macrophages. Consequently, while a unifying stressor may ultimately determine an NLRP3 inflammasome response, the circumstances under which pyroptosis occurs in response to a given stimulus depend on cell type and state.

Discovering context-dependent NLRP3 interactors is likely important for modulating the inflammasome response in specific diseases, warranting further investigation. In this study, we employed APEX2-based proximity labelling in iBMDMs to explore a subset of these interactors. The proximal proteome reveals trafficking of Nlrp3 away from ER contact towards actin-binding and endosome-associated proteins, implicating these organelles in the assembly of the inflammasome complex. This observation aligns with previous studies suggesting the involvement of endosomes in inflammasome activation [44][58][59][74] but has the potential to add granularity to our understanding of the molecular events driving this process. Several important questions remain—including the precise mechanism by which PI4P-rich endosomes recruit NLRP3, whether other organellar perturbations that accumulate acidic lipids activate or potentiate NLRP3 [157], how these organellar sites facilitate the necessary conformational changes in NLRP3 to initiate inflammasome assembly, and how different NLRP3 trafficking routes are regulated [158].

Contextualizing how different perturbations threshold NLRP3 inflammasome assembly requires characterizing the minimally sufficient series of cellular events triggered by inflammasome agonists and how NLRP3 ultimately integrates them. Recent studies have highlighted the loss of ER-endosome contact sites and disrupted endosomal trafficking as common perturbations induced by various inflammasome activating stressors [44][58][59]. We sought to investigate how nigericin and CL097, compounds with different mechanisms of action, induce distinct cellular states that overlap in their ability to activate NLRP3. Specifically, we characterized how these agonists impact the composition of organelles through spatial proteomics of mitochondria, lysosomes, EEA1-positive endosomes, and the Golgi following NLRP3 agonist treatment. Our findings reveal numerous alterations to cell state, many of which have been observed since the discovery of pathological mutations in NLRP3 [159] and the coining of the term “inflammasome” [6] over two decades ago. Notably, our data suggest that inflammasome agonists affect distinct retrograde trafficking pathways, such as COPI and particular endosome-TGN transport routes involved in Shiga toxin trafficking.

We speculate that numerous yet-to-be-discovered genetic, ageing, and lifestyle-related factors affect a cell’s capacity to mount an NLRP3 inflammasome response in sterile contexts, thereby contributing to disease pathogenesis. This model is supported by several observations: many small molecules or stressors, though insufficient for NLRP3 activation on their own, can either enhance or suppress NLRP3 responses [59][160] [161][162]; NLRP3 mutations cause a variety of clinical manifestations in particular tissue [27][28][163]; and several studies have linked NLRP3 activation to age-related decline [164][165][166][167]. Our spatial proteomics data serve as a cellular reference to characterize novel perturbations affecting NLRP3 activation as they emerge.

## Negative results and rationale

### TGN38 disruption does not require NLRP3 expression, and iBMDMs were not trackable for organelle-IPs

We stably introduced TMEM115-3xHA (the GolgiTag) or 3xFLAG-EEA1 (the EndoTag) into immortalized mouse bone marrow-derived macrophages (iBMDMs) with lentivirus and sorted for low expressing cells with FACS. We found that TMEM115 colocalized with TGN38 (the rodent homolog of TG46), a protein that cycles between the plasma membrane, endosomes, and the Golgi apparatus [168][169] in untreated cells (Negative Data Figure 1). 3xFLAG-EEA1 localized to endosomes, albeit inconsistently (Negative Data Figure 2). TMEM115 and EEA1 retained their localization after treatment with the potassium efflux-dependent inflammasome agonist nigericin, whereas TGN38 dispersed in a manner that did not depend on NLRP3 expression (Negative Data Figures 1-2). Similarly, TMEM115 and EEA1 retained their localization following treatment with CL097, a potassium efflux-independent agonist that caused only modest TGN38 redistribution (Negative Data Figures 1-2).

**Negative Data Figure 1.**
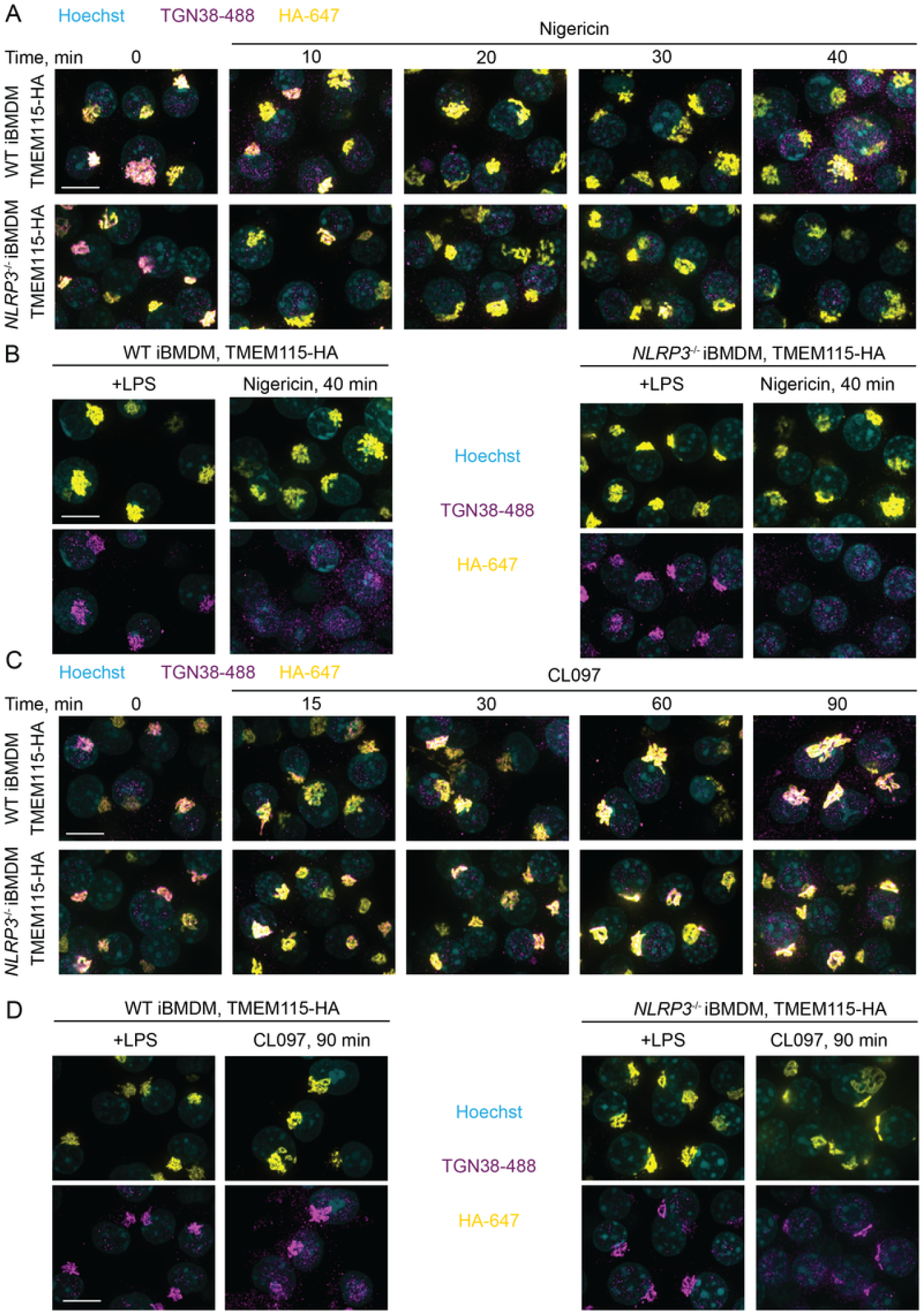
NLRP3 agonists do not disrupt TMEM115 (GolgiTag) localization in iBMDMs, and TGN38 dispersion does not depend on Nlrp3 expression. (A-D) Control and *Nlrp3^-/-^* iBMDMs reconstituted with TMEM115-3xHA were LPS primed (1 μg/mL, 4 h), treated with 20 μM Nigericin (A and B) or 75 μg/mL CL097 (C and D) for the indicated amount of time, fixed, immunostained, and imaged by spinning disk confocal microscopy. Maximum intensity projection images (z=8 μM, 29 steps), representative of n > 3 fields of view. Scalebar = 10 μM.

**Negative Data Figure 2.**
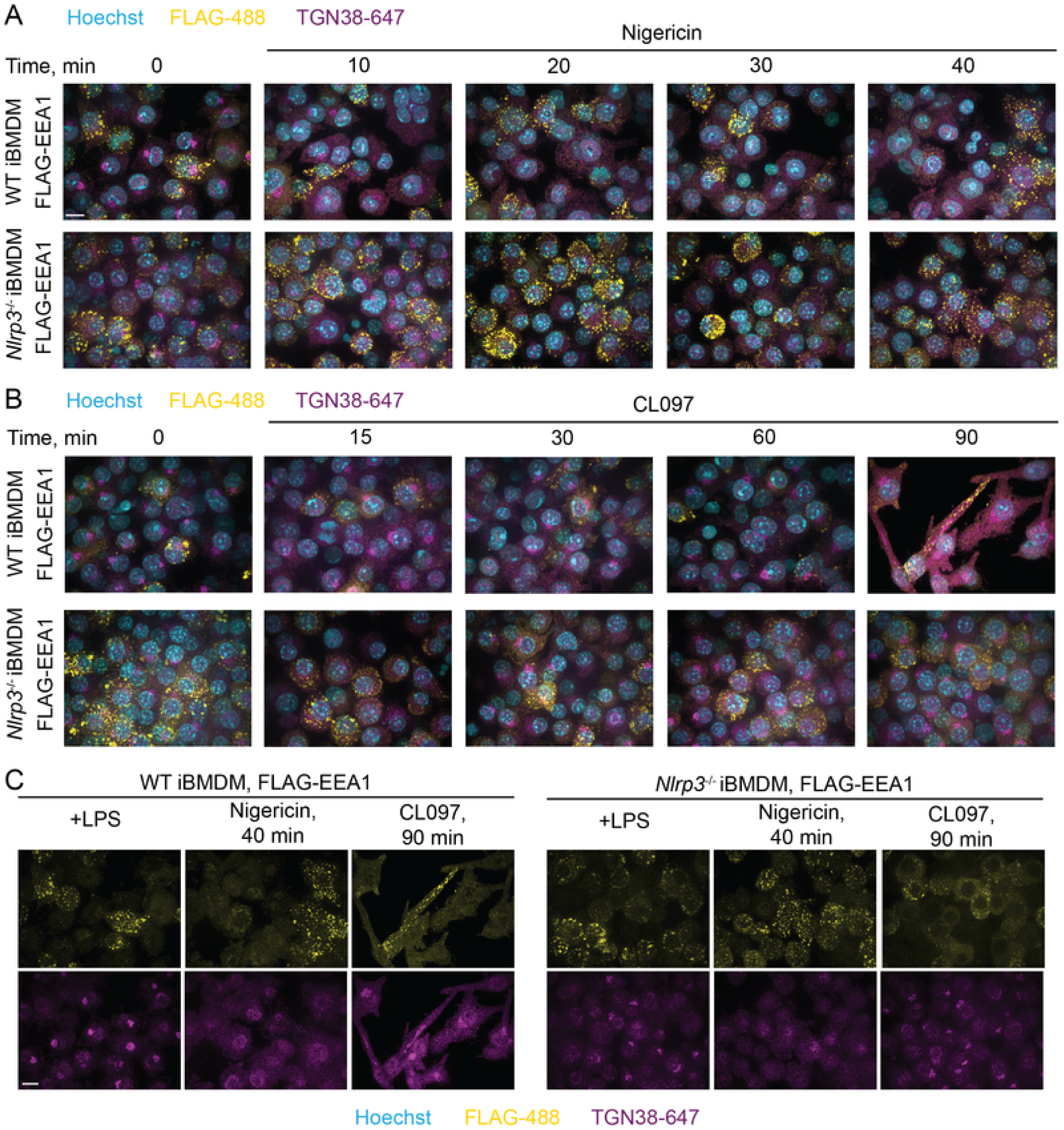
NLRP3 agonists do not disrupt EEA1 (EndoiTag) localization in iBMDMs, and TGN38 dispersion does not depend on Nlrp3 expression. (A-D) Control and *Nlrp3^-/-^* iBMDMs reconstituted with 3xFLAG-EEA1 were LPS primed (1 μg/mL, 4 h), treated with 20 μM Nigericin (A and C) or 75 μg/mL CL097 (B and C) for the indicated amount of time, fixed, immunostained, and imaged by spinning disk confocal microscopy. Maximum projection images are shown. Maximum intensity projection images (z=8 μM, 29 steps), representative of n>3 fields of view. Scalebar = 10 μM.

These data agree with our EndoIP and GolgiIP results in HEK293T cells (Figures 4 and 5) and with previous studies [57][58][59]. We cannot exclude the possibility that the presence of NLRP3 slightly exacerbates the endosomal TGN38/46 phenotype, which could occur because of aggregate-based Golgi or endolysosomal quality control pathways [170]. However, our data support a model where TGN38/46 dispersion into endosomes is a consequence of cellular stress rather than a feature of NLRP3 activation, in agreement with recent studies [58][59]. Further supporting this conclusion, a phenotypic CRISPR screen demonstrated that many cellular perturbations of transport likely unrelated to NLRP3 activation induce TGN38/46 peripheral redistribution [171].

We were, however, unsuccessful in isolating organelles using immunoprecipitation (organelle-IP) from these iBMDMs. This failure was due to our inability to achieve a gentle enough lysis to maintain organelle integrity, possibly because of the round morphology of the iBMDMs. For this reason, we pivoted to the toolkit of HEK293(T) cells that we developed for organelle-IPs rather than a more physiological system. HEK293(T) cells can undergo NLRP3 inflammasome responses when reconstituted with pathway components, and many trafficking proteins are conserved between different cell types. Thus, the relevant cellular stress induced by activating compounds (e.g., nigericin and CL097) are likely similar. While performing these experiments, we developed a method to isolate organelles from PMA-differentiated THP-1 macrophages by organelle-IP ([see methods](https://harperlab.pubpub.org/pub/nlrp3/release/1#nc20foiciwt)). The extended morphology of differentiated THP-1s likely provides more adequate surface area for gentle lysis by Dounce homogenization, thus facilitating organelle-IP approaches.

### NLRP3 activators accumulate HOOK2 and FHIP2A at the Golgi, but they do not modulate NLRP3 activity

Mechanisms of NLRP3 trafficking remain controversial. Some reports demonstrate that oligomerized NLRP3 undergoes dynein-mediated transport to the microtubule organizing center (MTOC) [158] in a manner that requires the aggresome adaptor HDAC6 [172]. However, high-resolution imaging techniques such as cryo-electron tomograms or super-resolution microscopy failed to detect the inflammasome at the MTOC [74][150], and the application of an HDAC6 degrader molecule did not affect inflammasome formation in most cellular contexts [173]. Additionally, compounds that induce potassium efflux or specific deletions in NLRP3 protein can traffic independently of the MTOC [158]. Thus, we checked whether dynein or dynein activating adaptors were spatially disrupted by either of the NLRP3 agonists.

We noticed that HOOK2 and FHIP2A (FAM160B1), but not other dynein activating adaptors, were highly enriched on the Golgi upon nigericin, CL097, and monensin treatment (Negative Data Figure 3A-D). The FHF complex comprises Hook, FHIP (FHF complex subunit Hook Interacting Protein), and AKTIP/FTS [174] [175]. Mammals encode three Hook proteins and four FHIPs, and different FHF compositions associate with different cargo [175][176]. HOOK2/FHIP2A-containing complexes associate with Rab1A-tagged ER-to-Golgi cargo [175]. HOOK2 also promotes aggresome formation [177] and, therefore, might threshold inflammasome activation.

**Negative Data Figure 3.**
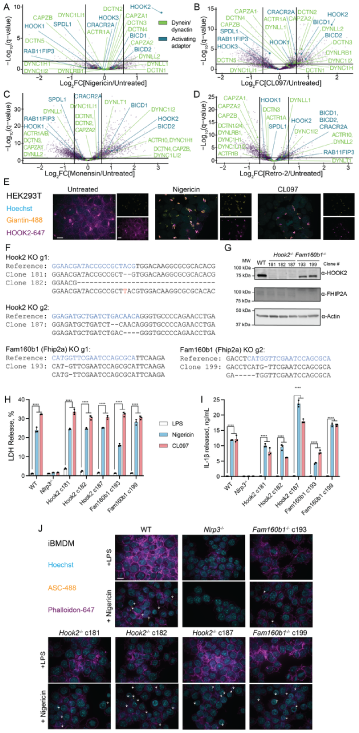
Inflammasome agonists cause Golgi retention of Hook2, but Hook2 and Fhip2a knockout does not affect inflammasome signalling in iBMDMs. (A-D) GolgiP volcano plots for (A) nigericin (20 μM, 30 min)-treated, (B) CL097 (75 μg/mL, 1 h)-treated, monensin (10 μM, 2 h)-treated, or (D) retro-2 (25 μM, 2 h)-treated versus untreated HEK293T^EG^ cells. Dynein/dynactin and dynein activating adaptors are colored as shown. *P* values were calculated from the Student’s t-test (two sided) and adjusted for multiple hypothesis correction using the Benjamini– Hochberg approach (q-value) with MSstats. Data represent n=3 or 4 biological replicates. For the full TMTplex experimental setup, see Figure 5—Supporting Figures 1 and 4. (E) Inflammasome agonists collapse peripheral/cytosolic HOOK2 staining to the Golgi. HEK293T cells were treated with nigericin (20 μM, 30 min) or CL097 (75 μg/mL, 1 hr), fixed, and immunostained for HOOK2 and giantin (Golgi marker). Maximum intensity projections (8 μM over 29 steps) are representative of 2 independent experiments each with n > 7 fields of view. Scale bar = 10 μm. (F) Miseq genomic DNA results from Hook2 and Fhip2a (Fam160b1) knockout iBMDM clones. The gRNA reference sequence is colored in blue, gaps are indicated with dashes, and insertions are indicated with red letters. (G) Western blotting demonstrates that genomic knockout clones do not express either Hook2 or Fhip2a. (H-I) Hook2 and Fhip2a knockout does not alter (H) LDH or (I) IL-1β release from iBMDMs. Cells were treated with LPS (1 μg/mL, 4 hr) prior to the addition of nigericin (20 μM, 1 hr) or CL097 (75 μg/mL, 2 hr). Bar graphs represent mean ± standard deviation, n=3 biological replicates. ****p<0.0001 from a two-way ANOVA with Tukey’s post-test. (J) Hook2 and Fhip2a knockout does not alter ASC speck formation. The indicated primed (4 h LPS, 1 μg/mL) iBMDMs were treated with nigericin (0 or 20 μM, 30 min), fixed, immunostained, and imaged. Arrow heads point to ASC specks. Scale bar = 10 microns. Representative maximum intensity projection images (8 μM over 29 steps) from n > 7 fields of view.

We first validated that nigericin and CL097 collapsed peripheral HOOK2 staining to the Golgi region in HEK293T cells (Negative Data Figure 3E). To test whether a HOOK2/FHIP2A complex modulates the NLRP3 inflammasome response, we generated clonal knockouts of Fhip2A or Hook2 in iBMDMs (Negative Data Figure 3F-G). These FHS knockouts did not perturb NLRP3 inflammasome signalling in response to LPS alone, LPS+nigericin, and LPS+CL097 treatment, as measured by the release of LDH or bioactive IL-1β into the cell culture supernatant, or the formation of ASC specks (Negative Data Figure 3H-J). Thus, different classes of NLRP3 inflammasome agonists accumulate Golgi-localized HOOK2 and FHIP2A, but their loss does not directly impact inflammasome activation in iBMDMs. Golgi accumulation of HOOK2 and FHIP2A might instead coincide with Golgi alkalinization or specific disruptions to retrograde trafficking, as they were not affected by retro-2 (Negative Data Figures 3A-D). However, this phenomenon warrants further study.

### Several contact site, lipid transfer, and Shiga toxin trafficking protein knockouts did not impact inflammasome activation

Inflammasome agonist treatment depleted several proteins associated with lipid transfer and organellar contact sites from the Golgi or shifted them between Golgi compartments that were differentially enriched by TMEM115-mediated GolgiIPs (Figure 5A-D). PLEKHA3 (FAPP1) knockdown increases Golgi PI4P without disrupting ER-TGN contact sites [68][178]. The proposed function of PLEKHA3 is to act as a bridge between ER-localized Sac1 and Golgi-localized PI4P, allowing Sac1 to dephosphorylate PI4P in *trans*. In contrast, PLEKHA8 (FAPP2) functions in membrane trafficking and lipid transfer [179].

We generated clonal Plekha3 and Plekha8 knockouts in iBMDMs (Negative Data Figures 4A-C). Neither single knockout potentiated nor damped the NLRP3 inflammasome response measured by the release of LDH or IL-1β (Negative Data Figures 4D-E). Additionally, bulk knockout of PLEKHA3 or PLEKHA8 in THP-1 macrophages did not potentiate or dampen NLRP3 signalling (Negative Data Figures 4F-G). These data suggest that Golgi PI4P accumulation alone does not affect inflammasome signalling, which agrees with recent findings that OSBPL10 (ORP10) knockout was insufficient for inflammasome activation [58].

**Negative Data Figure 4.**
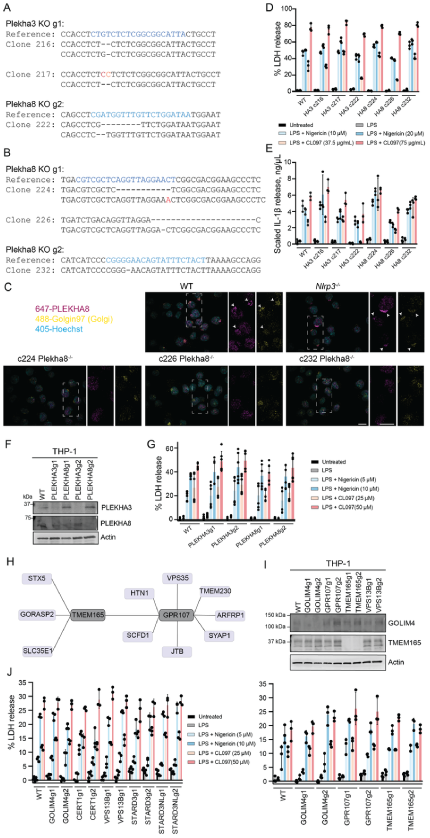
PLEKHA3 or PLEKHA8 knockout is not sufficient to affect NLRP3 inflammasome activation. (A-B) Miseq genomic DNA reads for Plekha3 and Plekha8 knockout iBMDM clones. The guide RNA sequences is colored blue, indels are colored red, SNPs are colored orange, and deletions are indicated with dashed lines. (C) Maximum intensity projections show lack of Plekha8 signal in genetic knockout clones. The indicated iBMDMs were, fixed, immunostained, and imaged. Arrows indicate colocalized Plekha8 and Golgin-97 (Golgi marker) signal in WT and *Nlrp3^-/-^* iBMDMs. Maximum intensity projections (8 μM over 29 steps) are representative of n > 5 fields of view. Scale bar = 10 μm (bottom right panel). (D-E) Plekha3 (HA3) or Plekha8 (HA8) clonal knockout does not affect Nlrp3 activation in iBMDMs. Cells were primed with ± LPS (4 h, 1 μg/mL) and then treated with the indicated concentration of compound (nigericin, 1 h; CL097, 2 h) prior to harvesting supernatant to assess (D) LDH release or (E) IL-1β release. IL-1β levels were normalized for cell number between genotypes with a paired maximum lysis control. Error bars represent ± standard deviation of n=3 biological replicates. (F) Western blot validation of bulk knockouts with two different gRNAs in THP-1 cells. (G) PLEKHA3 or PLEKHA8 bulk knockout does not affect NLRP3 activation in THP-1 cells. Cells were primed with ± LPS (3 h, 1 μg/mL) and then treated with the indicated concentration of compound (nigericin, 1 h; CL097, 2 h) prior to harvesting supernatant for LDH release assays. Error bars represent ± standard deviation of n=3 biological replicates. (H) OpenCell interactome for TMEM165 and GPR107. Many proteins involved in Shiga/ricin toxin transport interact with one another. (I) Immunoblots of the indicated bulk knockouts in THP-1 cells. (J) THP-1 bulk knockouts do not alter LDH release. Cells were treated with LPS (1 μg/mL, 3 h) prior to the addition of nigericin (1 h) or CL097 (2 h), as indicated. Bar graphs represent mean ± standard deviation, n=3 biological replicates.

We tested an additional suite of lipid transfer and organellar contact site proteins that were either depleted in our GolgiIP data (Figure 5) or enriched within PI4P-rich sites biotinylated by the P4C-APEX2 biosensor (Figures 6-7), including CERT1, VPS13B, STARD3, and STARD3NL. THP-1 cells expressing knockout guides did not dampen or potentiate inflammasome formation (Negative Data Figure 4J), but we could not validate protein-level knockout efficiency with the antibodies available in our lab. Therefore, we caution against overinterpreting these results and acknowledge that single knockouts might not be sufficient cellular stressors to cause an NLRP3-related phenotype.

We also noticed that proteins involved in Shiga toxin trafficking were enriched on endosomes following CL097 and nigericin treatment (Figures 4A-D). One of these proteins, GOLIM4, cycles between the *cis*-Golgi and endosomes[122][123]. Retro-2, a compound that potentiates NLRP3 activation [59], interacts with the ER exit site protein SEC16A, thereby blocking the downstream interaction between STX5 and GOLIM4 required for retrograde trafficking of Shiga toxins and ricin [124][140]. Endolysosomal pH disruption by monensin, which also potentiates inflammasome signalling [59], causes GOLIM4 accumulation in endosomes [123]. Manganese, which activates NLRP3 in microglia [127], also blocks Shiga toxin transport by rapidly dispersing GOLIM4 to endolysosomes [121][125]. We wondered whether there was a direct connection between disrupting Shiga/ricin toxin transport pathways and inflammasome activation.

First, we identified potential interactions among several Shiga/ricin toxin transport proteins within the OpenCell database [180](Negative Data Figure 4H). Furthermore, human mutations in TMEM165, one of these proteins, have been associated with “unexplained fever episodes” that coincide with NLRP3 gain-of-function mutations [28][181]. We knocked out either GOLIM4, GPR107, or TMEM165, but none of these single perturbations caused spurious inflammasome signalling in THP-1 cells (Negative Data Figure 4J). We cannot, however, rule out that these knockouts affect NLRP3 thresholding in other cell lines, such as primary human macrophages/microglia or transdifferentiated BLaER1 macrophages. Additionally, redundancy within the pathway might prevent the sufficiency of any single genetic perturbation in these cell lines. Since TMEM165 is involved in Golgi pH regulation, it is tempting to speculate that it might regulate the threshold of STING-related NLRP3 activation in particular cell types.

### APEX2-based proximity biotinylation does not reveal new LPS-dependent and LPS-independent NLRP3 interactors

Modifications that regulate Nlrp3 inflammasome signalling may occur under steady-state conditions or depend on transcriptional priming (Negative Data Figure 5A). To discern between these potential interactors, we employed APEX2-based proximity proteomics, utilizing our APEX2-P4C control ([as detailed in the main text](https://harperlab.pubpub.org/pub/nlrp3/release/1#nixv6bpg1mo)). Interestingly, we observed that cells treated with LPS exhibited increased biotinylation of proteins compared to untreated cells (Negative Data Figure 5B). This phenomenon could occur because LPS treatment increases cellular ROS [182][183], which might reduce the buffering capacity for free radicals and thus increase APEX2 enzymatic efficiency. Subsequently, proximity biotinylation experiments revealed the known interactor Pycard (Asc) but few other significant proteins (Negative Data Figures 5C-G). Since enzymes that post-translationally modify Nlrp3 might be short-lived, methods with extended labelling times, such as TurboID [184], might better capture these proximal proteome differences. Nonetheless, these findings serve as a resource for uncovering proteins proximal to PI4P, as the APEX2-P4C condition biotinylated many proteins (Negative Data Figure 5D-F).

**Negative Data Figure 5.**
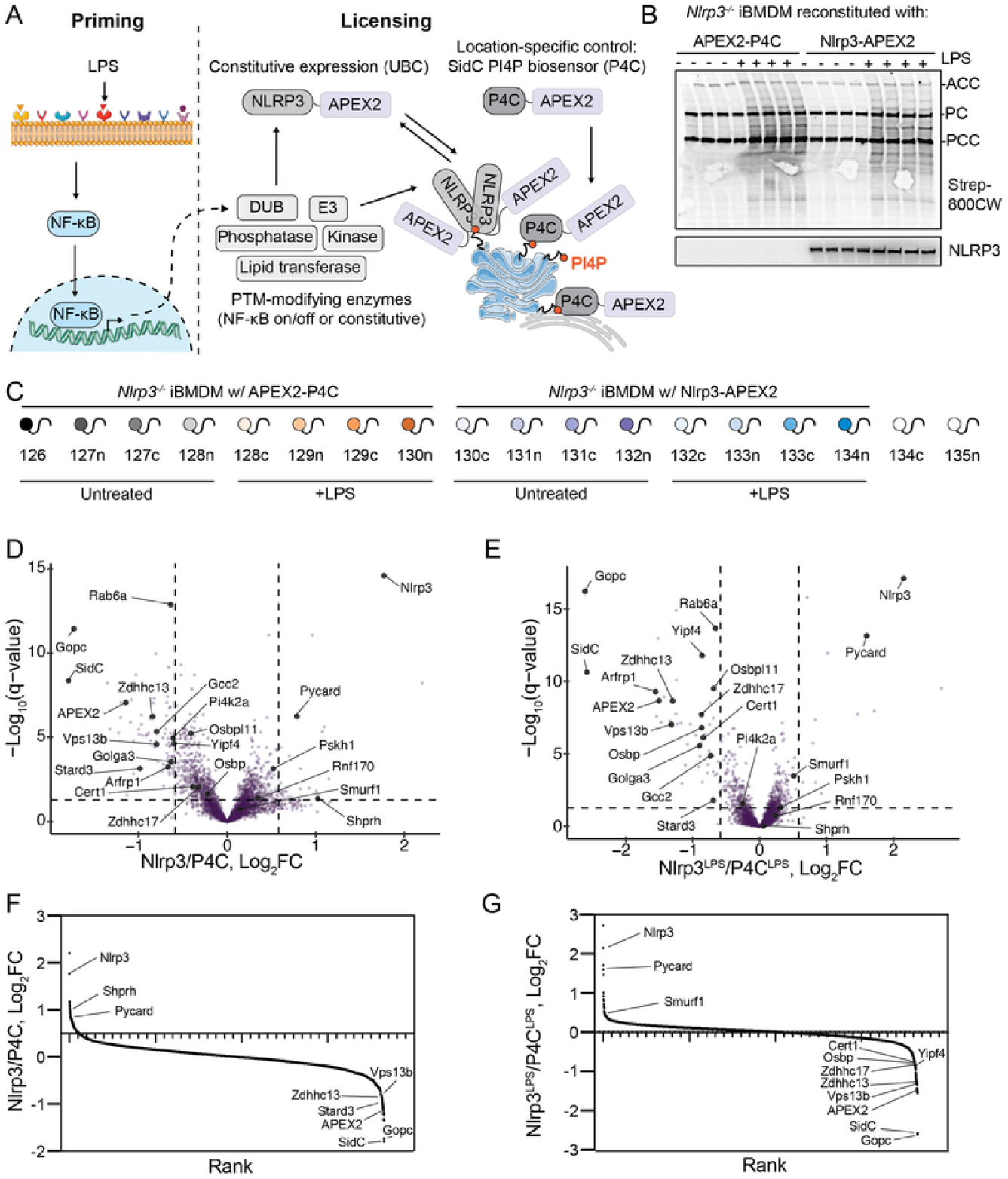
Proximity biotinylation of Nlrp3 and P4C ± LPS. (A) Schematic of APEX2-based proximity biotinylation during priming. While the lentivirally-reconstituted APEX2 fusions are constitutively expressed, LPS leads to proteome changes that could include enzymes that modify Nlrp3. Nearby proteins are indiscriminately biotinylated within a ∼20 nm falloff radius by APEX2. The P4C biosensor binds to PI4P and thus serves as a location-specific control. (B) APEX2 input immunoblots with the indicated antibodies/conjugates. LPS treatment (1 μg/mL, 4 h) appears to enhance biotinylation efficiency. (C) TMTplex experimental design. The indicated iBMDM cells were treated with or without LPS (1 μg/mL, 4 h) prior to labeling with H_2_O_2_ (10 μM, 1 min). All conditions were also treated with biotin phenol (500 μM, 45 min) prior to labeling. (D-E) APEX2 volcano plots for (D) untreated or (E) LPS (1 μg/mL, 4 h)-treated comparisons between Nlrp3 and P4C cell lines. *P* values were calculated from the Student’s t-test (two sided) and adjusted for multiple hypothesis correction using the Benjamini–Hochberg approach (q-value) with MSstats. Data represent n=4 biological replicates.

## Supporting figures

Figure 2 **supporting figure**

**Figure 2—Supporting Figure 1.**
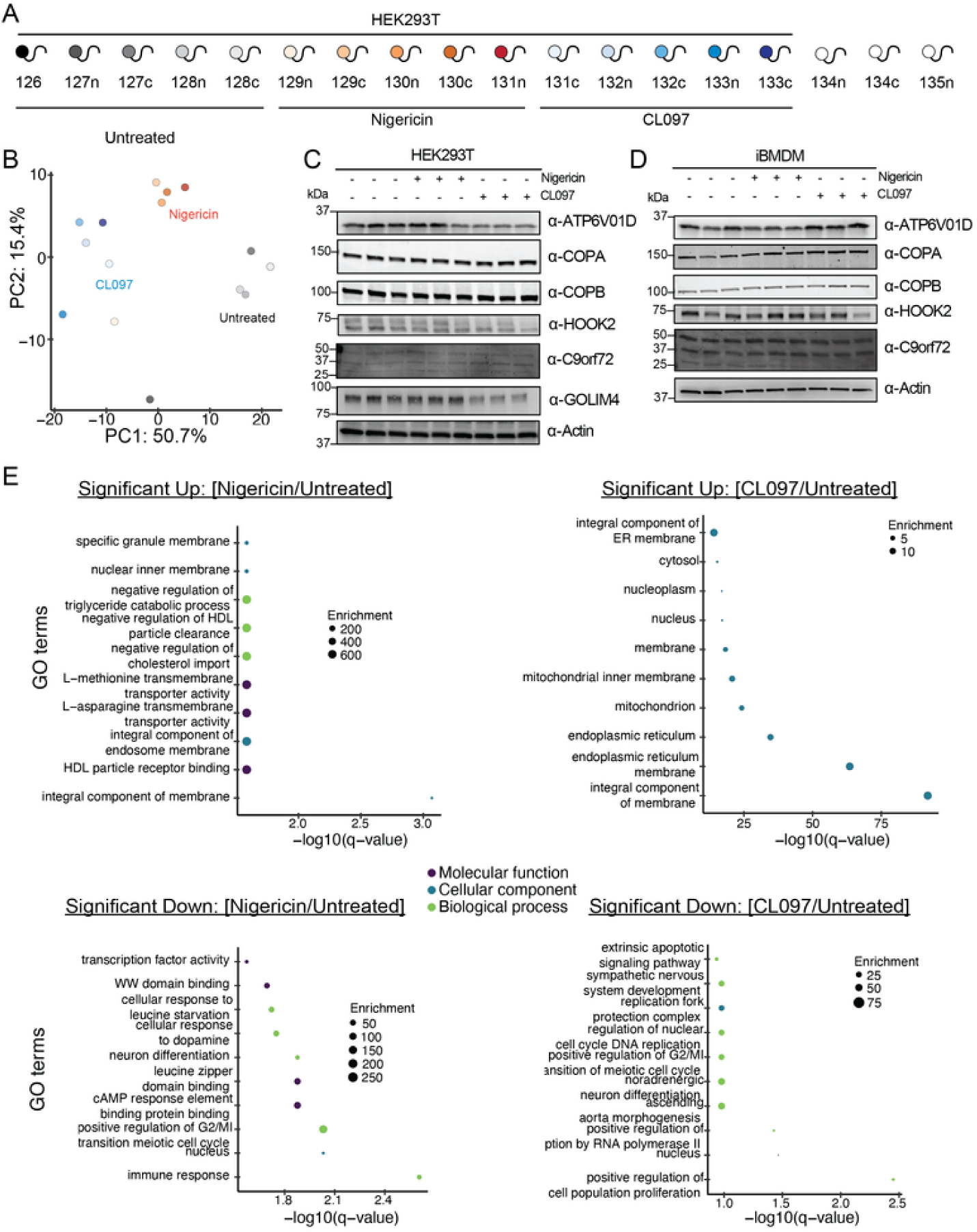
TMT design, GO term analysis, and other supporting data for the whole cell proteomics experiment. (A) TMTplex experimental design. Cells were treated with the indicated compounds (nigericin: 20 μM, 30 min; CL097: 75 μg/mL, 1 h) prior to harvesting lysates for whole cell proteomics. n=5 biological replicates per condition. (B) Principal component analysis (PCA) of the data colored as in (A). (C-D) Western blots from (C) HEK293T cells and (D) iBMDMs treated with nigericin (20 μM, 30 min) or CL097 (75 μg/mL, 1 h). n=3 biological replicates shown. (E) GO-term analysis of proteins significantly changing (q > 0.05, Log_2_FC > |0.5|) in response to the indicated treatment. To Figure 2: https://harperlab.pubpub.org/pub/nlrp3#n7w50fsfetx

Figure 3 **supporting figures**

**Figure 3—Supporting Figure 1.**
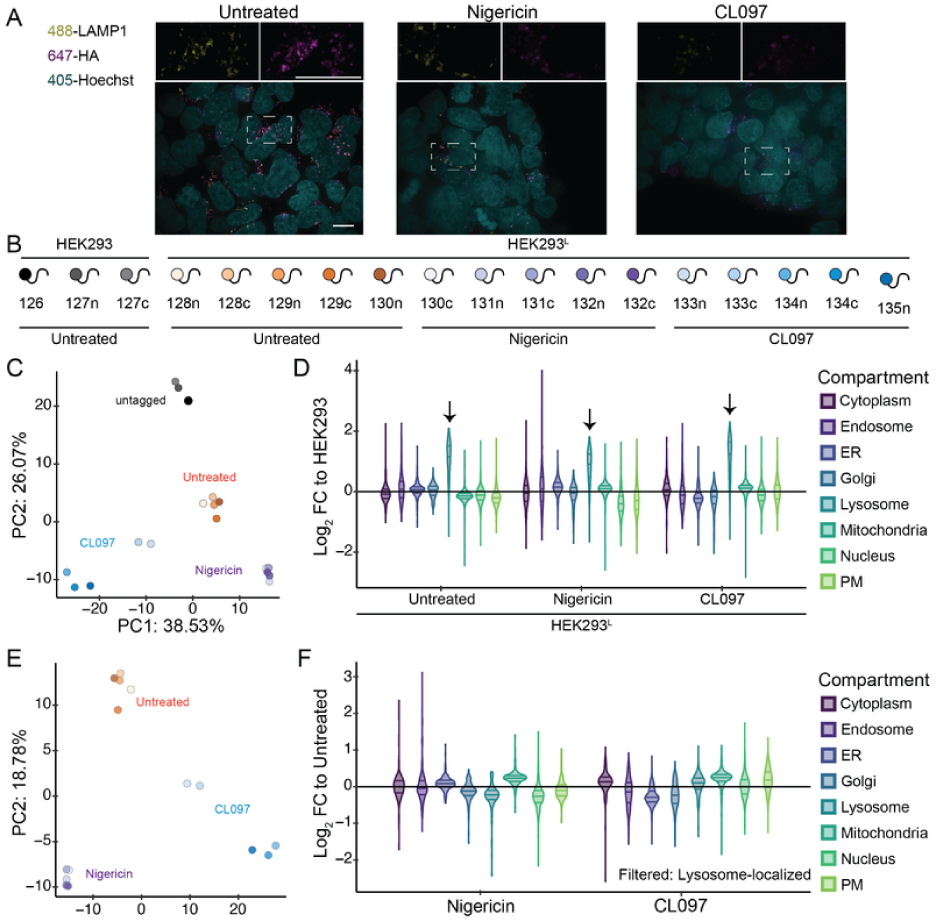
Experimental validation and design for the LysoIP experiment. (A) The LysoTag (TMEM192-HA) colocalizes with the lysosomal marker LAMP1 and maintains colocalization following treatment. HEK293^L^ cells were treated with nigericin (20 μM, 30 min) or CL097 (75 μg/mL, 1 h), fixed, and immunostained with the indicated antibody and Hoechst. Maximum intensity projection images (z=8 μM, 29 steps), representative of n > 6 fields of view. Scale bar (left panel), 10 μm. (B) TMTplex experimental design. Cells were treated with the indicated compounds (nigericin, 20 μM, 30 min; CL097, 75 μg/mL, 1 h) prior to LysoIP on anti-HA magnetic beads. The parental untagged HEK293 cell line serves as a background control, whereas the endogenously tagged HEK293 cell line (HEK293^L^) facilitates LysoIP. (C) Principal component analysis (PCA) colored as in (B) with all channels included. Replicates of a given condition correlate well. (D) Violin plots depicting the log_2_ fold change (FC) values for the indicated subcellular compartment in aggregate for the indicated LysoIP (HEK293^L^) compared to the background control (HEK293). Lysosomal proteins are enriched over all organellar groups for each condition, validating the approach. (E) PCA as colored as in (B) with background samples excluded. Replicates for a given condition cluster well. (F) Violin plots depicting the log_2_ FC values for the indicated subcellular compartment and given treatment condition (nigericin or CL097) compared to untreated (LysoTag) cells. Proteins were first filtered for significant lysosomal localization (any 293L/293: q < 0.05, Log_2_ FC > 0.5) To Figure 3: https://harperlab.pubpub.org/pub/nlrp3#nyjwc0zexqn

**Figure 3—Supporting Figure 2.**
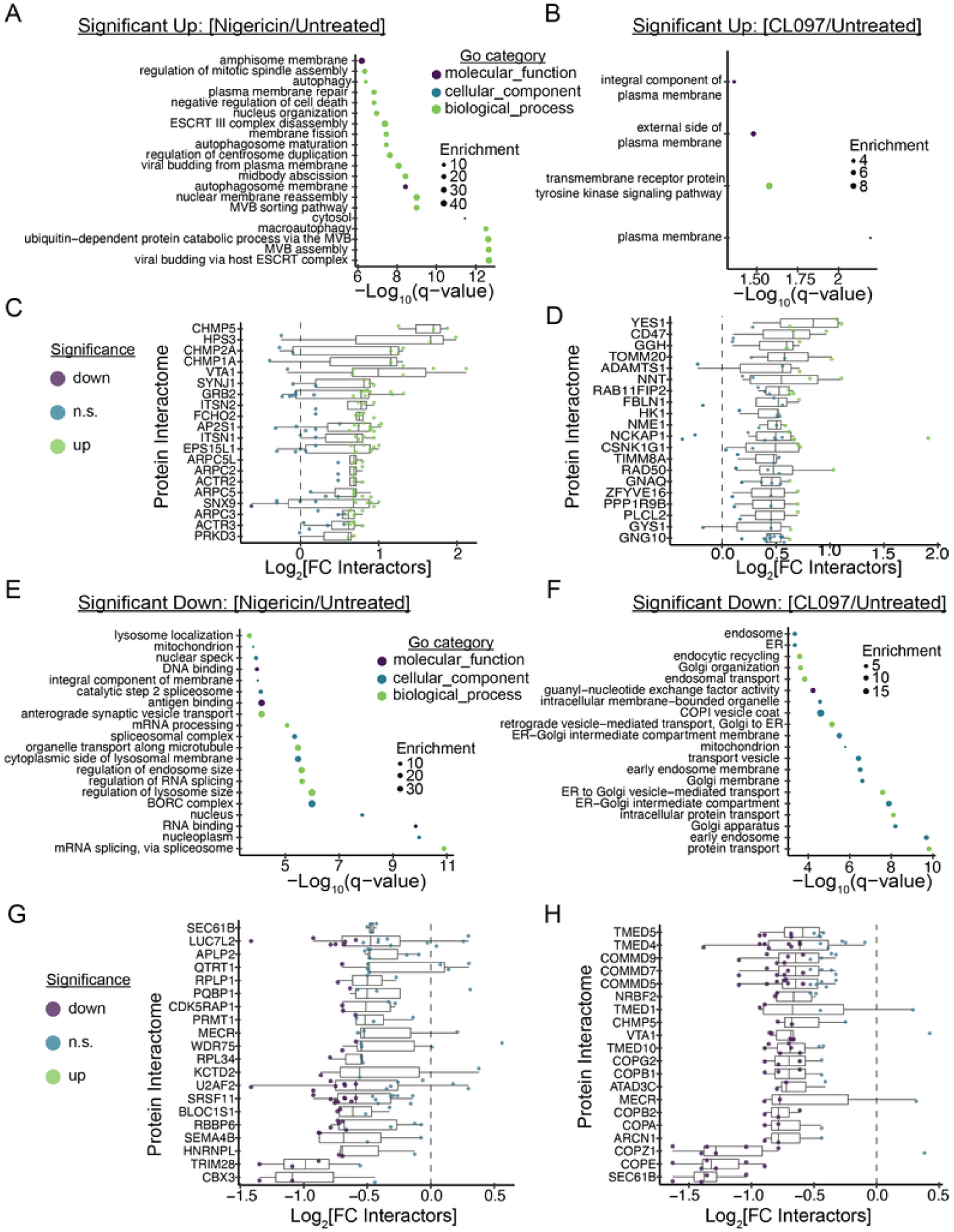
Protein group analysis for the LysoIP experiment. (A-B) GO-term analysis for proteins that significantly increased in the indicated LysoIP versus untreated cells (q < 0.05, Log_2_FC > 0.5). (C-D) Analysis of the top 20 Bioplex protein interactomes (>3 proteins detected) positively enriched in the indicated experiment. (E-F) GO-term analysis for proteins that significantly decreased in the indicated LysoIP versus untreated cells (q > 0.05, Log_2_ FC < −0.5). (G-H) Analysis of the top 20 Bioplex protein interactomes (>3 proteins detected) depleted in the indicated experiment. To Figure 3: https://harperlab.pubpub.org/pub/nlrp3#nyjwc0zexqn

**Figure 3—Supporting Figure 3.**
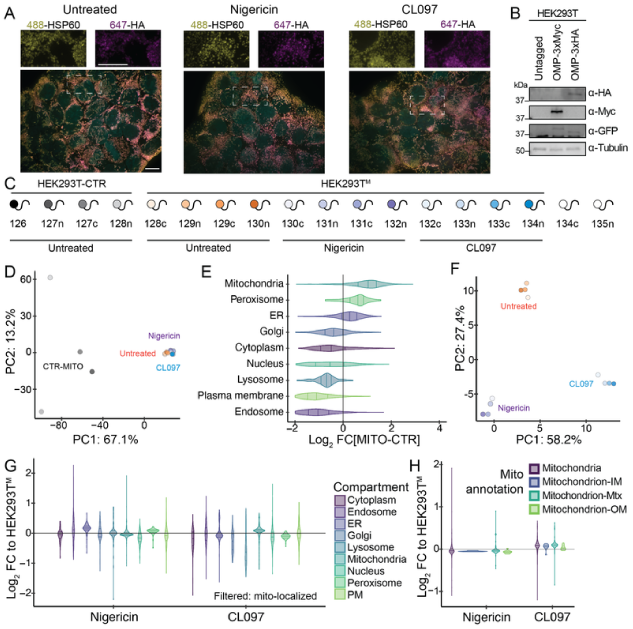
Experimental validation and design for the MitoIP experiment. (A) The MitoTag (OMP25-eGFP-3xHA) colocalizes with the mitochondrial marker HSP60 and maintains colocalization following treatment. HEK293T^M^ cells were treated with nigericin (20 μM, 30 min) or CL097 (75 μg/mL, 1 h), fixed, and immunostained with the indicated antibody and Hoechst. Maximum intensity projection images (z=8 μM, 29 steps), representative of n > 6 fields of view. Scale bar (left panel), 10 μm. (B) TMTplex experimental design. Cells were treated with the indicated compounds (nigericin, 20 μM, 30 min; CL097, 75μg/mL, 1 h) prior to MitoIP on anti-HA magnetic beads. 3xMyc-tagged HEK293T cells serve as the background control (HEK293T-CTR), whereas the 3xHA-tagged HEK293T cell line (HEK293T^M^) facilitates MitoIP. (C) Western blots of the HEK293T-CTR (myc- and eGFP-tagged) and HEK293T^M^ (HA- and eGFP-tagged) cell lines confirming protein expression at similar levels. (D) Principal component analysis (PCA) colored as in (C) with all channels included. Background control IPs separate well from MitoIPs. (E) Violin plots depicting the log_2_ fold change (FC) values for the indicated subcellular compartment in aggregate for the untreated MitoIP (HEK293T^M^) compared to the background control (HEK293T-CTR). Mitochondrial proteins are enriched over all organellar groups for each condition, validating the approach. (F) PCA colored as in (C) with background samples excluded. Replicates of a given condition cluster well. (G) Violin plots depicting the log_2_FC values for the indicated subcellular compartment and the given treatment condition (nigericin or CL097) compared to untreated (MitoTag) cells. Proteins were first filtered for significant mitochondrial localization (any 293T^M^/293T^Control^: q < 0.05, Log_2_ FC > 0.5). (H) Violin plots depicting the log_2_FC values for the indicated mitochondrial annotation and the given treatment condition compared to untreated (MitoTag) cells. Proteins were first filtered for significant mitochondrial localization (any 293T^M^/293T^Control^: q < 0.05, Log_2_FC > 0.5). To Figure 3: https://harperlab.pubpub.org/pub/nlrp3#nyjwc0zexqn

**Figure 3—Supporting Figure 4.**
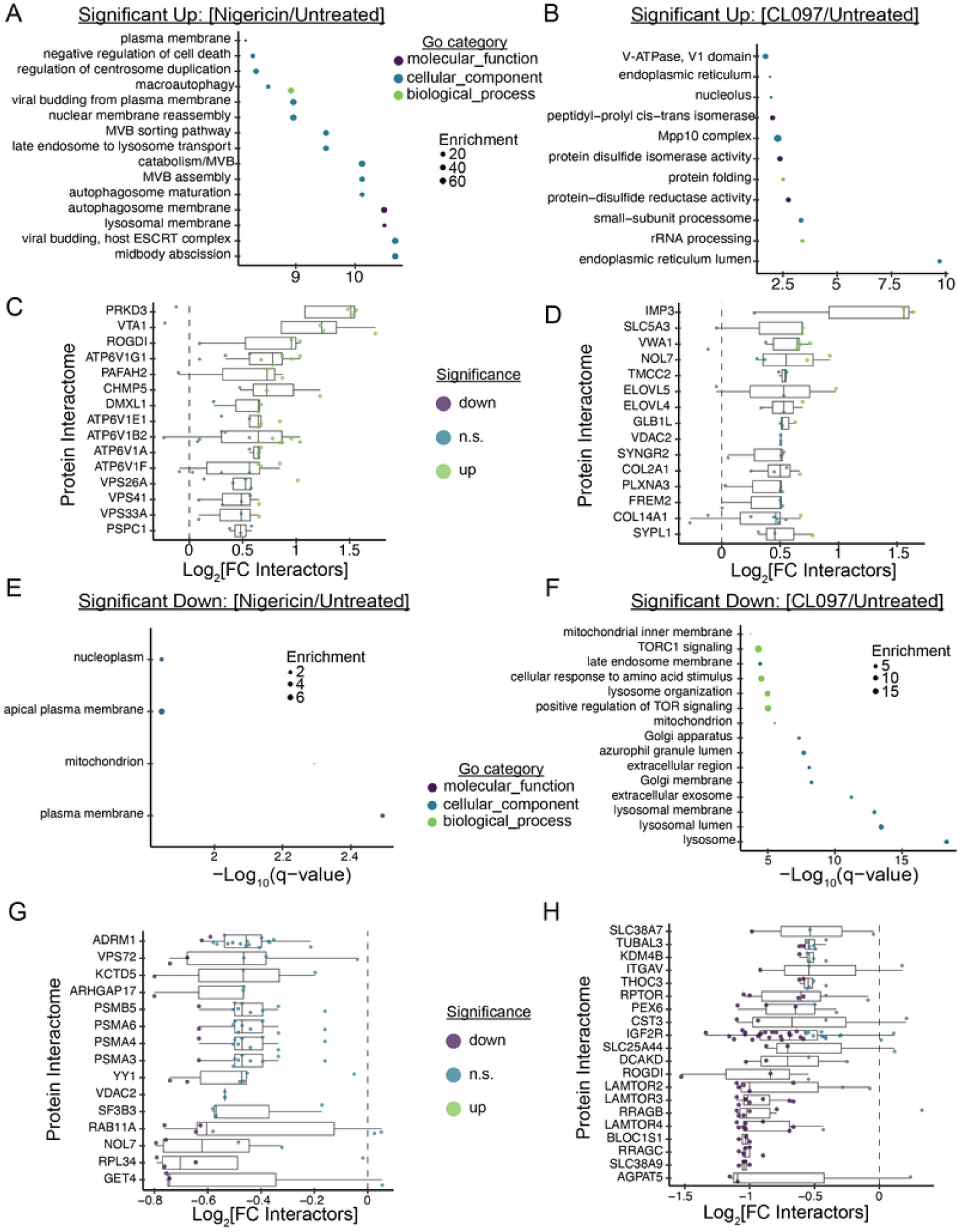
Protein group analysis for the MitoIP experiment. (A-B) GO-term analysis for proteins that significantly increased in the indicated MitoIP versus untreated cells (q < 0.05, Log_2_ FC > 0.5). (C-D) Analysis of the top 20 Bioplex protein interactomes (>3 proteins) positively enriched in the indicated experiment. (E-F) GO-term analysis for proteins that significantly decreased in the indicated MitoIP versus untreated cells (q > 0.05, Log_2_ FC < −0.5). (G-H) Analysis of the top 20 Bioplex protein interactomes (>3 proteins) depleted in the indicated experiment. To Figure 3: https://harperlab.pubpub.org/pub/nlrp3#nyjwc0zexqn

Figure 4 **supporting figures**

**Figure 4—Supporting Figure 1.**
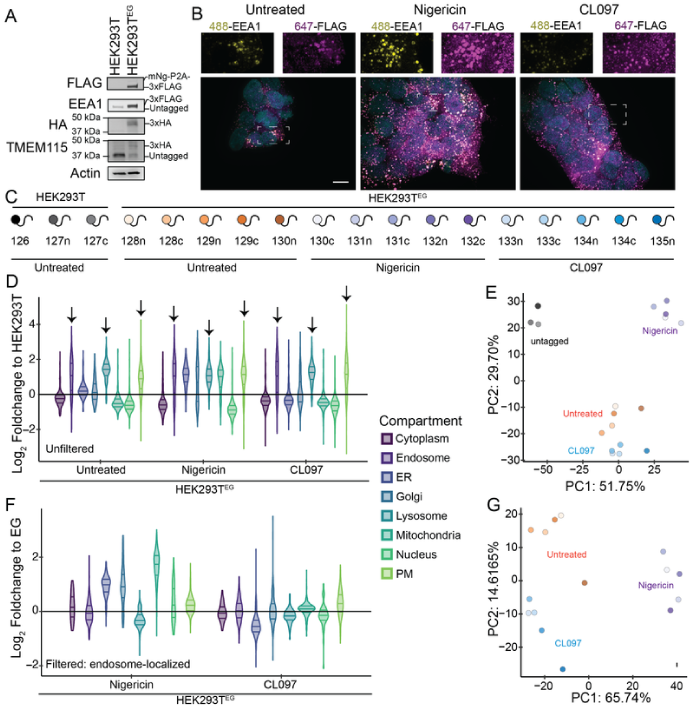
Experimental validation and design for the EndoIP experiment. (A) Western blot validation of HEK293T^EG^ cells. HEK293T cells were endogenously tagged with the EndoTag (3xFLAG-EEA1) and the GolgiTag (TMEM115-3xHA), then immunobloted with the indicated antibodies. (B) The EndoTag (3xFLAG-EEA1) localizes to endosomes and maintains localization following treatment. HEK293^EG^ cells were treated with nigericin (20 μM, 30 min) or CL097 (75 μg/mL, 1 h), fixed, and immunostained with the indicated antibody and Hoechst. Maximum intensity projection images (z=8 μM, 29 steps), representative of n > 6 fields of view. Scale bar (left panel), 10 μm. (C) TMTplex experimental design. Cells were treated with the indicated compounds (nigericin, 20 μM, 30 min; CL097, 75 μg/mL, 1 h) prior to EndoIP on anti-FLAG magnetic beads. The parental untagged HEK293T cell line serves as a background control, whereas the endogenously tagged HEK293T cell line (HEK293T^EG^) facilitates EndoIP. (D) Violin plots depicting the log_2_FC values for the indicated subcellular compartment in aggregate for the indicated EndoIP compared to the background control (HEK293T). Endosomal, lysosomal, and plasma membrane proteins (groups that likely traffic through endosomes) are enriched over all organellar annotations for each condition (indicated by arrows), validating the approach. (E) Principal component analysis (PCA) colored as in (C) with all channels included. Replicates of a given condition correlate well. (F) Violin plots depicting the log_2_ FC values for the indicated organelle annotation and the given treatment condition compared to untreated (EndoTag) cells. Proteins were first filtered for significant endosomal localization (any 293T^EG^/293T: q < 0.05, Log_2_ FC > 0.5). (G) PCA colored as in (C) with background samples excluded. Proteins were first filtered for significant endosomal localization (any 293T^EG^/293T: q < 0.05, Log_2_ FC > 0.5). Replicates of a given condition correlate well. To Figure 4: https://harperlab.pubpub.org/pub/nlrp3#n75ph0ar597

**Figure 4—Supporting Figure 2.**
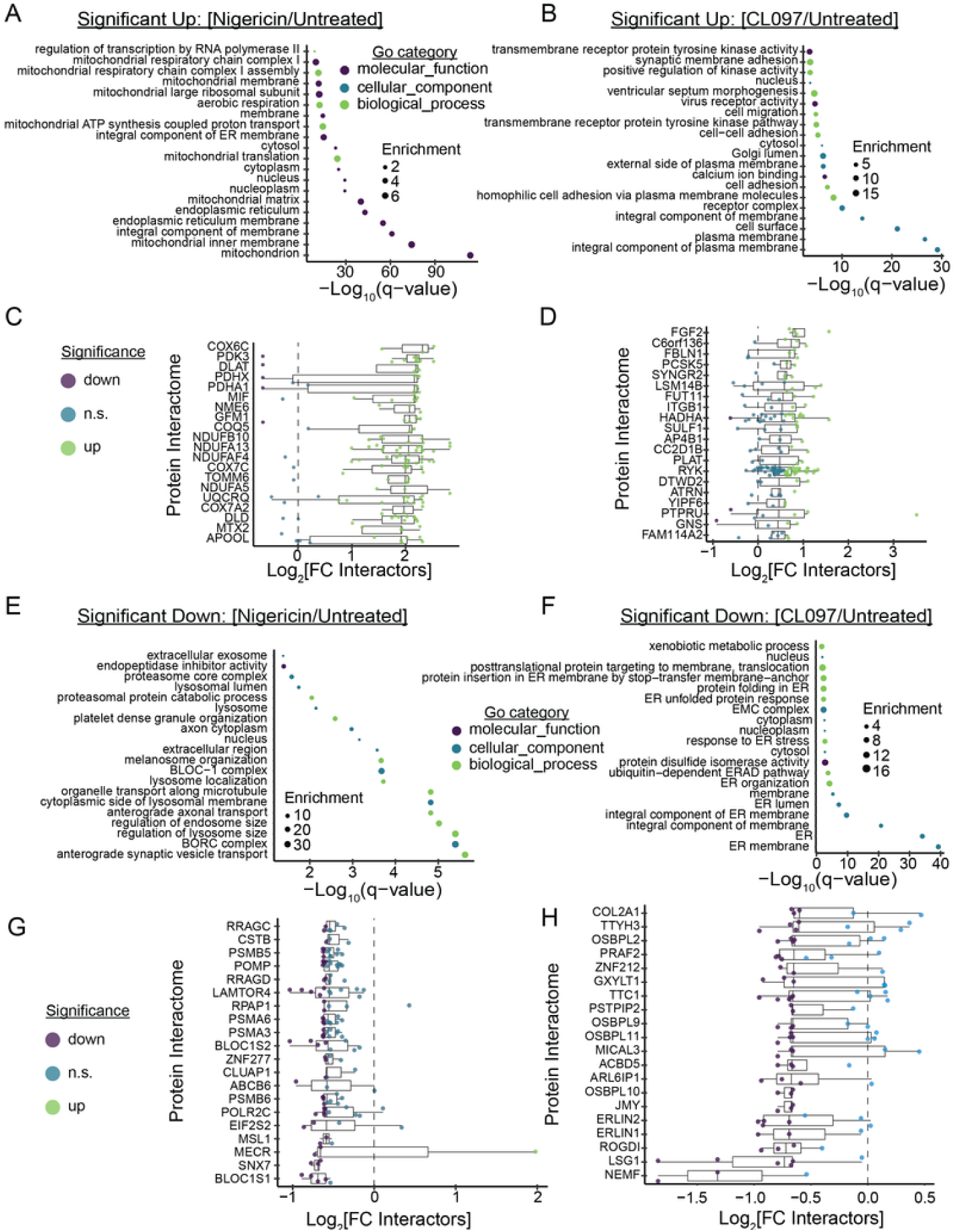
Protein group analysis for the EndoIP experiment. (A-B) GO-term analysis for proteins that significantly increased in the indicated EndoIP versus untreated cells (q < 0.05, Log_2_ FC > 0.5). (C-D) Analysis of the top 20 Bioplex protein interactomes (>3 proteins) positively enriched in the indicated experiment. (E-F) GO-term analysis for proteins that significantly decreased in the indicated EndoIP versus untreated cells (q < 0.05, Log_2_ FC < −0.5). (G-H) Analysis of the top 20 Bioplex protein interactomes (>3 proteins) depleted in the indicated experiment. To Figure 4: https://harperlab.pubpub.org/pub/nlrp3#n75ph0ar597

Figure 5 **supporting figures**

**Figure 5—Supporting Figure 1.**
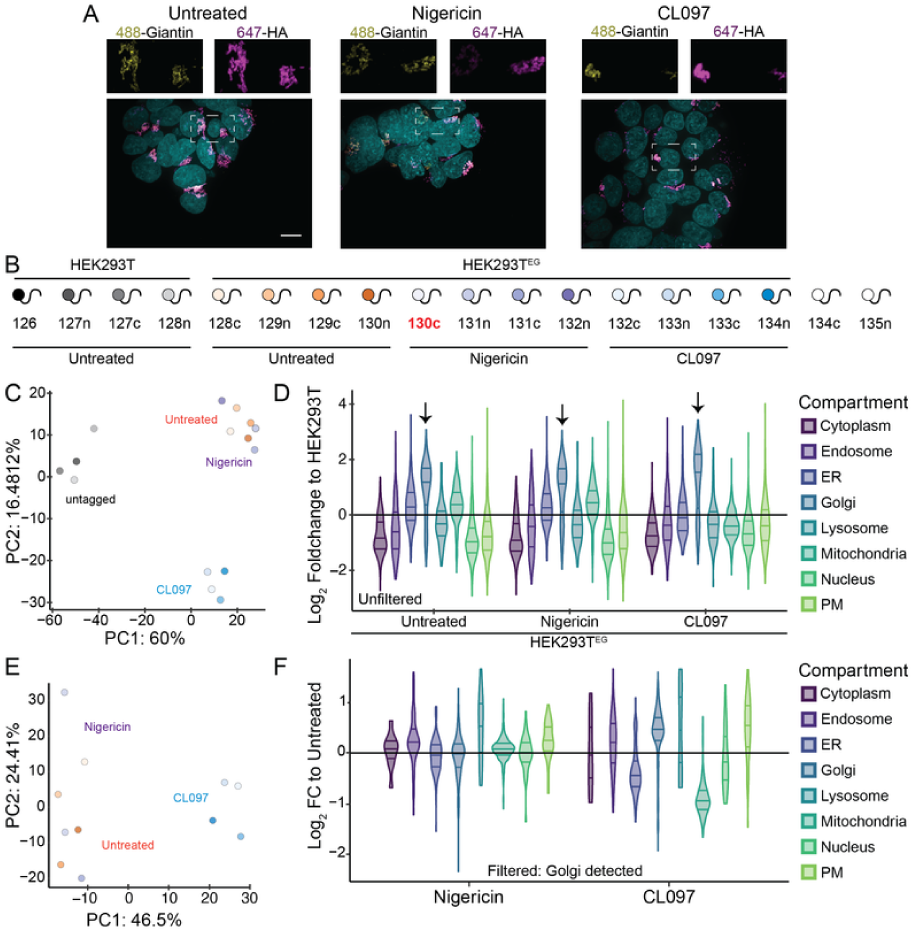
Experimental validation and design for the GolgiIP (nigericin, CL097) experiment. (A) The GolgiTag (TMEM115-3xHA) localizes to Golgi and maintains localization following treatment. HEK293T^EG^ cells were treated with nigericin (20 μM, 30 min) or CL097 (75 μg/mL, 1 h), fixed, and immunostained with the indicated antibody and Hoechst. Maximum intensity projection images (z=8 μM, 29 steps), representative of n>4 fields of view. Scale bar (left panel), 10 μm. (B) TMTplex experimental design. Cells were treated with the indicated compounds (nigericin, 20 μM, 30 min; CL097, 75μg/mL, 1 h) prior to GolgiIP on anti-HA magnetic beads. The parental untagged HEK293T cell line serves as a background control, whereas the endogenously tagged HEK293T cell line (HEK293T^EG^) facilitates GolgiIP. Channel 130n (bold red) is an outlier (insufficient material following IP as detected by the ratio check) and was excluded from analysis. (C) Principal component analysis (PCA) colored as in (B) with all channels included. Replicates of a given condition correlate well. (D) Violin plots depicting the log_2_ fold change (FC) values for the indicated subcellular compartment in aggregate for the indicated GolgiIP compared to the background control (HEK293T). GolgiIP predominantly enriches for Golgi proteins (indicated by arrows) across all experimental conditions. (E) PCA colored as in (B) with background samples excluded. Replicates of a given condition correlate well. (F) Violin plots depicting the log2 FC values for the indicated organelle annotation and the given treatment condition compared to untreated (GolgiTag) cells. Proteins were first filtered for significant Golgi localization (any 293T^EG^/293T: q < 0.05, Log_2_FC > 0.5). To Figure 5: https://harperlab.pubpub.org/pub/nlrp3#nkrmiksruzv

**Figure 5—Supporting Figure 2.**
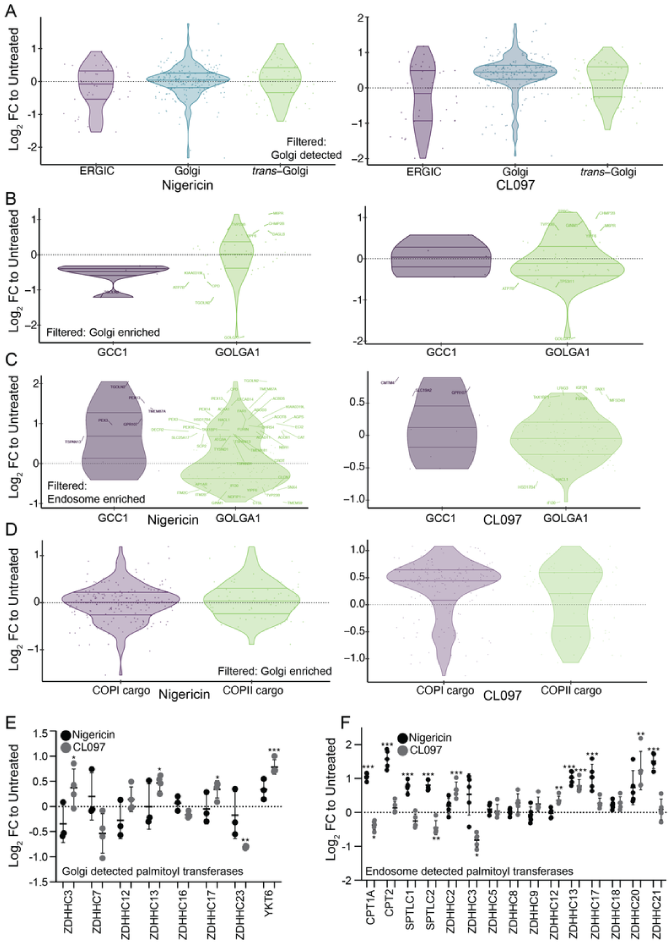
Golgi-related organellar terms and palmitoyltransferases. (A) Violin plots depicting the log_2_ fold change (FC) values for the indicated Golgi/ER subcompartment in aggregate for the indicated GolgiIP compared to the untreated control IP (HEK293T^EG^). Only Golgi enriched proteins are displayed (Log_2_FC > 0.5, q < 0.05 compared to the untagged control). Only Golgi enriched proteins are displayed (any 293T^EG^/293T: q < 0.05, Log_2_FC > 0.5). Annotations reported in Hein*, Peng*, Todorova*, McCarthy*, Kim*, and Liu* *et al.* (Leiden cluster annotations) [84]. (B-C) Violin plots depicting the log_2_FC values for GOLGA1/GCC1 cargo proteins in aggregate for the indicated (B) GolgiIP or (C) EndoIP compared to the untreated control IP (HEK293T^EG^). Only proteins enriched over the matched negative control IP are displayed (any 293T^EG^/293T: q < 0.05, Log_2_FC > 0.5). Significantly changing proteins are indicated. Annotations reported in Shin *et al.* [132]. (D) Violin plots depicting the log_2_FC values for COPI/COPII cargo proteins in aggregate for the indicated GolgiIP compared to the untreated positive control IP (HEK293T^EG^). Annotations reported in Adolf* and Rhiel* *et al.* [197]. (E-F) Log_2_FC values for (E) Golgi-enriched or (F) endosome-enriched (Log_2_FC > 0.5, q < compared to the untagged control) palmitoyl transferases in response nigericin (20 μM, 30 min) or CL097 (75 μg/mL, 1 h) compared to the untreated positive control. Error bars represent the standard deviation from n=3, 4, or 5 biological replicates. *P* values (B,C,E,F) were calculated from the Student’s t-test (two sided) and adjusted for multiple hypothesis correction using the Benjamini–Hochberg approach (q-value) with MSstats. *q<0.05, **q<0.01, ***q<0.001. To Figure 5: https://harperlab.pubpub.org/pub/nlrp3#nkrmiksruzv

**Figure 5—Supporting Figure 3.**
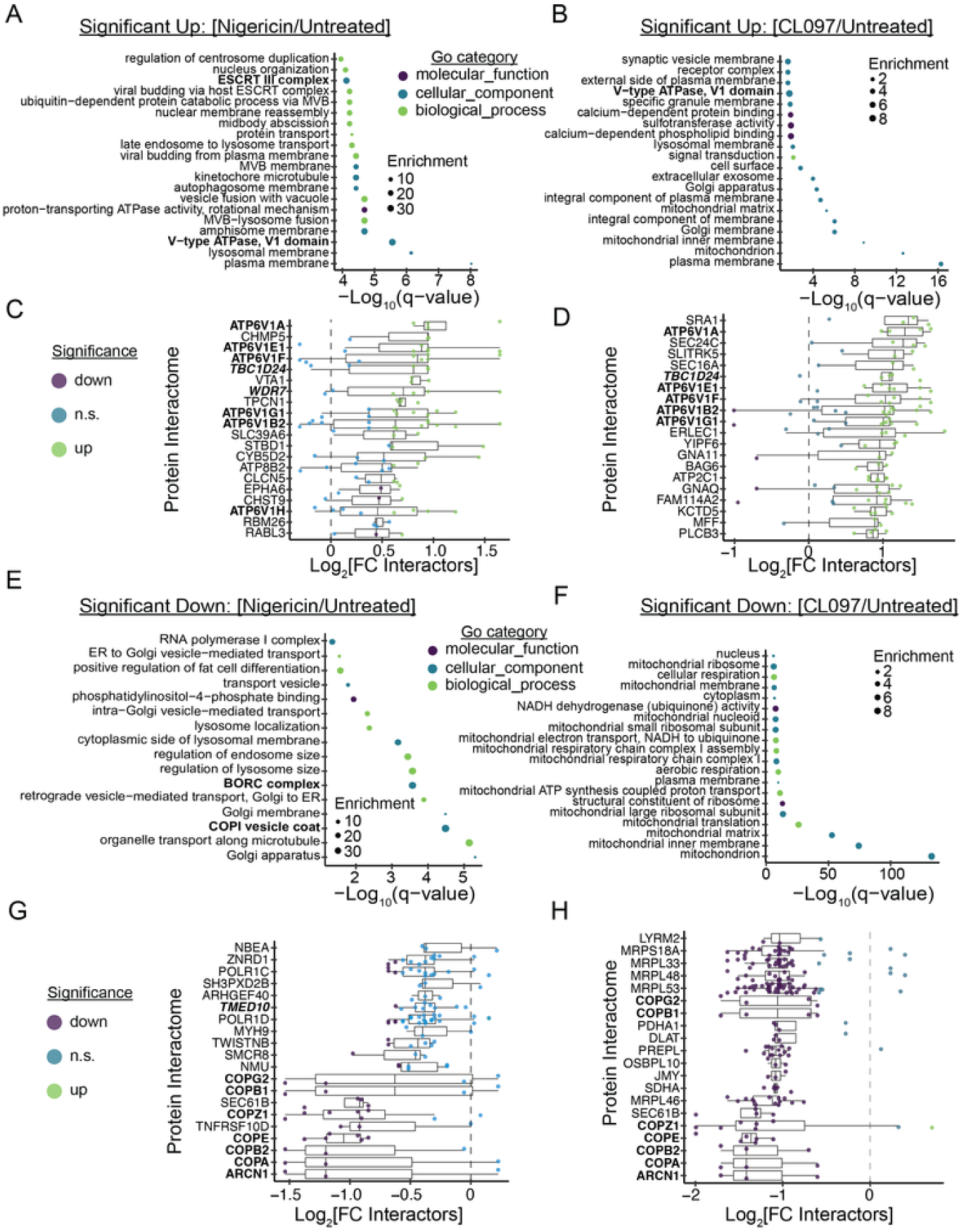
Protein group analysis for the GolgiIP experiment. (A-B) GO-term analysis for proteins that significantly increased in the indicated GolgiIP versus untreated cells (Log_2_FC > 0.5, q < 0.05). (C-D) Analysis of the top 20 Bioplex protein interactomes (>3 proteins) positively enriched in the indicated experiment. V-ATPase interactors are in bold and italicized. (E-F) GO-term analysis for proteins that significantly decreased in the indicated LysoIP versus untreated cells (Log_2_FC < −0.5, q < 0.05). (G-H) Analysis of the top 20 Bioplex protein interactomes (>3 proteins) depleted in the indicated experiment. COPI subunits are in bold, and their COPI-associated proteins are in bold and italicized. To Figure 5: https://harperlab.pubpub.org/pub/nlrp3#nkrmiksruzv

**Figure 5—Supporting Figure 4.**
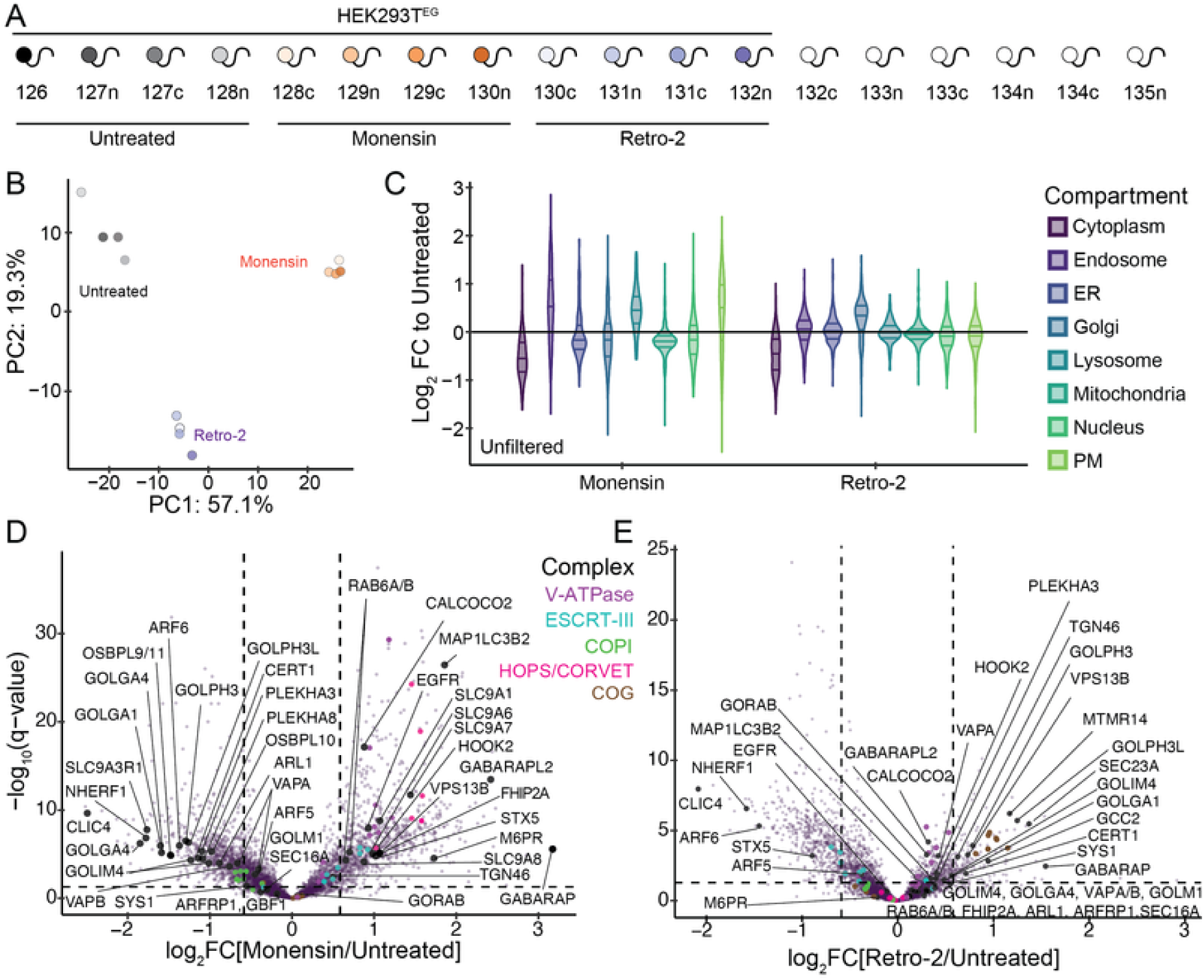
(A) TMTplex experimental design. Cells were treated with the indicated compounds (monensin, 10 μM, 2 h; retro-2, 25 μM, 2 h) prior to GolgiIP on anti-HA magnetic beads. No background control was included in this experiment. (B) Principal component analysis (PCA) colored as in (A) with all channels included. Replicates of a given condition correlate well. (C) Violin plots depicting the GolgiIP log_2_ fold change (FC) values for the indicated subcellular compartment in aggregate for the indicated treatment compared to the untreated control (HEK293T^EG^). (D-E) GolgiIP volcano plots for (D) monensin (10 μM, 2 h)-treated or (E) retro-2 (25 μM, 2 h)-treated versus untreated HEK293T^EG^ cells. Protein complexes are colored as indicated. *P* values were calculated from the Student’s t-test (two sided) and adjusted for multiple hypothesis correction using the Benjamini–Hochberg approach (q-value) with MSstats. Data represent n=4 biological replicates. To Figure 5: https://harperlab.pubpub.org/pub/nlrp3#nkrmiksruzv

**Figure 5—Supporting Figure 5.**
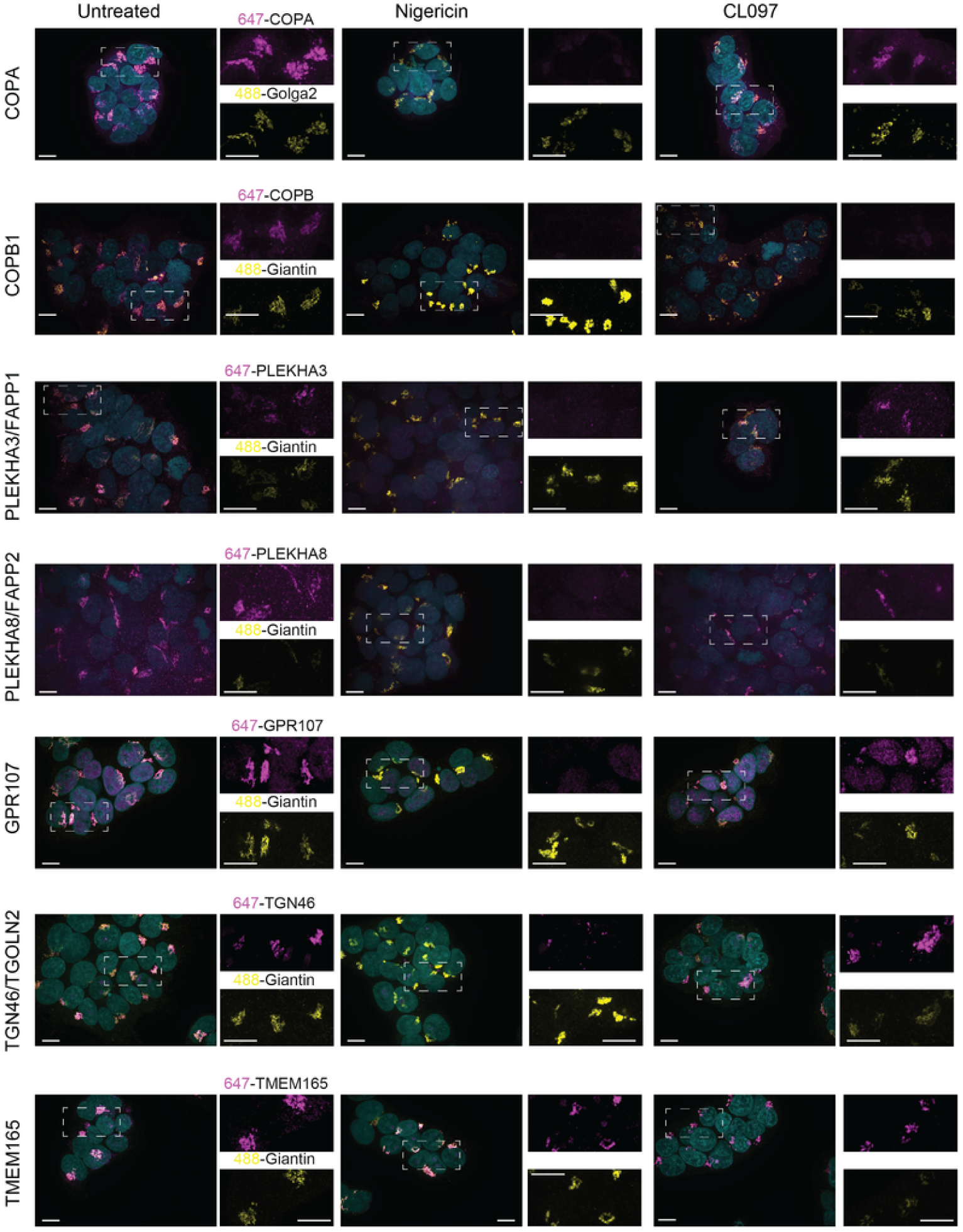
Golgi localization of COPA/B1, PLEKHA3/8, GPR107, TGN46, and TMEM165 in response to inflammasome agonists. HEK293T cells were treated with nigericin (20 μM, 30 min) or CL097 (75 μg/mL, 1 h), fixed, and immunostained with the indicated antibodies (488 and 647 channels, colored as indicated) and Hoechst (405 channel, cyan). Giantin and Golga2 are well established Golgi markers. Maximum intensity projection images (z=8 μM, 29 steps), representative of n > 6 fields of view from at least two independent experiments. Scale bars, 10 μm. To Figure 5: https://harperlab.pubpub.org/pub/nlrp3#nkrmiksruzv

Figure 6 **supporting figures**

**Figure 6—Supporting Figure 1.**
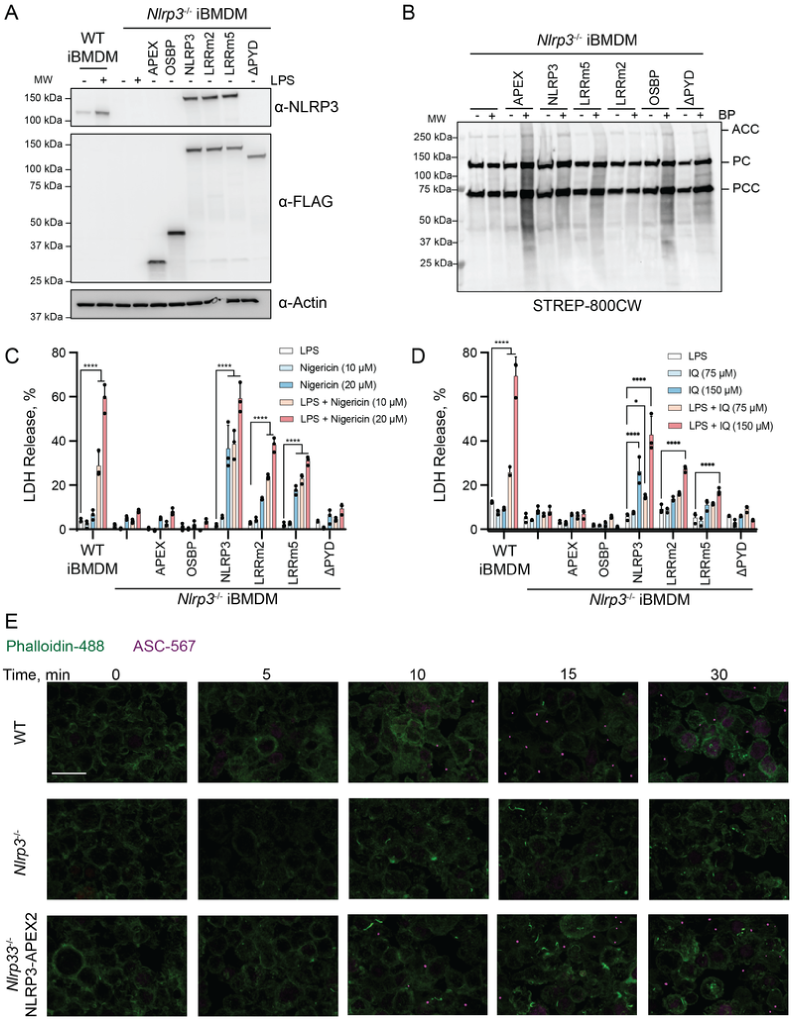
Validation of Nlrp3-APEX2 reconstitution. (A) Validation of stable Nlrp3-APEX2 reconstitution at levels similar to LPS-primed WT iBMDMs, following two FACS sorts. Immunoblots with the indicated antibodies for WT and *Nlrp3^-/-^* iBMDMs ± LPS (1 μg/mL, 4 h), in addition to *Nlrp3^-/-^* iBMDMs stably reconstituted with the indicated APEX2-FLAG construct. The cage-disrupting Nlrp3 mutations LRRm2 (N1008R, R1009E, E1010R, R1013E) and LRRm5 (H781E, Q782R, F785A) were reported in Andreeva *et al.* [37]. Nlrp3ΔPYD is I125M-END (no polybasic linker region). PH, pleckstrin homology domain. (B) Validation of APEX2 biotinylation activity. APEX2-expressing and control *Nlrp3^-/-^* iBMDMs ± biotin phenol (BP, 500 μM, 45 min) prior to labeling with H_2_O_2_ (10 μM, 1 min). Lysates were probed with STREP-800CW to detect biotinylated proteins. ACC, acetyl-CoA carboxylase. PC, pyruvate carboxylase. PCC, propionyl-CoA carboxylase. (C-D) Reconstituted Nlrp3-APEX2 rescues inflammasome activity. % LDH release (relative to cell culture media and a maximum release control) of *Nlrp3^-/-^* iBMDMs reconstituted with various APEX2 constructs. Cells were treated without LPS or primed with LPS (1 μg/mL) 4 h prior to treatment with (C) nigericin (indicated concentrations, 1 h) or (D) imiquimod (indicated concentrations, 2 h). Bar graphs represent mean ± standard deviation, n=3 replicates. *p<0.05, **p<0.01, ***p<0.001, ****p<0.0001 using a two-way ANOVA with Tukey’s post-test. (E) Timecourse of inflammasome speck assembly. The indicated primed (4 h LPS, 1 μg/mL) iBMDMs were treated with 20 μM Nigericin for the indicated periods of time, fixed, immunostained (Hoechst, Phalloidin, anti-ASC), and imaged. Maximum intensity projections are shown (z=8 μM, 29 steps). Scale bar (top left image) = 25 μm. Representative regions of n > 3 fields of view. To Figure 6: https://harperlab.pubpub.org/pub/nlrp3#nywwrxbdiu9

**Figure 6—Supporting Figure 2.**
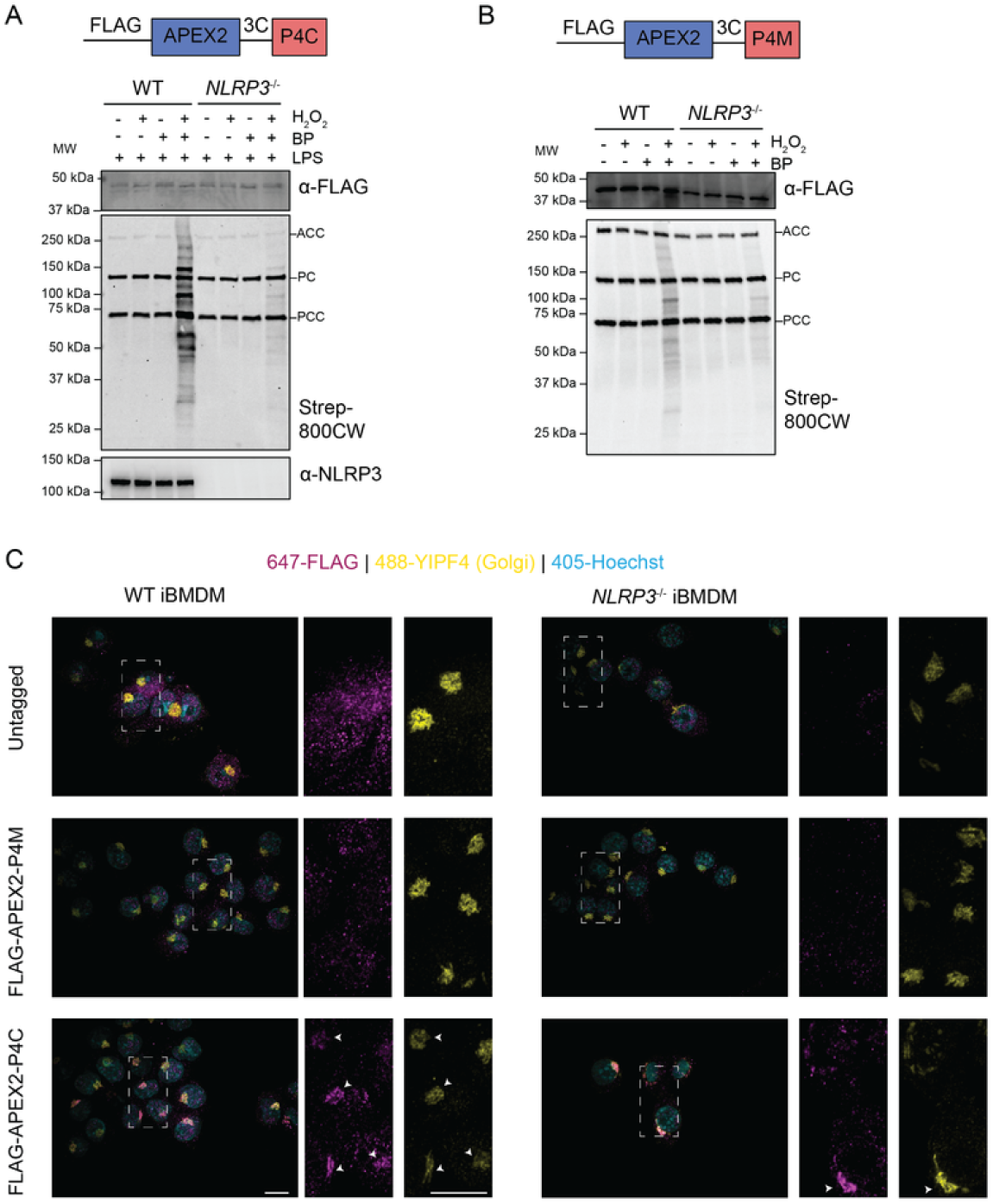
Validation of APEX2-P4C reconstitution. Validation of APEX2 biotinylation activity and PI4P biosensor Golgi localization. (A) Immunoblots of LPS-primed (1 μg/mL, 4 h) WT and *Nlrp3^-/-^* iBMDMs reconstituted with FLAG-APEX2-P4C (expected 48 kDa). Cells were stimulated ± biotin phenol (BP, 500 μM, 30 min) followed by ± H_2_O_2_ (10 μM, 1 min) prior to quenching and harvesting cells. ACC, acetyl-CoA carboxylase. PC, pyruvate carboxylase. PCC, propionyl-CoA carboxylase. n=1 biological replicate. (B) Immunoblots of WT and *Nlrp3^-/-^* iBMDMs reconstituted with FLAG-APEX2-P4M (expected 40 kDa). Cells were stimulated ± biotin phenol (BP, 500 μM, 30 min) followed by ± H_2_O_2_ (10 μM, 1 min) prior to quenching and harvesting cells. n=1 biological replicate. (C) The P4C construct demonstrates Golgi localization by immunofluorescence of fixed cells whereas the P4M construct does not. The indicated primed (4 h LPS, 1 μg/mL) iBMDMs were fixed, immunostained as indicated, and imaged with confocal microscopy. Maximum intensity projection images (z=8 μM, 29 steps) are shown. Arrows indicate colocalized FLAG (P4C construct) and YIPF4 (Golgi marker) signal. Representative images from n > 6 fields of view. Scale bar (bottom left panels), 10 μm. To Figure 6: https://harperlab.pubpub.org/pub/nlrp3#nywwrxbdiu9

**Figure 6—Supporting Figure 3.**
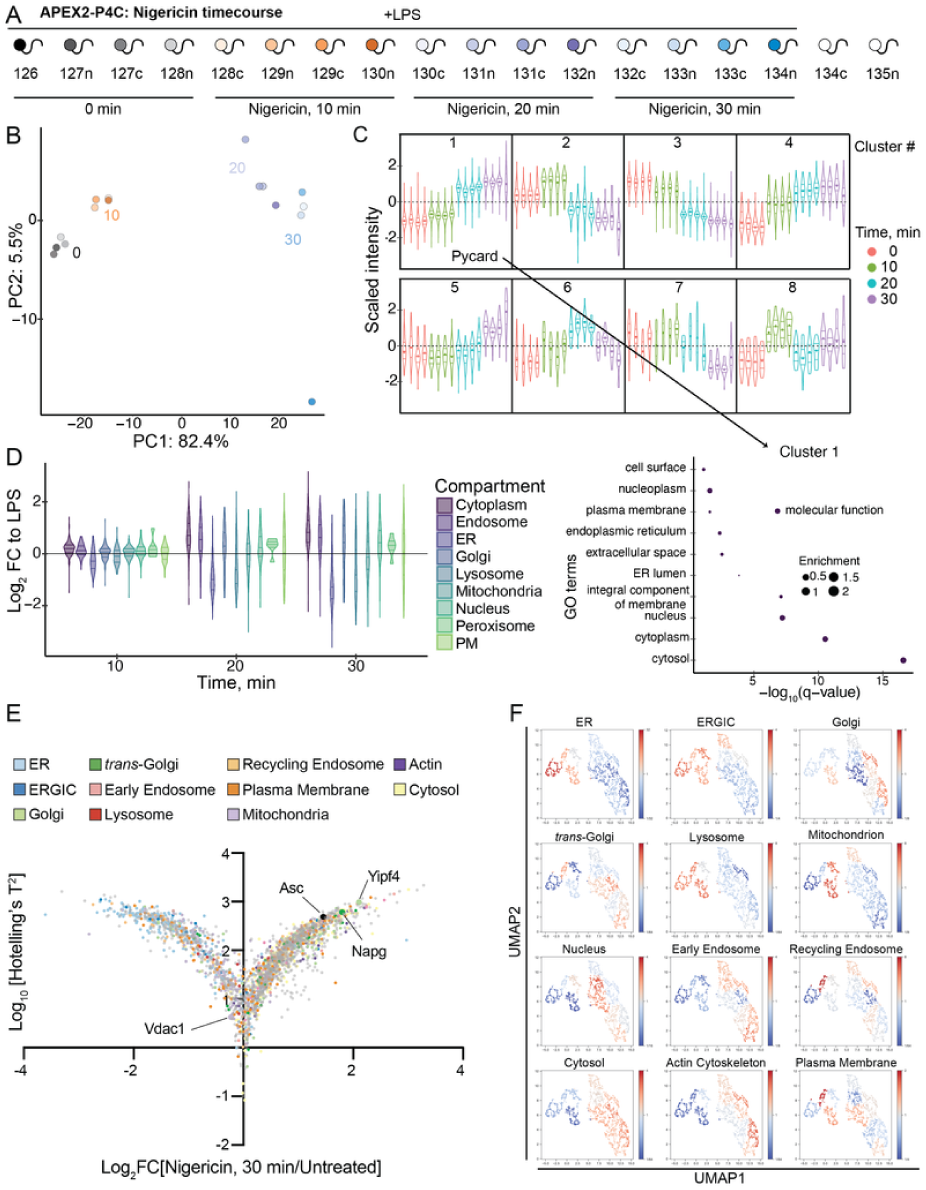
Supplemental data for the APEX2-P4C nigericin timecourse. (A) APEX2-P4C nigericin timecourse experimental design. Cells were primed with LPS (1 μg/mL, 4 h) then treated with nigericin (20 μM; 0, 10, 20, 30 min) prior to labeling with H_2_O_2_ (10 μM, 1 min). All conditions were also treated with biotin phenol (500 μM, 45 min) prior to labeling. (B) Principal component analysis (PCA) colored as in (A), showing timecourse-dependent replicate separation. (C) Hierarchical clusters of ANOVA-significant proteins within the dataset. The behavior of each cluster is shown with violin plots of the aggregated data, replicates colored as indicated. The cluster containing Pycard (Asc) is annotated. GO-term analysis is shown for the Asc-containing cluster. (D) Violin plots depicting the log_2_FC values for ER proteins detected in each experiment. Notably, proximity to ER proteins decreased rapidly. (E) Multivariate empirical Bayes analysis for total APEX2 data over the 30-minute stimulation timecourse. Plot displays log_10_(time course Hotelling’s T^2^ statistic) versus log_2_FC(30 min nigericin versus unstimulated). Proteins are colored by organelle annotation as indicated. (F) ANOVA-significant proteins the APEX2 timecourse were clustered with the Leiden algorithm. These clusters are colored as annotated on UMAP embeddings of the data by organelle enrichment. Organellar annotations for E,F reported in Hein*, Peng*, Todorova*, McCarthy*, Kim*, and Liu* *et al.* (graph-based annotations) [84]. To Figure 6: https://harperlab.pubpub.org/pub/nlrp3#nywwrxbdiu9

**Figure 6—Supporting Figure 4.**
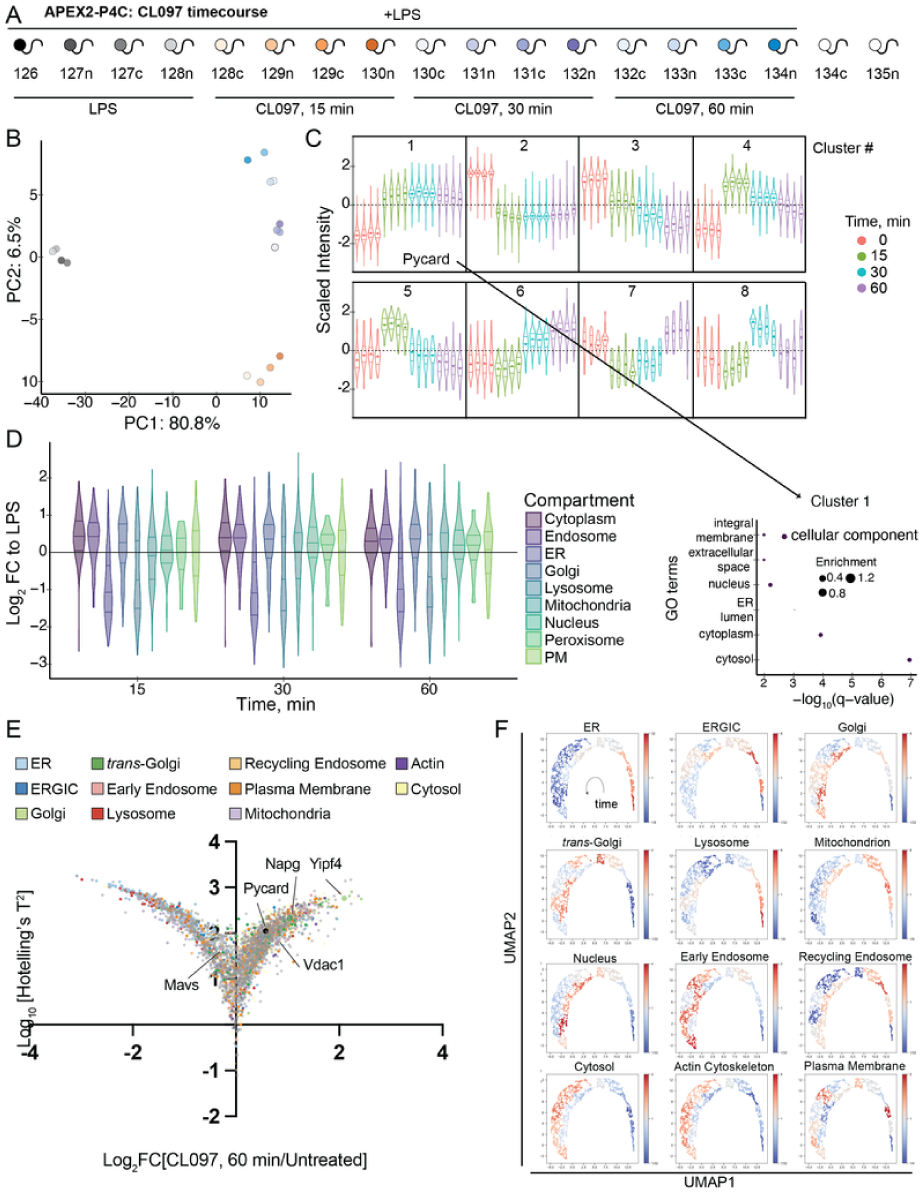
Supplemental data for the APEX2-P4C CL097 timecourse. (A) APEX2-P4C CL097 timecourse experimental design. Cells were primed with LPS (1 μg/mL, 4 h) then treated with CL097 (75 μg/mL; 0, 15, 30, 60 min) prior to labeling with H_2_O_2_ (10 μM, 1 min). All conditions were also treated with biotin phenol (500 μM, 45 min) prior to labeling. (B) Principal component analysis (PCA) colored as in (A), showing timecourse-dependent replicate separation. (C) Hierarchical clusters of ANOVA-significant proteins within the dataset. The behavior of each cluster is shown with violin plots of the aggregated data, replicates colored as indicated. The cluster containing Pycard (Asc) is annotated. GO-term analysis is shown for the Asc-containing cluster. (D) Violin plots depicting the log_2_FC values for ER proteins detected in each experiment. Notably, proximity to ER and lysosomal proteins decreased rapidly. (E) Multivariate empirical Bayes analysis for total APEX2 data over the 60-minute stimulation timecourse. Plot displays log_10_(time course Hotelling’s T^2^ statistic) versus log_2_(60 min CL097 versus unstimulated). Proteins are colored by organelle annotation as indicated. (F) ANOVA-significant proteins the APEX2 timecourse were clustered with the Leiden algorithm. These clusters are colored as annotated on UMAP embeddings of the data by organelle enrichment. Organellar annotations for E,F reported in Hein*, Peng*, Todorova*, McCarthy*, Kim*, and Liu* *et al*. (graph-based annotations) [84]. To Figure 6: https://harperlab.pubpub.org/pub/nlrp3#nywwrxbdiu9

**Figure 6—Supporting Figure 5.**
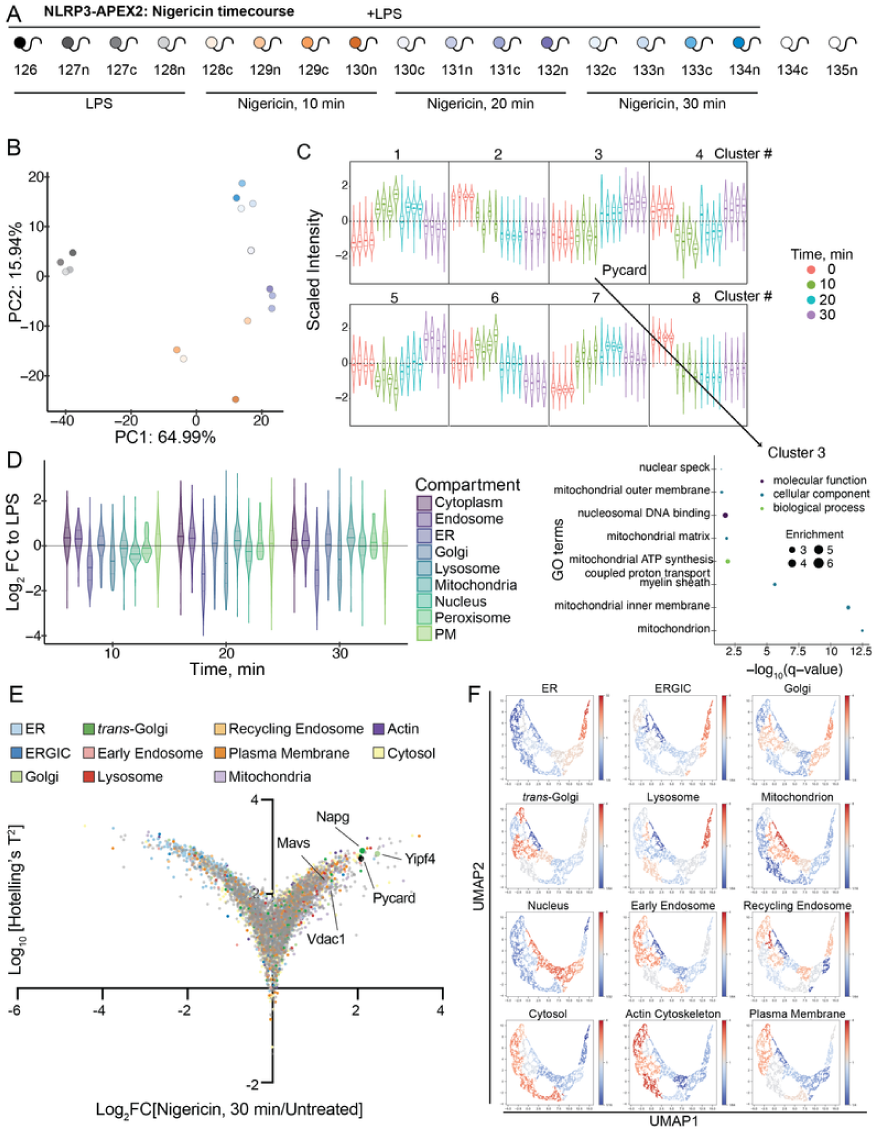
Supplemental data for the Nlrp3-APEX2 nigericin timecourse. A) Nlrp3-APEX nigericin timecourse experimental design. Cells were primed with LPS (1 μg/mL, 4 h) then treated with nigericin (20 μM; 0, 10, 20, 30 min) prior to labeling with H_2_O_2_ (10 μM, 1 min). All conditions were also treated with biotin phenol (500 μM, 45 min) prior to labeling. (B) Principal component analysis (PCA) colored as in (A), showing timecourse-dependent replicate separation. (C) Hierarchical clusters of ANOVA-significant proteins within the dataset. The behavior of each cluster is shown with violin plots of the aggregated data, replicates colored as indicated. The cluster containing Pycard (Asc) is annotated. GO-term analysis is shown for the Asc-containing cluster. (D) Violin plots depicting the log_2_FC values for ER proteins detected in each experiment. Notably, proximity to ER and lysosomal proteins decreased rapidly. (E) Multivariate empirical Bayes analysis for total APEX2 data over the 30-minute stimulation timecourse. Plot displays log_10_(time course Hotelling’s T^2^ statistic) versus log_2_(30 min nigericin versus unstimulated). Proteins are colored by organelle annotation as indicated. (F) ANOVA-significant proteins the APEX2 timecourse were clustered with the Leiden algorithm. These clusters are colored as annotated on UMAP embeddings of the data by organelle enrichment. Organellar annotations for E,F reported in Hein*, Peng*, Todorova*, McCarthy*, Kim*, and Liu* *et al*. (graph-based annotations) [84]. To Figure 6: https://harperlab.pubpub.org/pub/nlrp3#nywwrxbdiu9

**Figure 6—Supporting Figure 6.**
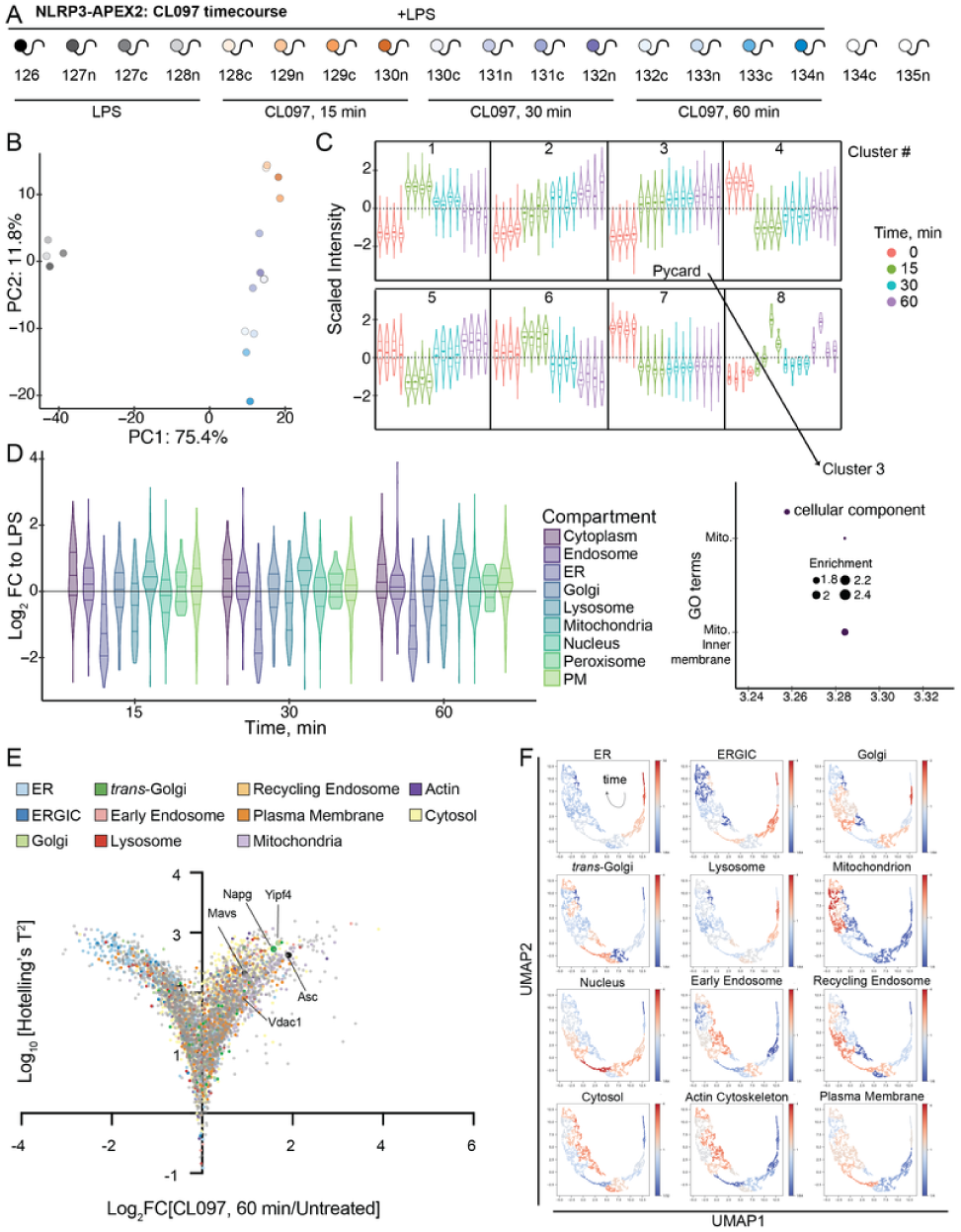
Supplemental data for the Nlrp3-APEX2 CL097 timecourse. (A) Nlrp3-APEX2 CL097 timecourse experimental design. Cells were primed with LPS (1 μg/mL, 4 h) then treated with CL097 (75 μg/mL; 0, 15, 30, 60 min) prior to labeling with H_2_O_2_ (10 μM, 1 min). All conditions were also treated with biotin phenol (500 μM, 45 min) prior to labeling. (B) Principal component analysis (PCA) colored as in (A), showing timecourse-dependent replicate separation. (C) Hierarchical clusters of ANOVA-significant proteins within the dataset. The behavior of each cluster is shown with violin plots of the aggregated data, replicates colored as indicated. The cluster containing Pycard (Asc) is annotated. GO-term analysis is shown for the Asc-containing cluster. (D) Violin plots depicting the log_2_FC values for ER proteins detected in each experiment. Notably, proximity to ER and lysosomal proteins decrease rapidly. (E) Multivariate empirical Bayes analysis for total APEX2 data over the 60-minute stimulation timecourse. Plot displays log_10_(time course Hotelling’s T^2^ statistic) versus log_2_ (60 min CL097 versus unstimulated). Proteins are colored by organelle annotation as indicated. (F) ANOVA-significant proteins the APEX2 timecourse were clustered with the Leiden algorithm. These clusters are colored as annotated on UMAP embeddings of the data by organelle enrichment. Organellar annotations for E,F reported in Hein*, Peng*, Todorova*, McCarthy*, Kim*, and Liu* *et al*. (graph-based annotations) [84].

**Figure 7—Supporting Figure 1.**
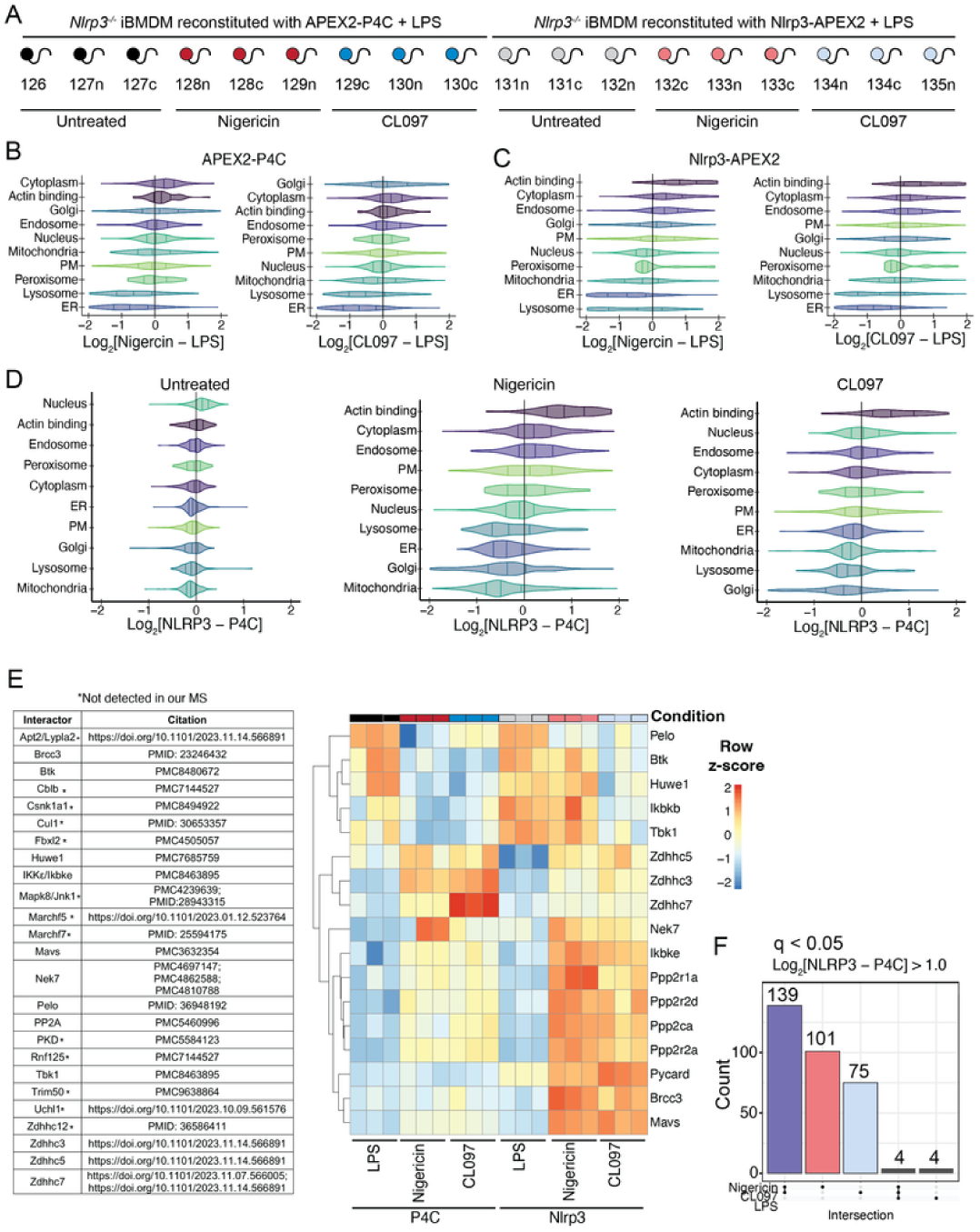
Supplemental data for proximity labeling with P4C and Nlrp3 in the same TMTplex. (A) Combined APEX2 experiment design. *Nlrp3^-/-^* iBMDMs expressing either APEX2-P4C (background control) or Nlrp3-APEX2 were primed with LPS (1 μg/mL, 4 h) then treated with nigericin (20 μM, 20 min) or CL097 (75 μg/mL, 25 min) prior to labeling with H_2_O_2_ (10 μM, 1 min). All conditions were also treated with biotin phenol (500 μM, 45 min) prior to labeling. (B-C) Violin plots depicting the log_2_FC values across the indicated subcellular compartments for the given treatment condition versus untreated cells. Comparisons are either for *Nlrp3^-/-^* iBMDMs expressing (B) APEX2-P4C or (C) Nlrp3-APEX2. (D) Violin plots depicting the log_2_FC values across the indicated subcellular compartments for the indicated Nlrp3-APEX2 condition to its matched P4C-APEX2 background condition. (E) Summary of high confidence Nlrp3 interactors from the literature (left) and a heatmap of those interactors present in the dataset (right), showing row z-score as annotated. (F) UpSet plot of proteins enriched in Nlrp3-APEX2 over APEX2-P4C (any matched comparison of Nlrp3/P4C: q < 0.05, Log_2_FC > 1.0). Intersections show multiple instances of Nlrp3-APEX2 enrichment over the APEX2-P4C background. To Figure 7: https://harperlab.pubpub.org/pub/nlrp3#njz6bdmzuz6

## Supplemental data, resources, and reagents

### Supplemental tables

Supplemental tables are hosted on Zenodo: 10.5281/zenodo.10975767

**Table S1.** Whole cell proteomics data and analysis (associated with Figure 2).

**Table S2.** LysoIP proteomics data and analysis (associated with Figure 3, Figure 3—Supporting Data Figure 1).

**Table S3.** MitoIP proteomics data and analysis (associated with Figure 3, Figure 3—Supporting Data Figure 3).

**Table S4.** EndoIP proteomics data and analysis (associated with Figure 4, Figure 4—Supporting Data Figure 1).

**Table S5.** GolgiIP (nigericin, CL097) proteomics data and analysis (associated with Figure 5, Figure 5—Supporting Data Figure 1).

**Table S6.** GolgiIP (monensin, retro-2) proteomics data and analysis (associated with Figure 5, Figure 5—Supporting Data Figure 4).

**Table S7.** APEX2 timecourse (P4C, nigericin) proteomics data and analysis (associated with Figure 6, Figure 6—Supporting Data Figure 3).

**Table S8.** APEX2 timecourse (P4C, CL097) proteomics data and analysis (associated with Figure 6, Figure 6—Supporting Data Figure 4).

**Table S9.** APEX2 timecourse (Nlrp3, nigericin) proteomics data and analysis (associated with Figure 6, Figure 6—Supporting Data Figure 5).

**Table S10.** APEX2 timecourse (Nlrp3, CL097) proteomics data and analysis (associated with Figure 6, Figure 6—Supporting Data Figure 6).

**Table S11.** Meta cluster data across timecourse data, and linear model timecourse data (associated with Figure 6).

**Table S12.** APEX2 combined (P4C and Nlrp3; CL097 and nigericin) proteomics data and analysis (associated with Figure 7).

**Table S13.** APEX2 priming (Nlrp3 and P4C, ±LPS) proteomics data and analysis (associated with Negative Data Figure 5).

**Table S14.** Organellar annotations (associated with multiple figures).

**Table S15.** MS data acquisition parameters and key resources used or made in this study. Includes cell lines, plasmids, sgRNA sequences, antibodies, and other reagents.

### Additional data, code, and resource availability

- Uncropped scans, raw confocal microscopy images, and other raw data will be made available on Zenodo in revision.
- Full sized PDFs of the figures are available on OSF: https://osf.io/z6j5e/
- Validated plasmids have been deposited in Addgene (see <u>Table S15</u>; some might be pending quality control). All other plasmids are available on request.
- Cell lines produced in this study (see <u>Table S15</u>) are available on reasonable request. We ask that all derivatives of these lines be made openly available to the research community without restrictive MTAs, in perpetuity.
- The mass spectrometry proteomics data have been deposited to the ProteomeXchange Consortium [185] via the PRIDE [186] partner repository with the dataset identifier PXD051489.
- Processed proteomics data are available as [supplemental data files above](https://harperlab.pubpub.org/pub/nlrp3/release/1#n9x2jmnbspr) and can also be browsed on our Shiny app: https://harperlab.connect.hms.harvard.edu/inflame/
- Code is available on https://github.com/priyaveeraraghavan/inflame
- The following extended protocols have been made available on protocols.io:

- dx.doi.org/10.17504/protocols.io.dm6gpzr3plzp/v1
- dx.doi.org/10.17504/protocols.io.kxygxyeeol8j/v1
- dx.doi.org/10.17504/protocols.io.5qpvok55zl4o/v1
- dx.doi.org/10.17504/protocols.io.261ge5p57g47/v1

### Materials acknowledgements

WT and *Nlrp3^-/-^* iBMDMs were a gift from Kate Fitzgerald (UMass Chan Medical School). HEK293 cells endogenously expressing the LysoTag were previously engineered in our lab [88]. pMD2.G was a gift from Didier Trono (Addgene plasmid # 12259; http://n2t.net/addgene:12259; RRID:Addgene_12259). psPAX2 was a gift from Didier Trono (Addgene plasmid # 12260; http://n2t.net/addgene:12260; RRID:Addgene_12260). LentiCRISPRv2-Opti was a gift from David Sabatini (Addgene plasmid # 163126; http://n2t.net/addgene:163126; RRID:Addgene_163126). LentiCRISPRv2-mCherry was a gift from Agata Smogorzewska (Addgene plasmid # 99154; http://n2t.net/addgene:99154; RRID:Addgene_99154). plenti-UBC-gate-3xHA-pGK-PUR was a gift from William Kaelin (Addgene plasmid # 107393; http://n2t.net/addgene:107393; RRID:Addgene_107393).

We thank bioicons (https://bioicons.com/) for aggregating open-sourced figures. Receptor-membrane-2, golgi-3d-2, dna-5 icons by Server (https://smart.servier.com/) are licensed under CC-BY 3.0 Unported (https://creativecommons.org/licenses/by/3.0/). The generic-bacterium icon by Pauline Franz (https://twitter.com/_paulinefranz) is licensed under CC0 (https://creativecommons.org/publicdomain/zero/1.0/). Mitochondria and Endoplasmic_Reticulum icons by jaiganesh (https://github.com/jaiganeshjg) are licensed under CC0 (https://creativecommons.org/publicdomain/zero/1.0/).

### Antibodies

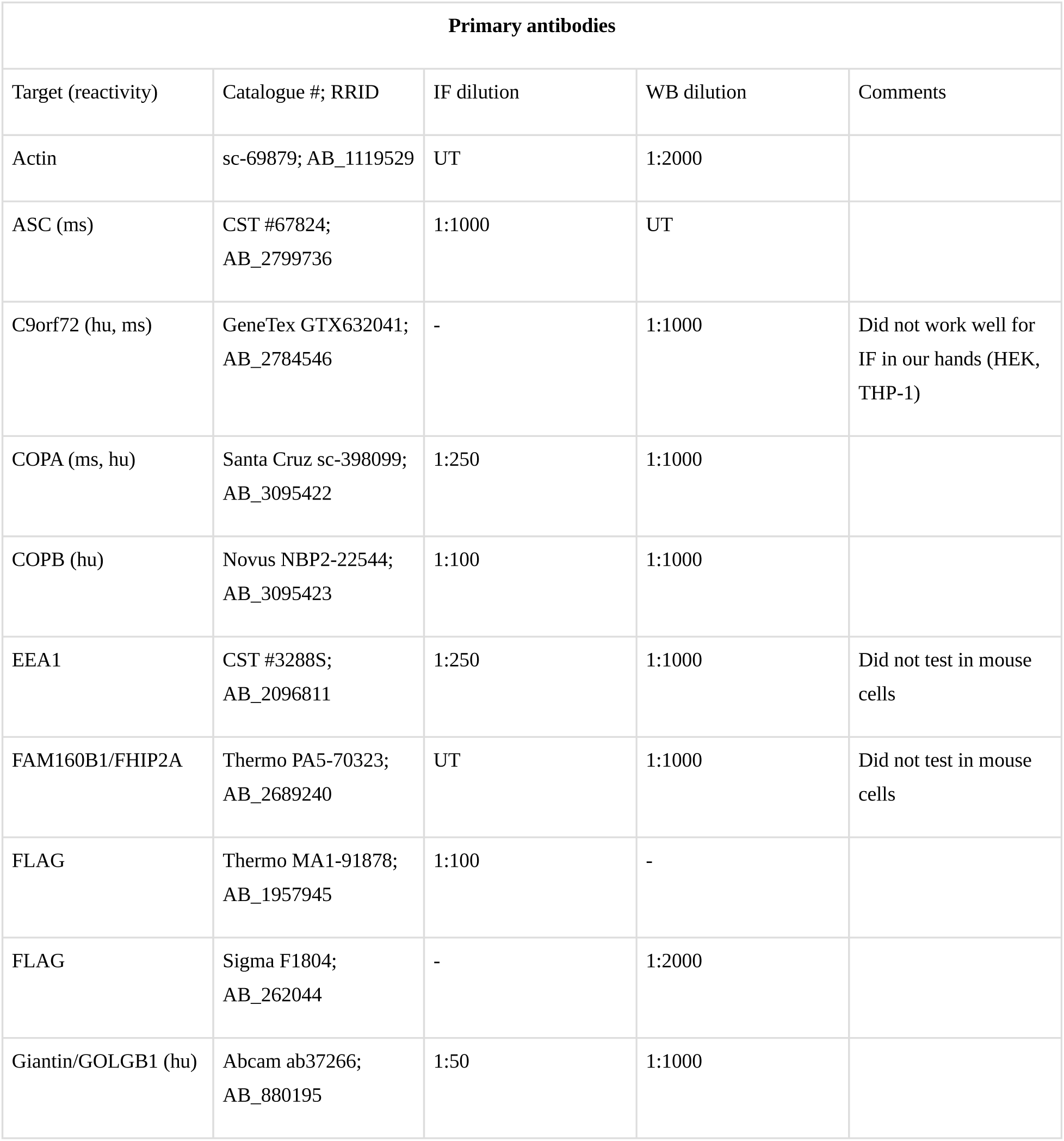
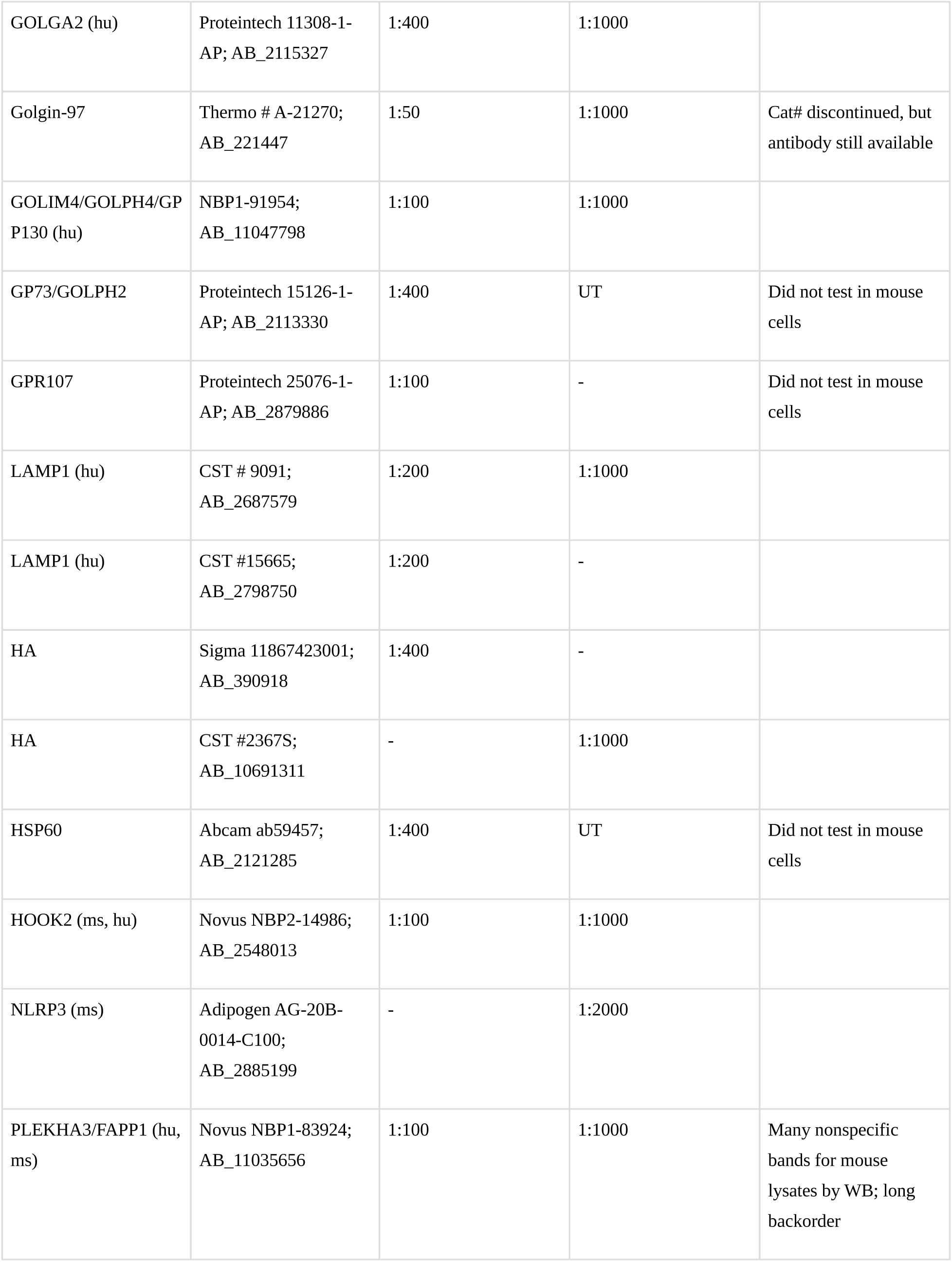
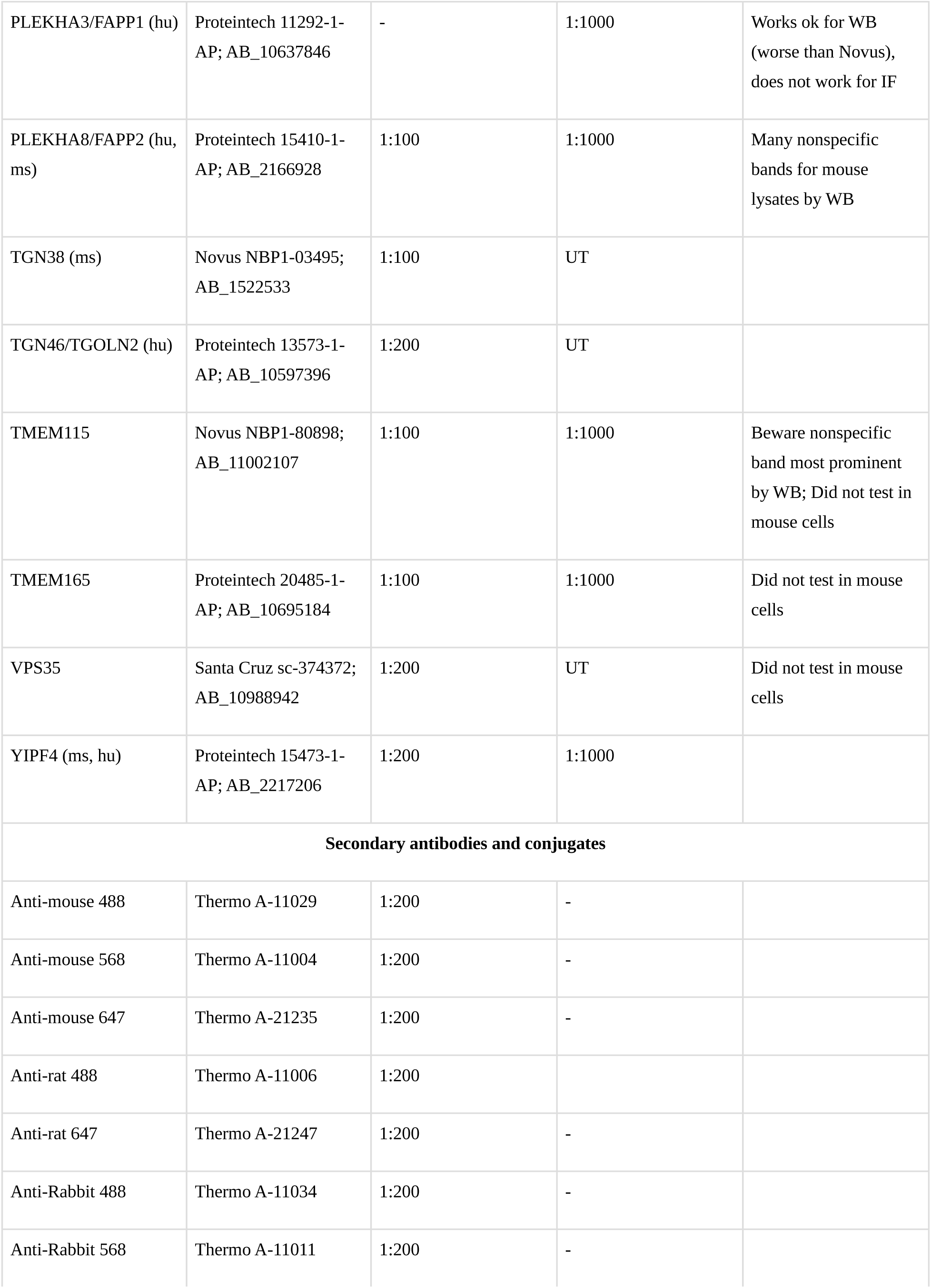
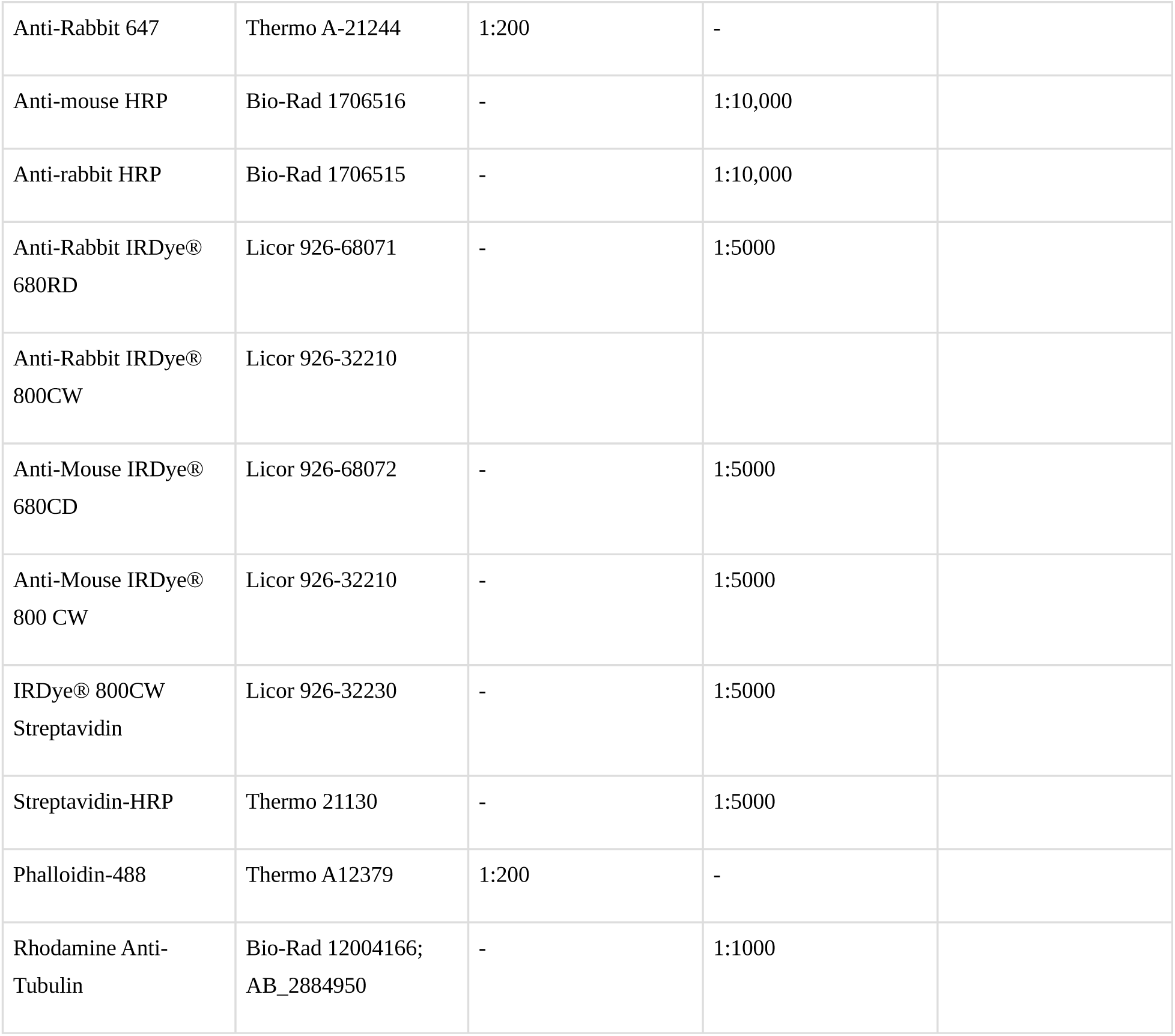

| Primary antibodies |  |  |  |  |
| --- | --- | --- | --- | --- |
| Target (reactivity) | Catalogue #; RRID | IF dilution | WB dilution | Comments |
| Actin | sc-69879; AB_1119529 | UT | 1:2000 |  |
| ASC (ms) | CST #67824;<br>AB_2799736 | 1:1000 | UT |  |
| C9orf72 (hu, ms) | GeneTex GTX632041;<br>AB_2784546 | - | 1:1000 | Did not work well for IF in our hands (HEK, THP-1) |
| COPA (ms, hu) | Santa Cruz sc-398099;<br>AB_3095422 | 1:250 | 1:1000 |  |
| COPB (hu) | Novus NBP2-22544;<br>AB_3095423 | 1:100 | 1:1000 |  |
| EEA1 | CST #3288S;<br>AB_2096811 | 1:250 | 1:1000 | Did not test in mouse cells |
| FAM160B1/FHIP2A | Thermo PA5-70323;<br>AB_2689240 | UT | 1:1000 | Did not test in mouse cells |
| FLAG | Thermo MA1-91878;<br>AB_1957945 | 1:100 | - |  |
| FLAG | Sigma F1804;<br>AB_262044 | - | 1:2000 |  |
| Giantin/GOLGB1 (hu) | Abcam ab37266;<br>AB_880195 | 1:50 | 1:1000 |  |
| GOLGA2 (hu) | Proteintech 11308-1-AP; AB_2115327 | 1:400 | 1:1000 |  |
| Golgin-97 | Thermo # A-21270; AB_221447 | 1:50 | 1:1000 | Cat# discontinued, but antibody still available |
| GOLIM4/GOLPH4/GP P130 (hu) | NBP1-91954; AB_11047798 | 1:100 | 1:1000 |  |
| GP73/GOLPH2 | Proteintech 15126-1-AP; AB_2113330 | 1:400 | UT | Did not test in mouse cells |
| GPR107 | Proteintech 25076-1-AP; AB_2879886 | 1:100 | - | Did not test in mouse cells |
| LAMP1 (hu) | CST # 9091; AB_2687579 | 1:200 | 1:1000 |  |
| LAMP1 (hu) | CST #15665; AB_2798750 | 1:200 | - |  |
| HA | Sigma 11867423001; AB_390918 | 1:400 | - |  |
| HA | CST #2367S; AB_10691311 | - | 1:1000 |  |
| HSP60 | Abcam ab59457; AB_2121285 | 1:400 | UT | Did not test in mouse cells |
| HOOK2 (ms, hu) | Novus NBP2-14986; AB_2548013 | 1:100 | 1:1000 |  |
| NLRP3 (ms) | Adipogen AG-20B-0014-C100; AB_2885199 | - | 1:2000 |  |
| PLEKHA3/FAPP1 (hu, ms) | Novus NBP1-83924; AB_11035656 | 1:100 | 1:1000 | Many nonspecific bands for mouse lysates by WB; long backorder |
| PLEKHA3/FAPP1 (hu) | Proteintech 11292-1-AP; AB_10637846 | - | 1:1000 | Works ok for WB (worse than Novus), does not work for IF |
| PLEKHA8/FAPP2 (hu, ms) | Proteintech 15410-1-AP; AB_2166928 | 1:100 | 1:1000 | Many nonspecific bands for mouse lysates by WB |
| TGN38 (ms) | Novus NBP1-03495; AB_1522533 | 1:100 | UT |  |
| TGN46/TGOLN2 (hu) | Proteintech 13573-1-AP; AB_10597396 | 1:200 | UT |  |
| TMEM115 | Novus NBP1-80898; AB_11002107 | 1:100 | 1:1000 | Beware nonspecific band most prominent by WB; Did not test in mouse cells |
| TMEM165 | Proteintech 20485-1-AP; AB_10695184 | 1:100 | 1:1000 | Did not test in mouse cells |
| VPS35 | Santa Cruz sc-374372; AB_10988942 | 1:200 | UT | Did not test in mouse cells |
| YIPF4 (ms, hu) | Proteintech 15473-1-AP; AB_2217206 | 1:200 | 1:1000 |  |
| **Secondary antibodies and conjugates** |  |  |  |  |
| Anti-mouse 488 | Thermo A-11029 | 1:200 | - |  |
| Anti-mouse 568 | Thermo A-11004 | 1:200 | - |  |
| Anti-mouse 647 | Thermo A-21235 | 1:200 | - |  |
| Anti-rat 488 | Thermo A-11006 | 1:200 |  |  |
| Anti-rat 647 | Thermo A-21247 | 1:200 | - |  |
| Anti-Rabbit 488 | Thermo A-11034 | 1:200 | - |  |
| Anti-Rabbit 568 | Thermo A-11011 | 1:200 | - |  |
| Anti-Rabbit 647 | Thermo A-21244 | 1:200 | - |  |
| Anti-mouse HRP | Bio-Rad 1706516 | - | 1:10,000 |  |
| Anti-rabbit HRP | Bio-Rad 1706515 | - | 1:10,000 |  |
| Anti-Rabbit IRDye® 680RD | Licor 926-68071 | - | 1:5000 |  |
| Anti-Rabbit IRDye® 800CW | Licor 926-32210 |  |  |  |
| Anti-Mouse IRDye® 680CD | Licor 926-68072 | - | 1:5000 |  |
| Anti-Mouse IRDye® 800 CW | Licor 926-32210 | - | 1:5000 |  |
| IRDye® 800CW Streptavidin | Licor 926-32230 | - | 1:5000 |  |
| Streptavidin-HRP | Thermo 21130 | - | 1:5000 |  |
| Phalloidin-488 | Thermo A12379 | 1:200 | - |  |
| Rhodamine Anti-Tubulin | Bio-Rad 12004166; AB_2884950 | - | 1:1000 |  |

## Methods

### Statement of AI use

[ChatGPT3.5](https://chat.openai.com/) was used to revise specific sections of text for clarity, but was never prompted for *de novo* writing. All text was checked for plagiarism with [Grammarly](https://www.grammarly.com/premium) and [EasyBib](https://www.easybib.com/).

### Cloning

Homology directed repair templates for endogenous tagging were synthesized by Twist Bioscience or Genscript. cDNAs were obtained from the 9090 collection or synthesized by Twist Bioscience. Stable expression plasmids were generally made with Gateway technology (Thermo) or with ClonExpress (Vazyme) using the manufacturers’ protocols. Site-directed mutagenesis was carried out using the Quick-Change Site Directed Mutagenesis Kit (New England Biolabs) as per the manufacturer’s instructions. Site directed mutagenesis primers were often designed with NEB base changer (https://nebasechanger.neb.com/). CRISPR/Cas9 guides were cloned following protocols from the Zhang lab. Primers and other small oligos were synthesized by IDT or Genewiz. Sequences for plasmids used in the study can be found in <u>Table S15</u>.

### Cell culture and differentiation

HEK293(T) and THP-1 cells were obtained from ATCC, whereas iBMDMs were a gift from Kate Fitzgerald (UMass Chan Medical School). Cells used in this study were not validated for identity beyond morphology and function, but they were regularly tested for mycoplasma (Lonza #LT07-318). All cells were cultured at 37 °C with 5% CO_2_. HEK293(T) cells and iBMDMs were cultured in DMEM supplemented with GlutaMAX™, Penicillin-Streptomycin, and 10% FBS. 293T cells were cultured without antibiotics for transfection. THP-1 cells were cultured in RPMI supplemented with GlutaMAX™, Penicillin-Streptomycin, and 10% FBS. iBMDMs and THP-1s were swapped into phenol red-free and FBS-free medium for LDH release assays to minimize assay background. HEK293(T) cells and iBMDMs were frozen in 90% DMEM/10% DMSO. THP-1 cells were frozen in 90% FBS/10% DMSO. THP-1 monocytes were differentiated with Phorbol 12-myristate 13-acetate (PMA; 24 h, 25 ng/μL) followed by rest in complete RPMI (24 h) prior to their use in inflammasome assays. HEK293(T) cells were selected with 1-2 μg/mL puromycin; iBMDMs were selected with 5 μg/mL puromycin; THP-1 cells were selected with 2.5 μg/mL puromycin.

### Cell culture chemical compounds

Nigericin (Sigma N7143) was resuspended in ethanol (20 mM) and stored at −20 °C. LPS-B5 (Invivogen tlrl-b5lps) was resuspended in ultrapure water, aliquoted, and stored at −20 °C. Monensin (Cayman Chemicals #16488) was dissolved in ethanol (10 mM) immediately before use. CL097 (Sigma SML2566) was resuspended in ultrapure water (5 mg/mL) with dropwise addition of 35% HCl, aliquoted, and stored at −80 °C. Imiquimod hydrochloride (MedChemExpress HY-B0180A) was resuspended in ultrapure water with sonication (4 mg/mL), aliquoted, and stored at −80 °C. Biotin tyramide (Iris biotech LS-3500) was resuspended in DMSO (500 mM), aliquoted, and stored at −80 °C. PMA (Sigma P1585) was resuspended in DMSO (1 mg/mL), aliquoted, and stored at −80 °C. All other chemical reagents are listed in <u>Table S15</u>.

### Western blotting

Adherent cells were scraped in PBS and pelleted by centrifugation (500 xg, 5 min). Suspension cells were collected by centrifugation (500 xg, 5 min), resuspended in PBS, and pelleted by centrifugation (500 xg, 5 min). Unless otherwise specified, cells were then lysed with RIPA buffer (Thermo 89901) supplemented with protease and phosphatase inhibitors. Each six-well equivalent of cells (or ∼1 M suspension cells) was resuspended in ∼70 µL lysis buffer and placed on ice (1 h). Cell lysates were then centrifuged to pellet insoluble material (13,000 rpm, 10 min, 4°). Protein concentrations of the clarified lysates were determined with the DC protein assay, and equal concentrations of clarified lysate (between 0.5-2 mg/mL) were boiled (except for GPCRs) in 1× LDS buffer with 25 mM DTT. Equal amounts (between 7.5-15 μL) of lysate were run on 26-well 4%–20% Tris Glycine gels (BioRad) and transferred onto nitrocellulose membranes with an iBlot2 (20V, 10 min; Thermo). Membranes were blocked with PBS-Tween20(0.05%)-BSA(3%) (PBST-BSA; 1 h, RT) followed by incubation with the [indicated antibody resuspended in Intercept® (TBS) Blocking Buffer at the indicated dilution](https://harperlab.pubpub.org/pub/nlrp3/release/1#nnrfhqzjgfl) (overnight, 4°C) with gentle rocking. Membranes were washed 3 times with PBST over ∼30 min prior to incubation with the [appropriate secondary antibodies](https://harperlab.pubpub.org/pub/nlrp3/release/1#nnrfhqzjgfl) (1 h, RT), if applicable. Membranes were again washed 3 times with PBST over ∼30 min prior to developing images of blots using Enhanced-Chemiluminescence or fluorescent signal on a BioRad ChemiDoc imager.

### Lentivirus production and THP-1/iBMDM spinfection

For an alternative extended protocol, see https://www.addgene.org/protocols/lentivirus-production. HEK293T cells used to produce virus were always passaged before becoming fully confluent, and were kept in passage for less than 3 weeks. To produce lentivirus, 2.6 x 10^6^ 293T cells were seeded in each well of a 6-well dish in 2 mL antibiotic-free DMEM (targeting ∼60-80% confluence the next day). The next day, transfection complexes were made by mixing 250 ng pMD2.G (Addgene#12259), 750 ng psPAX2 (Addgene#11260), and 1 μg lentiviral vector in OptiMEM (50 μL). In a second tube, Transporter 5® (10 μL) or PEI Max® (1 mg/mL, 10 μL) was added to OptiMEM (50 μL). The tubes were mixed, incubated (20 min, RT), and then added to cells dropwise with gentle mixing. 6-8 h later, wells were exchanged into fresh DMEM without antibiotic. 48 h after transfection, virus was harvested by collecting and centrifuging the cell culture supernatant (1000 xg, 5 min) to remove cell debris. The clarified supernatant was then incubated with Lenti-X™ Concentrator at a 3 medium:1 concentrator volumetric ratio (gentle inversion, overnight, 4°). The concentrated virus was centrifuged (1500 xg, 45 min, 4 °C) and subsequently resuspended in fresh RPMI (for spinfecting THP-1 cells) or DMEM (for spinfecting iBMDMs). Generally, virus was concentrated 20X (e.g., 2 mL of virus-containing supernatant was concentrated to 100 μL) and 100-200 μL of the concentrated virus was then used for spinfection (below). Excess virus was snap frozen and stored at −80 °C for future use.

To spinfect THP-1s, 1 M cells were counted and diluted into 1.8 mL medium in a 6-well. Concentrated virus (200 μL) in RPMI was added, followed by polybrene (1.6 μL, 10 mg/mL). The 6-well plates were wrapped in parafilm and centrifuged (1000 xg, 2 h, 30-37 °C). Following centrifugation, THP-1 cells were resuspended but left in viral medium. The next day, medium was replaced by centrifugation (500 xg, 5 min; 3 mL medium added). The following day, cells were expanded into 10 cm dishes (10 mL medium). Cells were selected/sorted 72 h post-spinfection.

To spinfect iBMDMs, cells were seeded at approximately ∼20% confluence in 6-well dishes. The next day, the medium was replaced with fresh DMEM (1.8 mL). Concentrated virus (200 μL) in DMEM was added, followed by polybrene (1.6 μL, 10 mg/mL). Plates were centrifuged (1000 xg, 2 h, 30-37 °C) then returned to the incubator. The next day, the cells were expanded into 10 cm dishes. Cells were selected/sorted 72 h post-spinfection.

### CRISPR Knockouts

sgRNAs were designed with CHOPCHOP or CRISPick, while other sgRNAs were picked based on data in the literature (<u>Table S15</u>). Oligos containing desired sgRNA sequences were annealed and assembled into the appropriate vector with the Golden Gate cloning protocols provided by the Zhang lab.

Bulk knockouts in THP-1 cells expressing lentiCRISPRv2-puro were selected with puromycin (2.5 μg/mL, 7 d), recovered (3-6 d), and immediately used for experiments. Bulk knockouts in THP-1s expressing lentiCRISPRV2-mCherry were sorted for mCherry-positive cells (using the parental line as a negative control) with a Sony SH800 Cell Sorter operating in purity mode. For each sorted cell line, 100,000 cells were obtained and expanded for experiments within 7 d. Cell pellets for western blots were taken on the same day as cells were plated for inflammasome assays. For clonal iBMDMs spinfected with lentiCRISPRv2-puro vectors, cells were selected with puromycin (5 μg/mL, 7 d) then single-cell cloned into 96-well dishes with a Sony SH800 Cell Sorter operating in single-cell mode. Clonal knockouts were first validated at the genomic level with Miseq using OutKnocker 2.0 [187] to align gDNA reads (see <u>Table S15</u> for genomic primers). Subsequently, knockouts were validated at the protein level by Western Blot or immunofluorescence.

### Endogenous tagging

To endogenously tag EEA1 (mNeonGreen-P2A-3xFLAG-EEA1) and TMEM115 (TMEM115-3xHA-P2A-mNeonGreen), homology directed repair (HDR) templates were designed manually and synthesized by Twist Bioscience. Each HDR template contains ∼500 bp of homology to the genome on each side of the insert and mutations to avoid repeated CRISPR/Cas9 cutting at two different genomic sites. sgRNAs were synthesized with the Precision gRNA Synthesis Kit (Invitrogen A29377) using the manufacturer’s protocols. Purified sgRNAs were aliquoted (1 μL, ∼2 μg/μL) and stored at −80 °C. Sequences for gRNAs and HDR templates are in <u>Table S15</u>.

Electroporation was carried out as described previously (dx.doi.org/10.17504/protocols.io.ewov1qykkgr2/v1). RNP complexes and endotoxin-free HDR templates were electroporated into HEK293T cells with the Neon™ Transfection System and 10 μL Kit (MPK1025). Buffer R (10 μL) was added to a sterile 1.5 mL tube, followed by purified Cas9 protein (6 μg) and sgRNA (1.2 μg). After mixing by gentle pipetting, complexes were incubated (10 min, RT). Meanwhile, HEK293T cells were resuspended in 1 x 10^6^ per 5 μL (2 x 10^8^ cells/mL) in Buffer R. Electrolytic buffer was added to a neon tube (3 mL) and placed in the electroporator. Following incubation, HDR template was added (1 μL, 2000 ng/μL) to the 1.5 mL tube containing the RNP complex, followed by HEK293T cells (10 μL, 2 x 10^8^ cells/mL). Concentrations of the various components were high enough such that the final mixture was between 21-25 μL. Half of the mixture (10 μL) was taken up with the neon pipet carefully to avoid air bubbles, which can cause arching. Cells were electroporated (1150 V, pulse width 20, 2 pulses) and then immediately plated into fresh complete DMEM (2 mL) in a six-well dish. The process was repeated once and the second set of cells were combined into the same well. After recovering (3-5 d), mNeonGreen-positive HEK293T cells were single-cell cloned into 96-well dishes using a Sony SH800 Cell Sorter operating in single-cell mode. Cells were checked for successful incorporation of the endogenous tags by western blot and screened for proper tag localization by immunofluorescence.

### LDH and IL-1β release assays

For extended step-by-step protocols, see dx.doi.org/10.17504/protocols.io.dm6gpzr3plzp/v1.

Confluent iBMDMs were washed with PBS, trypsinized, and neutralized with complete DMEM. Resuspended iBMDMs were counted, and subsequently 200,000 cells were seeded in 96-well dishes (phenol red-free complete DMEM, 200 μL). Each genotype was plated in triplicate for every treatment condition, in addition to a triplicate of cell culture medium as a background control. The next day, cells were carefully exchanged into phenol red- and serum-free DMEM ± LPS (1 μg/mL, 100 μL). 4 h later, LPS-containing medium (1 μg/mL) with activators or lysis buffer was added to the appropriate wells (100 μL, 2X as the indicated concentration on figures). 1 h (nigericin, lysis control) or 2 h (CL097) later, plates were gently tapped, centrifuged (1000 xg, 5 min), and 150 μL of the supernatant was transferred to fresh 96-well plates. Transferred supernatant (10 μL) and cell culture medium (40 μL) were taken for the CyQUANT™ LDH Cytotoxicity Assay, following the manufacturers’ protocol. Transferred supernatant (1 μL) and medium (11.5 μL) were taken for the Lumit® IL-1β Mouse Immunoassay in 384-well plates, following the manufacturers’ protocols.

THP-1 monocytes were counted and then 10,000 cells in complete RPMI (100 μL) were seeded in each well of a 96-well dish. Each genotype was plated in triplicate for every treatment condition, in addition to a triplicate of cell culture medium. Controls for each genotype included an untreated triplicate of cells and a triplicate of cells for maximum LDH release (lysis buffer). RPMI containing PMA was added (100 μL, 50 ng/mL). 24 h later, medium was carefully exchanged into phenol red-fee complete RPMI (200 μL). 24 h later, cells were exchanged into phenol red- and serum-free RPMI ± LPS (1 μg/mL). 3 h later, medium containing activators, lysis control, or LPS was added to the appropriate wells (100 μL, 2X as the indicated concentration on figures). 1-2 h later, plates were tapped, centrifuged (1000 xg, 5 min), and 150 μL of the supernatant was transferred to fresh 96-well plates. Transferred supernatant (50 μL) was taken for the CyQUANT™ LDH Cytotoxicity Assay, following the manufacturers’ protocol. Transferred supernatant (12.5 μL) was taken for the Lumit® IL-1β Human Immunoassay in 384-well plates, following the manufacturers’ protocols.

% maximum LDH release for each replicate was calculated as follows: (Treatment – Average[medium])/(Average[Geneotype-matched maximum lysis control] – Average[medium]) x 100%. IL-1β values were calculated with linear interpolation in GraphPad Prism. For more details and example spreadsheets, see the protocols.io associated with these methods above.

### Immunofluorescence

For an extended step-by-step IF protocol, see dx.doi.org/10.17504/protocols.io.kxygxyeeol8j/v1.

Poly-L-lysine solution (Sigma P8920) was diluted in PBS (0.01%), added (0.5 mL) to 24-well glass bottom plates (Cellvis P24-1.5H-N), and placed in a cell culture incubator (1 – 24 h, 37 °C, 5% CO_2_). Wells were washed 3x with PBS (1 mL) and replaced with complete DMEM (1 mL). Cells were plated such that they would not be more than ∼70% confluent the next day (∼15% for iBMDM, ∼30% for HEK293T). The next day, medium was carefully exchanged with fresh DMEM (1 mL) with or without compounds, as indicated in the figures. For certain experiments, iBMDMs were sometimes first primed with DMEM containing LPS (1 mL, 1 μg/mL, 4 h) prior to the direct addition of compounds to the medium. Treatments were staggered so that all wells were ready to fix at the same time.

Following treatment, medium was aspirated and cells were immediately fixed by adding pre-warmed paraformaldehyde (PFA; Electron Microscopy Sciences 15710) freshly diluted into PBS (4%; 1 mL, 37 °C, 10 min gentle rocking). Triton-X 100 in PBS was added (0.4%; 1 mL) directly to the PFA solution, quickly aspirated, and then cells were permeabilized with triton-X 100 in PBS (0.4%; RT, 10 min with gentle rocking). Cells were washed 3x with Tween-20 in PBS (PBST; 0.02%, RT, 10 min) and then blocked with PBST-Block (PBST supplemented with 3% BSA and 22.52 mg/mL glycine; RT, 0.5 mL, 1 h). Primary antibodies were diluted into PBST-Block at the concentrations indicated in <u>Table S15</u> and incubated overnight with gentle rocking (200 μL, 4 °C). The next day, wells were washed 3x with PBST (1 mL). From here on, all steps were done with aluminum foil wrapping to minimize fluorophore quenching. The appropriate fluorescent secondary antibodies, and sometimes also phalloidin (<u>Table S15</u>), were added (200 μL in PBST-Block, RT, 90 min, gentle rocking). Wells were washed 3x with PBST (1 mL, 10 min) prior to incubation with Hoechst 33342 (5 μM in PBS, 0.5 mL, RT, 10 min). Wells were washed 3x with PBS (RT, 10 min) and kept at RT for confocal imaging within the same day.

Wells were imaged with either a Nikon Ti or Nikon Ti2 equipped with a Yokogawa CSU-X1 spinning disk confocal microscope. Images were collected on an ORCA-Fusion BT sCMOS camera (Hamamatsu) controlled by NIS-Elements software (Nikon, version AR, RRID:SCR_014329). A Nikon Plan Apo ×40/1.30 NA oil objective lens (ASC specks) or ×100/1.40 NA oil objective lens (all other experiments) was used. Relevant laser lines for experiments included a 405nm laser with a ET455/50 emission filter (Chroma) [405 nm], a 488nm laser with a ET525/50 emission filter (Chroma) [488 nm], a 561nm laser with a ET620/60 emissions filter (Chroma) [568 nm], and a 640nm laser with a ET700/75 emission filter (Chroma) [647 nm] controlled by an Acousto-Optic Tunable Filter system. Confocal z-stacks were taken (8 μM, 29 steps) and maximum intensity projection images were calculated with a FIJI [188] script.

When multiple antibodies were used, the control organellar antibody on the 488 channel (e.g., EEA1, Giantin, GOLGA2, Golgin-97, HSP60, YIPF4) was first checked for bleed-through into the 647 channel at the maximum laser intensity and exposure times at least double those used in the multiple antibody experiments. For calculating normalized Golgi intensities (Figure 5F), the Golgi marker (either GOLGA2 or Giantin) was used as a mask, and the total intensities of the Golgi marker and the indicated protein (COPA, COPB, PLEKHA3, PLEKHA8) were extracted with CellProfiler [189]. The ratio of these intensities was calculated and normalized within a set of biological replicates to the average value of the untreated control.

### Whole cell proteomics

HEK293T cells were seeded 1:8 from confluent 6-well plates (2 mL medium). 2 d later, nigericin (2 μL, 20 mM, 30 min) or CL097 (30 μL, 5 mg/mL, 1 h) was added. Following incubation, cells were harvested by scraping cells in ice-cold PBS (1 mL), pelleting cells (1000 xg, 5 min), and resuspending them in lysis buffer (250 µL; 8 M urea, 200 mM EPPS pH=8.5, cOmplete Protease Inhibitor cocktail tablet). Cells were passed through a 21G needle 15 times to shear DNA and complete lysis. Lysates were centrifuged (13,000 RPM, 15 min), and the clarified supernatant was quantified with the DC protein assay Kit II (Bio rad) using a 1:5 dilution in urea-free buffer. 80 µg of protein from each replicate was processed according to the following protocol: dx.doi.org/10.17504/protocols.io.kqdg3x4n1g25/v1

### Organelle-IPs and peptide preparation

For full step-by-step protocols for organelle isolation, including with THP-1 macrophages, see dx.doi.org/10.17504/protocols.io.261ge5p57g47/v1. These protocols synthesize information from a variety of sources [79][82][83][84][87][110] (also: dx.doi.org/10.17504/protocols.io.4r3l24kxxg1y/v2, dx.doi.org/10.17504/protocols.io.6qpvrdjrogmk/v2, and dx.doi.org/10.17504/protocols.io.bybjpskn).

HEK293(T) cells were seeded 1:8 from a confluent 15-cm plate (20 mL medium). 2 d later, nigericin (20 μL, 20 mM, 30 min) or CL097 (300 μL, 5 mg/mL, 1 h) was added. Following incubation with compounds, cells were harvested and processed following the protocol above. Notably, these samples were processed with bead incubation times of 30 min (LysoIP, GolgiIP, MitoIP) or 50 min (EndoIP), rather than our more up-to-date recommendations (protocols.io above). Additionally, CL097-treated cells were labile and thus harvested in their growth medium prior to one wash with PBS.

### APEX2 biotinylation, immunoprecipitation, and peptide preparation

For full step-by-step protocols for APEX2 proximity biotinylation, see the following: dx.doi.org/10.17504/protocols.io.5qpvok55zl4o/v1

For timecourse APEX2 experiments (Figure 6), cells expressing Nlrp3-APEX2 or APEX2-P4C were seeded 1:10 in 10 cm plates (10 mL medium). 2 d later, the medium was replaced with fresh DMEM with LPS (10 mL, 1 μg/mL, 4 h). Nigericin (10 μL, 20 mM; 10, 20, or 30 min) or CL097 (15, 30, 60 min) was added.

Depending on the time point, biotin phenol (500 mM, 10 μL) was added 45 min prior to harvest. Cells were harvested and processed in batches following [this extended protocol](https://www.protocols.io/view/apex2-based-proximity-biotinylation-of-nlrp3-and-p-5qpvok55zl4o/v1).

For single point APEX2 experiments (Figure 7, Negative Data Figure 5), cells expressing Nlrp3-APEX2 or APEX2-P4C were seeded 1:10 in 10 cm plates (10 mL medium). 2 d later, medium was replaced with fresh DMEM with or without LPS (10 mL, 1 μg/mL, 4 h). Nigericin (10 μL, 20 mM) or CL097 was added (150 μL, 5 mg/mL). Depending on the time point, biotin phenol (500 mM, 10 μL) was added 45 min prior to harvest. Cells were harvested and processed in batches following [this extended protocol](https://www.protocols.io/view/apex2-based-proximity-biotinylation-of-nlrp3-and-p-5qpvok55zl4o/v1).

### Quantitative proteomics

Mass spectrometry data were acquired as summarized in <u>Table S15</u>. Database searching included all *H. sapiens* (2020-02-25) or *M. musculus* (2022-1-26) entries from UniProt. Transgenes (e.g., Nlrp3-APEX2 or APEX2-P4C) were appended to these files whenever applicable. The databases were concatenated with one composed of all protein sequences in that database in reversed order (decoys). Peptides were identified with Comet searches [190][191] (https://uwpr.github.io/Comet/). For hrMS2-FAIMS, searches were performed using a 50-ppm precursor ion tolerance for total protein level profiling, and the product ion tolerance was set to 0.02 Da. TMTpro labels on lysine residues and peptide N termini (+304.207 Da) and carbamidomethylation of cysteine residues (+57.021 Da) were set as static modifications, and oxidation of methionine residues (+15.995 Da) was set as a variable modification. Peptides identifications were filtered with linear discriminate analysis followed by additional filtering to achieve a ∼2% final protein FDR (using the relevant target and decoy databases) [192] [193][194][195][196].

Peptide identifications were filtered for summed signal-to-noise ratio (SNR > 200) across the TMTplex and for precursor signals that contained an isolation purity of >0.5 of the MS1 isolation window. For each protein, the filtered peptide–spectrum match TMTpro raw intensities were summed and Log_2_ normalized to create protein quantification values (weighted average). For protein TMT quantifications, peptides across TMT channels were summed and normalized using MSstats [151] with either Tukey’s median polish (proximity labelling datasets) or MSstats (all other datasets) statistical implementations.

### Data analysis

Code can be found at https://github.com/priyaveeraraghavan/inflame.

Time:Bait interaction model (Figure 6G): For each treatment type (nigericin or CL097), gene intensities were jointly normalized across the datasets (NLRP3 or P4C bait) using MSStats. Genes with non-zero values in at least 3 samples in both datasets were retained for further analysis. The normalized gene intensities were fit using a linear model with the formula ∼time+bait+time:bait using lm() in R. Coefficient p-values were adjusted for false discovery rate (FDR) using the Benjamini-Hochberg approach (q-value).

APEX2 timecourse single-dataset clustering (Figure 6—Supporting Figures 3-6): Each APEX2 timecourse dataset was filtered for genes that significantly changed across conditions (ANOVA q < 0.05) and for detection in all APEX2 timecourse datasets. Genes were then clustered based on their Log_2_ fold changes at each timepoint with respect to t=0 using the Leiden algorithm. These clusters were visualized on a UMAP embedding of the data.

APEX2 timecourse multi-dataset cluster integration (Figure 6E-F): Gene cluster similarities were determined by computing the Jaccard index between two clusters in different datasets. Clusters with similarity above an empirically determined threshold were considered “connected”, and each connected component of clusters is defined as a meta_cluster.

Gene group organelle enrichment: Whenever specified, we used a mapping of genes to organelles from the graph-based annotations Hein*, Peng*, Todorova*, McCarthy*, Kim*, and Liu* *et al* [84]. Otherwise, in-house organellar annotations were used. For Enrichment of organelle genes in each gene group was computed using Fisher’s exact test. For enrichment analysis of COPI/COPII cargo proteins, annotations reported in Adolf* and Rhiel* *et al.* [197] were used. For enrichment of proteins on vesicles captured by GCC1 (GCC88) and GOLGA1 (Golgin-97), annotations reported in Shin *et al.* [132] were used.

## Competing interests

J.W.H. is a consultant and founder of Caraway Therapeutics (a wholly owned subsidiary of Merck & Co, Inc) and is a member of the scientific advisory board for Lyterian Therapeutics. P.V. completed this work as a volunteer outside the scope of her employment at Dyno Therapeutics. L.R.H and J.A.P. declare no competing interest.

## Acknowledgements

We thank all members of the laboratory of J.W.H. for feedback. We especially thank lab members Ian Smith for analysis scripts and computational feedback; Frances Hundley for organelleIP training and feedback; Miguel Gonzalez-Lozano, Ian Smith, Sharan Swarup, and Alex Panov for mass spectrometry advice; Felix Kraus and Melissa Hoyer for imaging advice; Chan Lee, Tao Fu, and Frances Hundley for daily discussions; Enya Miguel Whelan, Myzi Ormenaj, and Julia Paoli for general laboratory support. We thank Gyan Prakash (Churchman Lab, Harvard Medical School) for virus encoding the MitoTag and MitoControlTag. We thank Liudmila Andreeva (University Tübingen), Liron David (Ben-Gurion University), Romeo Ricci (Laboratoire de biochimie et de biologie moléculaire), Antonella De Matteis and Rossella Venditti (Telethon Institute of Genetics and Medicine), Emma McKay (Queen’s University Belfast), and John Pulice (Harvard Medical School) for critical discussions. We thank Megan Hochstrasser (Arcadia Science) for extensive pubpub platform advice and custom CSS for pubpub figure captions. We thank Kate Fitzgerald (UMass Chan Medical School) for WT and Nlrp3-knockout iBMDMs. We acknowledge the Nikon Imaging Center (Now called the Core for Imaging Technology & Education, Harvard Medical School) for imaging assistance. This work was supported by NIH grants R01NS083524 to J.W.H and R01GM132129 to J.A.P. We apologize for any missing scientific citations due to oversights or the limitations of our knowledge, and we will add missing references as necessary.

## Author contributions

L.R.H. conceived the study with help from J.W.H. L.R.H. performed all experiments, with proteomics support from J.A.P. L.R.H and P.V. analyzed the data. P.V. built the Shiny app. The paper was written by L.R.H. with input from all authors.

## Peer review

Peer review, solicited and edited by Isabella Rauch, will appear here. We invite the community to comment on our work below or on specific passages through in-line comments, [as explained in this video](https://www.youtube.com/watch?v=f90SKxJOcRY).

## Notes

https://harperlab.pubpub.org/pub/nlrp3/

## References

1. Vande Walle L, Lamkanfi M. (2023). Drugging the NLRP3 inflammasome: from signalling mechanisms to therapeutic targets. 10.1038/s41573-023-00822-2

2. Remick BC, Gaidt MM, Vance RE. (2023). Effector-Triggered Immunity. 10.1146/annurev-immunol-101721-031732

3. 3. Brubaker SW, Bonham KS, Zanoni I, Kagan JC. (2015). Innate Immune Pattern Recognition: A Cell Biological Perspective. 10.1146/annurev-immunol-032414-112240

4. Barnett KC, Li S, Liang K, Ting JP-Y. (2023). A 360° view of the inflammasome: Mechanisms of activation, cell death, and diseases. 10.1016/j.cell.2023.04.025

5. Liston A, Masters SL. (2017). Homeostasis-altering molecular processes as mechanisms of inflammasome activation. 10.1038/nri.2016.151↩

6. Martinon F, Burns K, Tschopp J. (2002). The Inflammasome. 10.1016/s1097-2765(02)00599-3

7. Srinivasula SM, Poyet J-L, Razmara M, Datta P, Zhang Z, Alnemri ES. (2002). The PYRIN-CARD Protein ASC Is an Activating Adaptor for Caspase-1. 10.1074/jbc.c200179200

8. Kayagaki N, Stowe IB, Lee BL, O’Rourke K, Anderson K, Warming S, Cuellar T, Haley B, Roose-Girma M, Phung QT, Liu PS, Lill JR, Li H, Wu J, Kummerfeld S, Zhang J, Lee WP, Snipas SJ, Salvesen GS, Morris LX, Fitzgerald L, Zhang Y, Bertram EM, Goodnow CC, Dixit VM. (2015). Caspase-11 cleaves gasdermin D for non-canonical inflammasome signalling. 10.1038/nature15541

9. He W, Wan H, Hu L, Chen P, Wang X, Huang Z, Yang Z-H, Zhong C-Q, Han J. (2015). Gasdermin D is an executor of pyroptosis and required for interleukin-1β secretion. 10.1038/cr.2015.139

10. Shi J, Zhao Y, Wang K, Shi X, Wang Y, Huang H, Zhuang Y, Cai T, Wang F, Shao F. (2015). Cleavage of GSDMD by inflammatory caspases determines pyroptotic cell death. 10.1038/nature15514

11. Thornberry NA, Bull HG, Calaycay JR, Chapman KT, Howard AD, Kostura MJ, Miller DK, Molineaux SM, Weidner JR, Aunins J, Elliston KO, Ayala JM, Casano FJ, Chin J, Ding GJ-F, Egger LA, Gaffney EP, Limjuco G, Palyha OC, Raju SM, Rolando AM, Salley JP, Yamin T-T, Lee TD, Shively JE, MacCross M, Mumford RA, Schmidt JA, Tocci MJ. (1992). A novel heterodimeric cysteine protease is required for interleukin-1βprocessing in monocytes. 10.1038/356768a0

12. Cerretti DP, Kozlosky CJ, Mosley B, Nelson N, Van Ness K, Greenstreet TA, March CJ, Kronheim SR, Druck T, Cannizzaro LA, Huebner K, Black RA. (1992). Molecular Cloning of the Interleukin-1β Converting Enzyme. 10.1126/science.1373520

13. Liu X, Zhang Z, Ruan J, Pan Y, Magupalli VG, Wu H, Lieberman J. (2016). Inflammasome-activated gasdermin D causes pyroptosis by forming membrane pores. 10.1038/nature18629

14. Ding J, Wang K, Liu W, She Y, Sun Q, Shi J, Sun H, Wang D-C, Shao F. (2016). Pore-forming activity and structural autoinhibition of the gasdermin family. 10.1038/nature18590

15. Shi J, Gao W, Shao F. (2017). Pyroptosis: Gasdermin-Mediated Programmed Necrotic Cell Death. 10.1016/j.tibs.2016.10.004

16. Kayagaki N, Kornfeld OS, Lee BL, Stowe IB, O’Rourke K, Li Q, Sandoval W, Yan D, Kang J, Xu M, Zhang J, Lee WP, McKenzie BS, Ulas G, Payandeh J, Roose-Girma M, Modrusan Z, Reja R, Sagolla M, Webster JD, Cho V, Andrews TD, Morris LX, Miosge LA, Goodnow CC, Bertram EM, Dixit VM. (2021). NINJ1 mediates plasma membrane rupture during lytic cell death. 10.1038/s41586-021-03218-7

17. Phulphagar K, Kühn LI, Ebner S, Frauenstein A, Swietlik JJ, Rieckmann J, Meissner F. (2021). Proteomics reveals distinct mechanisms regulating the release of cytokines and alarmins during pyroptosis. 10.1016/j.celrep.2021.108826

18. Wei B, Billman ZP, Nozaki K, Goodridge HS, Miao EA. (2024). NLRP3, NLRP6, and NLRP12 are inflammasomes with distinct expression patterns. 10.1101/2024.02.05.579000

19. Bauernfeind FG, Horvath G, Stutz A, Alnemri ES, MacDonald K, Speert D, Fernandes-Alnemri T, Wu J, Monks BG, Fitzgerald KA, Hornung V, Latz E. (2009). Cutting Edge: NF-κB Activating Pattern Recognition and Cytokine Receptors License NLRP3 Inflammasome Activation by Regulating NLRP3 Expression. 10.4049/jimmunol.0901363

20. Koller BH, Nguyen M, Snouwaert JN, Gabel CA, Ting JP-Y. (2024). Species-specific NLRP3 regulation and its role in CNS autoinflammatory diseases. 10.1016/j.celrep.2024.113852

21. Mulvey CM, Breckels LM, Crook OM, Sanders DJ, Ribeiro ALR, Geladaki A, Christoforou A, Britovšek NK, Hurrell T, Deery MJ, Gatto L, Smith AM, Lilley KS. (2021). Spatiotemporal proteomic profiling of the pro-inflammatory response to lipopolysaccharide in the THP-1 human leukaemia cell line. 10.1038/s41467-021-26000-9

22. Gritsenko A, Yu S, Martin-Sanchez F, Diaz-del-Olmo I, Nichols E-M, Davis DM, Brough D, Lopez-Castejon G. (2020). Priming Is Dispensable for NLRP3 Inflammasome Activation in Human Monocytes In Vitro. 10.3389/fimmu.2020.565924

23. Juliana C, Fernandes-Alnemri T, Kang S, Farias A, Qin F, Alnemri ES. (2012). Non-transcriptional Priming and Deubiquitination Regulate NLRP3 Inflammasome Activation. 10.1074/jbc.m112.407130

24. Seoane PI, Lee B, Hoyle C, Yu S, Lopez-Castejon G, Lowe M, Brough D. (2020). The NLRP3–inflammasome as a sensor of organelle dysfunction. 10.1083/jcb.202006194

25. McKee CM, Coll RC. (2020). NLRP3 inflammasome priming: A riddle wrapped in a mystery inside an enigma. 10.1002/jlb.3mr0720-513r

26. Tapia-Abellán A, Angosto-Bazarra D, Martínez-Banaclocha H, de Torre-Minguela C, Cerón-Carrasco JP, Pérez-Sánchez H, Arostegui JI, Pelegrin P. (2019). MCC950 closes the active conformation of NLRP3 to an inactive state. 10.1038/s41589-019-0278-6

27. Cosson C, Riou R, Patoli D, Niu T, Rey A, Groslambert M, De Rosny C, Chatre E, Allatif O, Henry T, Venet F, Milhavet F, Boursier G, Belot A, Jamilloux Y, Merlin E, Duquesne A, Grateau G, Savey L, Jacques Maria AT, Pagnier A, Poutrel S, Lambotte O, Mallebranche C, Ardois S, Richer O, Lemelle I, Rieux-Laucat F, Bader-Meunier B, Amoura Z, Melki I, Cuisset L, Touitou I, Geyer M, Georgin-Lavialle S, Py BF. (2024). Functional diversity of NLRP3 gain-of-function mutants associated with CAPS autoinflammation. 10.1084/jem.20231200

28. Welzel T, Kuemmerle-Deschner JB. (2021). Diagnosis and Management of the Cryopyrin-Associated Periodic Syndromes (CAPS): What Do We Know Today? 10.3390/jcm10010128

29. He Y, Zeng MY, Yang D, Motro B, Núñez G. (2016). NEK7 is an essential mediator of NLRP3 activation downstream of potassium efflux. 10.1038/nature16959

30. Schmid-Burgk JL, Chauhan D, Schmidt T, Ebert TS, Reinhardt J, Endl E, Hornung V. (2016). A Genome-wide CRISPR (Clustered Regularly Interspaced Short Palindromic Repeats) Screen Identifies NEK7 as an Essential Component of NLRP3 Inflammasome Activation. 10.1074/jbc.c115.700492

31. Shi H, Wang Y, Li X, Zhan X, Tang M, Fina M, Su L, Pratt D, Bu CH, Hildebrand S, Lyon S, Scott L, Quan J, Sun Q, Russell J, Arnett S, Jurek P, Chen D, Kravchenko VV, Mathison JC, Moresco EMY, Monson NL, Ulevitch RJ, Beutler B. (2015). NLRP3 activation and mitosis are mutually exclusive events coordinated by NEK7, a new inflammasome component. 10.1038/ni.3333

32. Sharif H, Wang L, Wang WL, Magupalli VG, Andreeva L, Qiao Q, Hauenstein AV, Wu Z, Núñez G, Mao Y, Wu H. (2019). Structural mechanism for NEK7-licensed activation of NLRP3 inflammasome. 10.1038/s41586-019-1295-z

33. Yissachar N, Salem H, Tennenbaum T, Motro B. (2006). Nek7 kinase is enriched at the centrosome, and is required for proper spindle assembly and mitotic progression. 10.1016/j.febslet.2006.10.069

34. Wei J-H, Seemann J. (2017). Golgi ribbon disassembly during mitosis, differentiation and disease progression. 10.1016/j.ceb.2017.03.008

35. Fielding AB, Royle SJ. (2013). Mitotic inhibition of clathrin-mediated endocytosis. 10.1007/s00018-012-1250-8

36. Schmacke NA, O’Duill F, Gaidt MM, Szymanska I, Kamper JM, Schmid-Burgk JL, Mädler SC, Mackens-Kiani T, Kozaki T, Chauhan D, Nagl D, Stafford CA, Harz H, Fröhlich AL, Pinci F, Ginhoux F, Beckmann R, Mann M, Leonhardt H, Hornung V. (2022). IKKβ primes inflammasome formation by recruiting NLRP3 to the trans-Golgi network. 10.1016/j.immuni.2022.10.021

37. Andreeva L, David L, Rawson S, Shen C, Pasricha T, Pelegrin P, Wu H. (2021). NLRP3 cages revealed by full-length mouse NLRP3 structure control pathway activation. 10.1016/j.cell.2021.11.011↩

38. Hochheiser IV, Pilsl M, Hagelueken G, Moecking J, Marleaux M, Brinkschulte R, Latz E, Engel C, Geyer M. (2022). Structure of the NLRP3 decamer bound to the cytokine release inhibitor CRID3. 10.1038/s41586-022-04467-w

39. Yu X, Matico RE, Miller R, Chauhan D, Van Schoubroeck B, Grauwen K, Suarez J, Pietrak B, Haloi N, Yin Y, Tresadern GJ, Perez-Benito L, Lindahl E, Bottelbergs A, Oehlrich D, Van Opdenbosch N, Sharma S. (2024). Structural basis for the oligomerization-facilitated NLRP3 activation. 10.1038/s41467-024-45396-8

40. Akbal A, Dernst A, Lovotti M, Mangan MSJ, McManus RM, Latz E. (2022). How location and cellular signaling combine to activate the NLRP3 inflammasome. 10.1038/s41423-022-00922-w

41. D P, CA G. (1994). Interleukin-1 beta maturation and release in response to ATP and nigericin. Evidence that potassium depletion mediated by these agents is a necessary and common feature of their activity

42. Muñoz-Planillo R, Kuffa P, Martínez-Colón G, Smith BL, Rajendiran TM, Núñez G. (2013). K+ Efflux Is the Common Trigger of NLRP3 Inflammasome Activation by Bacterial Toxins and Particulate Matter. 10.1016/j.immuni.2013.05.016

43. Pétrilli V, Papin S, Dostert C, Mayor A, Martinon F, Tschopp J. (2007). Activation of the NALP3 inflammasome is triggered by low intracellular potassium concentration. 10.1038/sj.cdd.4402195

44. Mangan MSJ, Akbal A, Rivara S, Horvath G, Walch P, Hegermann J, Kaiser R, Duthie F, Selkrig J, Gerhard R, Wachten D, Latz E. (2024). Cytosolic sodium accumulation is a cellular danger signal triggering endocytic dysfunction and NLRP3 inflammasome activation. 10.1101/2024.03.26.586774

45. Martinon F, Pétrilli V, Mayor A, Tardivel A, Tschopp J. (2006). Gout-associated uric acid crystals activate the NALP3 inflammasome. 10.1038/nature04516

46. Rajamäki K, Lappalainen J, Öörni K, Välimäki E, Matikainen S, Kovanen PT, Eklund KK. (2010). Cholesterol Crystals Activate the NLRP3 Inflammasome in Human Macrophages: A Novel Link between Cholesterol Metabolism and Inflammation. 10.1371/journal.pone.0011765

47. Duewell P, Kono H, Rayner KJ, Sirois CM, Vladimer G, Bauernfeind FG, Abela GS, Franchi L, Nuñez G, Schnurr M, Espevik T, Lien E, Fitzgerald KA, Rock KL, Moore KJ, Wright SD, Hornung V, Latz E. (2010). NLRP3 inflammasomes are required for atherogenesis and activated by cholesterol crystals. 10.1038/nature08938

48. Hornung V, Bauernfeind F, Halle A, Samstad EO, Kono H, Rock KL, Fitzgerald KA, Latz E. (2008). Silica crystals and aluminum salts activate the NALP3 inflammasome through phagosomal destabilization. 10.1038/ni.1631

49. Dostert C, Pétrilli V, Van Bruggen R, Steele C, Mossman BT, Tschopp J. (2008). Innate Immune Activation Through Nalp3 Inflammasome Sensing of Asbestos and Silica. 10.1126/science.1156995

50. Halle A, Hornung V, Petzold GC, Stewart CR, Monks BG, Reinheckel T, Fitzgerald KA, Latz E, Moore KJ, Golenbock DT. (2008). The NALP3 inflammasome is involved in the innate immune response to amyloid-β. 10.1038/ni.1636

51. Ising C, Venegas C, Zhang S, Scheiblich H, Schmidt SV, Vieira-Saecker A, Schwartz S, Albasset S, McManus RM, Tejera D, Griep A, Santarelli F, Brosseron F, Opitz S, Stunden J, Merten M, Kayed R, Golenbock DT, Blum D, Latz E, Buée L, Heneka MT. (2019). NLRP3 inflammasome activation drives tau pathology. 10.1038/s41586-019-1769-z

52. Codolo G, Plotegher N, Pozzobon T, Brucale M, Tessari I, Bubacco L, de Bernard M. (2013). Triggering of Inflammasome by Aggregated α–Synuclein, an Inflammatory Response in Synucleinopathies. 10.1371/journal.pone.0055375

53. Mariathasan S, Weiss DS, Newton K, McBride J, O’Rourke K, Roose-Girma M, Lee WP, Weinrauch Y, Monack DM, Dixit VM. (2006). Cryopyrin activates the inflammasome in response to toxins and ATP. 10.1038/nature04515

54. Kanneganti T-D, Özören N, Body-Malapel M, Amer A, Park J-H, Franchi L, Whitfield J, Barchet W, Colonna M, Vandenabeele P, Bertin J, Coyle A, Grant EP, Akira S, Núñez G. (2006). Bacterial RNA and small antiviral compounds activate caspase-1 through cryopyrin/Nalp3. 10.1038/nature04517

55. Groß CJ, Mishra R, Schneider KS, Médard G, Wettmarshausen J, Dittlein DC, Shi H, Gorka O, Koenig P-A, Fromm S, Magnani G, Ćiković T, Hartjes L, Smollich J, Robertson AAB, Cooper MA, Schmidt-Supprian M, Schuster M, Schroder K, Broz P, Traidl-Hoffmann C, Beutler B, Kuster B, Ruland J, Schneider S, Perocchi F, Groß O. (2016). K + Efflux-Independent NLRP3 Inflammasome Activation by Small Molecules Targeting Mitochondria. 10.1016/j.immuni.2016.08.010

56. Billingham LK, Stoolman JS, Vasan K, Rodriguez AE, Poor TA, Szibor M, Jacobs HT, Reczek CR, Rashidi A, Zhang P, Miska J, Chandel NS. (2022). Mitochondrial electron transport chain is necessary for NLRP3 inflammasome activation. 10.1038/s41590-022-01185-3

57. Chen J, Chen ZJ. (2018). PtdIns4P on dispersed trans-Golgi network mediates NLRP3 inflammasome activation. 10.1038/s41586-018-0761-3

58. Zhang Z, Venditti R, Ran L, Liu Z, Vivot K, Schürmann A, Bonifacino JS, De Matteis MA, Ricci R. (2022). Distinct changes in endosomal composition promote NLRP3 inflammasome activation. 10.1038/s41590-022-01355-3

59. Lee B, Hoyle C, Wellens R, Green JP, Martin-Sanchez F, Williams DM, Matchett BJ, Seoane PI, Bennett H, Adamson A, Lopez-Castejon G, Lowe M, Brough D. (2023). Disruptions in endocytic traffic contribute to the activation of the NLRP3 inflammasome. 10.1126/scisignal.abm7134

60. Dong R, Saheki Y, Swarup S, Lucast L, Harper JW, De Camilli P. (2016). Endosome-ER Contacts Control Actin Nucleation and Retromer Function through VAP-Dependent Regulation of PI4P. 10.1016/j.cell.2016.06.037

61. Balla A, Tuymetova G, Barshishat M, Geiszt M, Balla T. (2002). Characterization of Type II Phosphatidylinositol 4-Kinase Isoforms Reveals Association of the Enzymes with Endosomal Vesicular Compartments. 10.1074/jbc.m111807200

62. Wong K, Meyers R, Cantley LC. (1997). Subcellular Locations of Phosphatidylinositol 4-Kinase Isoforms. 10.1074/jbc.272.20.13236

63. Phillips MJ, Voeltz GK. (2015). Structure and function of ER membrane contact sites with other organelles. 10.1038/nrm.2015.8

64. Guillén-Samander A, De Camilli P. (2022). Endoplasmic Reticulum Membrane Contact Sites, Lipid Transport, and Neurodegeneration. 10.1101/cshperspect.a041257

65. Mesmin B, Bigay J, Moser von Filseck J, Lacas-Gervais S, Drin G, Antonny B. (2013). A Four-Step Cycle Driven by PI(4)P Hydrolysis Directs Sterol/PI(4)P Exchange by the ER-Golgi Tether OSBP. 10.1016/j.cell.2013.09.056

66. de Saint-Jean M, Delfosse V, Douguet D, Chicanne G, Payrastre B, Bourguet W, Antonny B, Drin G. (2011). Osh4p exchanges sterols for phosphatidylinositol 4-phosphate between lipid bilayers. 10.1083/jcb.201104062

67. Zewe JP, Wills RC, Sangappa S, Goulden BD, Hammond GR. (2018). SAC1 degrades its lipid substrate PtdIns4P in the endoplasmic reticulum to maintain a steep chemical gradient with donor membranes. 10.7554/elife.35588

68. Venditti R, Masone MC, Rega LR, Di Tullio G, Santoro M, Polishchuk E, Serrano IC, Olkkonen VM, Harada A, Medina DL, La Montagna R, De Matteis MA. (2019). The activity of Sac1 across ER–TGN contact sites requires the four-phosphate-adaptor-protein-1. 10.1083/jcb.201812021

69. Guo C, Chi Z, Jiang D, Xu T, Yu W, Wang Z, Chen S, Zhang L, Liu Q, Guo X, Zhang X, Li W, Lu L, Wu Y, Song B-L, Wang D. (2018). Cholesterol Homeostatic Regulator SCAP-SREBP2 Integrates NLRP3 Inflammasome Activation and Cholesterol Biosynthetic Signaling in Macrophages. 10.1016/j.immuni.2018.08.021

70. Williams DM, Peden AA. (2023). S-acylation of NLRP3 provides a nigericin sensitive gating mechanism that controls access to the Golgi. 10.1101/2023.11.14.566891

71. Yu T, Hou D, Zhao J, Lu X, Greentree WK, Zhao Q, Yang M, Conde D-G, Linder ME, Lin H. (2024). NLRP3 Cys126 palmitoylation by ZDHHC7 promotes inflammasome activation. 10.1016/j.celrep.2024.114070

72. Wang L, Cai J, Zhao X, Ma L, Zeng P, Zhou Lingli, Liu Y, Yang S, Cai Z, Zhang S, Zhou Liang, Yang J, Liu T, Jin S, Cui J. (2023). Palmitoylation prevents sustained inflammation by limiting NLRP3 inflammasome activation through chaperone-mediated autophagy. 10.1016/j.molcel.2022.12.002

73. Hammond GRV, Ricci MMC, Weckerly CC, Wills RC. (2022). An update on genetically encoded lipid biosensors. 10.1091/mbc.e21-07-0363

74. Liu Y, Zhai H, Alemayehu H, Boulanger J, Hopkins LJ, Borgeaud AC, Heroven C, Howe JD, Leigh KE, Bryant CE, Modis Y. (2023). Cryo-electron tomography of NLRP3-activated ASC complexes reveals organelle co-localization. 10.1038/s41467-023-43180-8

75. Donzelli M, Draetta GF. (2003). Regulating mammalian checkpoints through Cdc25 inactivation. 10.1038/sj.embor.embor887

76. Marescal O, Cheeseman IM. (2020). Cellular Mechanisms and Regulation of Quiescence. 10.1016/j.devcel.2020.09.029

77. Tsitsiridis G, Steinkamp R, Giurgiu M, Brauner B, Fobo G, Frishman G, Montrone C, Ruepp A. (2022). CORUM: the comprehensive resource of mammalian protein complexes–2022. 10.1093/nar/gkac1015

78. Huttlin EL, Bruckner RJ, Navarrete-Perea J, Cannon JR, Baltier K, Gebreab F, Gygi MP, Thornock A, Zarraga G, Tam S, Szpyt J, Gassaway BM, Panov A, Parzen H, Fu S, Golbazi A, Maenpaa E, Stricker K, Guha Thakurta S, Zhang T, Rad R, Pan J, Nusinow DP, Paulo JA, Schweppe DK, Vaites LP, Harper JW, Gygi SP. (2021). Dual proteome-scale networks reveal cell-specific remodeling of the human interactome. 10.1016/j.cell.2021.04.011

79. Chen WW, Freinkman E, Wang T, Birsoy K, Sabatini DM. (2016). Absolute Quantification of Matrix Metabolites Reveals the Dynamics of Mitochondrial Metabolism. 10.1016/j.cell.2016.07.040

80. Ray GJ, Boydston EA, Shortt E, Wyant GA, Lourido S, Chen WW, Sabatini DM. (2020). A PEROXO-Tag Enables Rapid Isolation of Peroxisomes from Human Cells. 10.1016/j.isci.2020.101109

81. Chantranupong L, Saulnier JL, Wang W, Jones DR, Pacold ME, Sabatini BL. (2020). Rapid purification and metabolomic profiling of synaptic vesicles from mammalian brain. 10.7554/elife.59699

82. Park H, Hundley FV, Yu Q, Overmyer KA, Brademan DR, Serrano L, Paulo JA, Paoli JC, Swarup S, Coon JJ, Gygi SP, Wade Harper J. (2022). Spatial snapshots of amyloid precursor protein intramembrane processing via early endosome proteomics. 10.1038/s41467-022-33881-x

83. Fasimoye R, Dong W, Nirujogi RS, Rawat ES, Iguchi M, Nyame K, Phung TK, Bagnoli E, Prescott AR, Alessi DR, Abu-Remaileh M. (2023). Golgi-IP, a tool for multimodal analysis of Golgi molecular content. 10.1073/pnas.2219953120

84. Hein MY, Peng D, Todorova V, McCarthy F, Kim K, Liu C, Savy L, Januel C, Baltazar-Nunez R, Bax S, Vaid S, Vangipuram M, Ivanov IE, Byrum JR, Pradeep S, Gonzalez CG, Aniseia Y, Wang E, Creery JS, McMorrow AH, Burgess J, Sunshine S, Yeung-Levy S, DeFelice BC, Mehta SB, Itzhak DN, Elias JE, Leonetti MD. (2023). Global organelle profiling reveals subcellular localization and remodeling at proteome scale. 10.1101/2023.12.18.572249

85. Bond C, Hugelier S, Xing J, Sorokina EM, Lakadamyali M. (2024). Multiplexed DNA-PAINT Imaging of the Heterogeneity of Late Endosome/Lysosome Protein Composition. 10.1101/2024.03.18.585634

86. Hornung V, Latz E. (2010). Critical functions of priming and lysosomal damage for NLRP3 activation. 10.1002/eji.200940185

87. Abu-Remaileh M, Wyant GA, Kim C, Laqtom NN, Abbasi M, Chan SH, Freinkman E, Sabatini DM. (2017). Lysosomal metabolomics reveals V-ATPase- and mTOR-dependent regulation of amino acid efflux from lysosomes. 10.1126/science.aan6298

88. Eapen VV, Swarup S, Hoyer MJ, Paulo JA, Harper JW. (2021). Quantitative proteomics reveals the selectivity of ubiquitin-binding autophagy receptors in the turnover of damaged lysosomes by lysophagy. 10.7554/elife.72328

89. Thomas PD, Ebert D, Muruganujan A, Mushayahama T, Albou L, Mi H. (2021). PANTHER: Making genome-scale phylogenetics accessible to all. 10.1002/pro.4218

90. Laflamme C, McKeever PM, Kumar R, Schwartz J, Kolahdouzan M, Chen CX, You Z, Benaliouad F, Gileadi O, McBride HM, Durcan TM, Edwards AM, Healy LM, Robertson J, McPherson PS. (2019). Implementation of an antibody characterization procedure and application to the major ALS/FTD disease gene C9ORF72. 10.7554/elife.48363

91. Amick J, Roczniak-Ferguson A, Ferguson SM. (2016). C9orf72 binds SMCR8, localizes to lysosomes, and regulates mTORC1 signaling. 10.1091/mbc.e16-01-0003

92. Balendra R, Isaacs AM. (2018). C9orf72-mediated ALS and FTD: multiple pathways to disease. 10.1038/s41582-018-0047-2

93. Banerjee P, Mehta AR, Nirujogi RS, Cooper J, James OG, Nanda J, Longden J, Burr K, McDade K, Salzinger A, Paza E, Newton J, Story D, Pal S, Smith C, Alessi DR, Selvaraj BT, Priller J, Chandran S. (2023). Cell-autonomous immune dysfunction driven by disrupted autophagy in C9orf72 - ALS iPSC-derived microglia contributes to neurodegeneration. 10.1126/sciadv.abq0651

94. Trageser KJ, Yang E-J, Smith C, Iban-Arias R, Oguchi T, Sebastian-Valverde M, Iqbal UH, Wu H, Estill M, Al Rahim M, Raval U, Herman FJ, Zhang YJ, Petrucelli L, Pasinetti GM. (2023). Inflammasome-Mediated Neuronal-Microglial Crosstalk: a Therapeutic Substrate for the Familial C9orf72 Variant of Frontotemporal Dementia/Amyotrophic Lateral Sclerosis. 10.1007/s12035-023-03315-w

95. Jadot M, Colmant C, Wattiaux-De Coninck S, Wattiaux R. (1984). Intralysosomal hydrolysis of glycyl-l-phenylalanine 2-naphthylamide. 10.1042/bj2190965

96. Thiele DL, Lipsky PE. (1990). Mechanism of L-leucyl-L-leucine methyl ester-mediated killing of cytotoxic lymphocytes: dependence on a lysosomal thiol protease, dipeptidyl peptidase I, that is enriched in these cells. 10.1073/pnas.87.1.83

97. Jia J, Wang F, Bhujabal Z, Peters R, Mudd M, Duque T, Allers L, Javed R, Salemi M, Behrends C, Phinney B, Johansen T, Deretic V. (2022). Stress granules and mTOR are regulated by membrane atg8ylation during lysosomal damage. 10.1083/jcb.202207091

98. Lee C, Lamech L, Johns E, Overholtzer M. (2020). Selective Lysosome Membrane Turnover Is Induced by Nutrient Starvation. 10.1016/j.devcel.2020.08.008

99. Jacquin E, Leclerc-Mercier S, Judon C, Blanchard E, Fraitag S, Florey O. (2017). Pharmacological modulators of autophagy activate a parallel noncanonical pathway driving unconventional LC3 lipidation. 10.1080/15548627.2017.1287653

100. Xu Y, Cheng S, Zeng H, Zhou P, Ma Y, Li L, Liu X, Shao F, Ding J. (2022). ARF GTPases activate Salmonella effector SopF to ADP-ribosylate host V-ATPase and inhibit endomembrane damage-induced autophagy. 10.1038/s41594-021-00710-6

101. Fujita N, Itoh T, Omori H, Fukuda M, Noda T, Yoshimori T. (2008). The Atg16L Complex Specifies the Site of LC3 Lipidation for Membrane Biogenesis in Autophagy. 10.1091/mbc.e07-12-1257

102. Fletcher K, Ulferts R, Jacquin E, Veith T, Gammoh N, Arasteh JM, Mayer U, Carding SR, Wileman T, Beale R, Florey O. (2018). The WD 40 domain of ATG 16L1 is required for its non-canonical role in lipidation of LC 3 at single membranes. 10.15252/embj.201797840

103. Fischer TD, Wang C, Padman BS, Lazarou M, Youle RJ. (2020). STING induces LC3B lipidation onto single-membrane vesicles via the V-ATPase and ATG16L1-WD40 domain. 10.1083/jcb.202009128

104. Skowyra ML, Schlesinger PH, Naismith TV, Hanson PI. (2018). Triggered recruitment of ESCRT machinery promotes endolysosomal repair. 10.1126/science.aar5078

105. Tan JX, Finkel T. (2022). A phosphoinositide signalling pathway mediates rapid lysosomal repair. 10.1038/s41586-022-05164-4

106. Baik SH, Ramanujan VK, Becker C, Fett S, Underhill DM, Wolf AJ. (2023). Hexokinase dissociation from mitochondria promotes oligomerization of VDAC that facilitates NLRP3 inflammasome assembly and activation. 10.1126/sciimmunol.ade7652

107. Subramanian N, Natarajan K, Clatworthy MR, Wang Z, Germain RN. (2013). The Adaptor MAVS Promotes NLRP3 Mitochondrial Localization and Inflammasome Activation. 10.1016/j.cell.2013.02.054

108. Zhou R, Yazdi AS, Menu P, Tschopp J. (2010). A role for mitochondria in NLRP3 inflammasome activation. 10.1038/nature09663

109. Iyer SS, He Q, Janczy JR, Elliott EI, Zhong Z, Olivier AK, Sadler JJ, Knepper-Adrian V, Han R, Qiao L, Eisenbarth SC, Nauseef WM, Cassel SL, Sutterwala FS. (2013). Mitochondrial Cardiolipin Is Required for Nlrp3 Inflammasome Activation. 10.1016/j.immuni.2013.08.001

110. Chen WW, Freinkman E, Sabatini DM. (2017). Rapid immunopurification of mitochondria for metabolite profiling and absolute quantification of matrix metabolites. 10.1038/nprot.2017.104

111. Bussi C, Heunis T, Pellegrino E, Bernard EM, Bah N, Dos Santos MS, Santucci P, Aylan B, Rodgers A, Fearns A, Mitschke J, Moore C, MacRae JI, Greco M, Reinheckel T, Trost M, Gutierrez MG. (2022). Lysosomal damage drives mitochondrial proteome remodelling and reprograms macrophage immunometabolism. 10.1038/s41467-022-34632-8

112. Dinkova-Kostova AT, Kostov RV, Canning P. (2017). Keap1, the cysteine-based mammalian intracellular sensor for electrophiles and oxidants. 10.1016/j.abb.2016.08.005

113. Zhao C, Gillette DD, Li X, Zhang Z, Wen H. (2014). Nuclear Factor E2-related Factor-2 (Nrf2) Is Required for NLRP3 and AIM2 Inflammasome Activation. 10.1074/jbc.m114.563114

114. Cruz CM, Rinna A, Forman HJ, Ventura ALM, Persechini PM, Ojcius DM. (2007). ATP Activates a Reactive Oxygen Species-dependent Oxidative Stress Response and Secretion of Proinflammatory Cytokines in Macrophages. 10.1074/jbc.m608083200

115. Zhong Z, Umemura A, Sanchez-Lopez E, Liang S, Shalapour S, Wong J, He F, Boassa D, Perkins G, Ali SR, McGeough MD, Ellisman MH, Seki E, Gustafsson AB, Hoffman HM, Diaz-Meco MT, Moscat J, Karin M. (2016). NF-κB Restricts Inflammasome Activation via Elimination of Damaged Mitochondria. 10.1016/j.cell.2015.12.057

116. Podinovskaia M, Prescianotto-Baschong C, Buser DP, Spang A. (2021). A novel live-cell imaging assay reveals regulation of endosome maturation. 10.7554/elife.70982

117. Ohkuma S, Poole B. (1978). Fluorescence probe measurement of the intralysosomal pH in living cells and the perturbation of pH by various agents. 10.1073/pnas.75.7.3327

118. Morgens DW, Wainberg M, Boyle EA, Ursu O, Araya CL, Tsui CK, Haney MS, Hess GT, Han K, Jeng EE, Li A, Snyder MP, Greenleaf WJ, Kundaje A, Bassik MC. (2017). Genome-scale measurement of off-target activity using Cas9 toxicity in high-throughput screens. 10.1038/ncomms15178

119. Tian S, Muneeruddin K, Choi MY, Tao L, Bhuiyan RH, Ohmi Y, Furukawa Keiko, Furukawa Koichi, Boland S, Shaffer SA, Adam RM, Dong M. (2018). Genome-wide CRISPR screens for Shiga toxins and ricin reveal Golgi proteins critical for glycosylation. 10.1371/journal.pbio.2006951

120. Selyunin AS, Iles LR, Bartholomeusz G, Mukhopadhyay S. (2017). Genome-wide siRNA screen identifies UNC50 as a regulator of Shiga toxin 2 trafficking. 10.1083/jcb.201704015

121. Mukhopadhyay S, Linstedt AD. (2012). Manganese Blocks Intracellular Trafficking of Shiga Toxin and Protects Against Shiga Toxicosis. 10.1126/science.1215930

122. Linstedt AD, Mehta A, Suhan J, Reggio H, Hauri HP. (1997). Sequence and overexpression of GPP130/GIMPc: evidence for saturable pH-sensitive targeting of a type II early Golgi membrane protein. 10.1091/mbc.8.6.1073

123. Puri S, Bachert C, Fimmel CJ, Linstedt AD. (2002). Cycling of Early Golgi Proteins Via the Cell Surface and Endosomes Upon Lumenal pH Disruption. 10.1034/j.1600-0854.2002.30906.x

124. Forrester A, Rathjen SJ, Daniela Garcia-Castillo M, Bachert C, Couhert A, Tepshi L, Pichard S, Martinez J, Munier M, Sierocki R, Renard H-F, Augusto Valades-Cruz C, Dingli F, Loew D, Lamaze C, Cintrat J-C, Linstedt AD, Gillet D, Barbier J, Johannes L. (2020). Functional dissection of the retrograde Shiga toxin trafficking inhibitor Retro-2. 10.1038/s41589-020-0474-4

125. Mukhopadhyay S, Bachert C, Smith DR, Linstedt AD. (2010). Manganese-induced Trafficking and Turnover of thecis-Golgi Glycoprotein GPP130. 10.1091/mbc.e09-11-0985

126. Puthenveedu MA, Bruns JR, Weisz OA, Linstedt AD. (2003). Basolateral Cycling Mediated by a Lumenal Domain Targeting Determinant. 10.1091/mbc.e02-10-0692

127. Sarkar S, Rokad D, Malovic E, Luo J, Harischandra DS, Jin H, Anantharam V, Huang X, Lewis M, Kanthasamy A, Kanthasamy AG. (2019). Manganese activates NLRP3 inflammasome signaling and propagates exosomal release of ASC in microglial cells. 10.1126/scisignal.aat9900

128. Yu S, Green J, Wellens R, Lopez-Castejon G, Brough D. (2020). Bafilomycin A1 enhances NLRP3 inflammasome activation in human monocytes independent of lysosomal acidification. 10.1111/febs.15619

129. Levic DS, Ryan S, Marjoram L, Honeycutt J, Bagwell J, Bagnat M. (2020). Distinct roles for luminal acidification in apical protein sorting and trafficking in zebrafish. 10.1083/jcb.201908225

130. Liu B, Carlson RJ, Pires IS, Gentili M, Feng E, Hellier Q, Schwartz MA, Blainey PC, Irvine DJ, Hacohen N. (2023). Human STING is a proton channel. 10.1126/science.adf8974

131. Muschalik N, Munro S. (2018). Golgins. 10.1016/j.cub.2018.01.006

132. Shin JJH, Crook OM, Borgeaud AC, Cattin-Ortolá J, Peak-Chew SY, Breckels LM, Gillingham AK, Chadwick J, Lilley KS, Munro S. (2020). Spatial proteomics defines the content of trafficking vesicles captured by golgin tethers. 10.1038/s41467-020-19840-4

133. Gaidt MM, Ebert TS, Chauhan D, Ramshorn K, Pinci F, Zuber S, O’Duill F, Schmid-Burgk JL, Hoss F, Buhmann R, Wittmann G, Latz E, Subklewe M, Hornung V. (2017). The DNA Inflammasome in Human Myeloid Cells Is Initiated by a STING-Cell Death Program Upstream of NLRP3. 10.1016/j.cell.2017.09.039

134. Xun J, Zhang Z, Lv B, Lu D, Yang H, Shang G, Tan JX. (2024). A conserved ion channel function of STING mediates noncanonical autophagy and cell death. 10.1038/s44319-023-00045-x

135. Welch LG, Peak-Chew S-Y, Begum F, Stevens TJ, Munro S. (2021). GOLPH3 and GOLPH3L are broad-spectrum COPI adaptors for sorting into intra-Golgi transport vesicles. 10.1083/jcb.202106115

136. Taylor RJ, Tagiltsev G, Briggs JAG. (2022). The structure of COPI vesicles and regulation of vesicle turnover. 10.1002/1873-3468.14560

137. Blackburn JB, D’Souza Z, Lupashin VV. (2019). Maintaining order: COG complex controls Golgi trafficking, processing, and sorting. 10.1002/1873-3468.13570

138. Ungar D, Oka T, Brittle EE, Vasile E, Lupashin VV, Chatterton JE, Heuser JE, Krieger M, Waters MG. (2002). Characterization of a mammalian Golgi-localized protein complex, COG, that is required for normal Golgi morphology and function. 10.1083/jcb.200202016

139. Miller EA, Barlowe C. (2010). Regulation of coat assembly—sorting things out at the ER. 10.1016/j.ceb.2010.04.003

140. Stechmann B, Bai S-K, Gobbo E, Lopez R, Merer G, Pinchard S, Panigai L, Tenza D, Raposo G, Beaumelle B, Sauvaire D, Gillet D, Johannes L, Barbier J. (2010). Inhibition of Retrograde Transport Protects Mice from Lethal Ricin Challenge. 10.1016/j.cell.2010.01.043

141. Xiao L, Magupalli VG, Wu H. (2022). Cryo-EM structures of the active NLRP3 inflammasome disc. 10.1038/s41586-022-05570-8

142. Stehlik C, Lee SH, Dorfleutner A, Stassinopoulos A, Sagara J, Reed JC. (2003). Apoptosis-Associated Speck-Like Protein Containing a Caspase Recruitment Domain Is a Regulator of Procaspase-1 Activation. 10.4049/jimmunol.171.11.6154

143. Dick MS, Sborgi L, Rühl S, Hiller S, Broz P. (2016). ASC filament formation serves as a signal amplification mechanism for inflammasomes. 10.1038/ncomms11929

144. Cai X, Chen J, Xu H, Liu S, Jiang Q-X, Halfmann R, Chen ZJ. (2014). Prion-like Polymerization Underlies Signal Transduction in Antiviral Immune Defense and Inflammasome Activation. 10.1016/j.cell.2014.01.063

145. Lu A, Magupalli VG, Ruan J, Yin Q, Atianand MK, Vos MR, Schröder GF, Fitzgerald KA, Wu H, Egelman EH. (2014). Unified Polymerization Mechanism for the Assembly of ASC-Dependent Inflammasomes. 10.1016/j.cell.2014.02.008

146. Masumoto J, Taniguchi S, Ayukawa K, Sarvotham H, Kishino T, Niikawa N, Hidaka E, Katsuyama T, Higuchi T, Sagara J. (1999). ASC, a Novel 22-kDa Protein, Aggregates during Apoptosis of Human Promyelocytic Leukemia HL-60 Cells. 10.1074/jbc.274.48.33835

147. Hung V, Udeshi ND, Lam SS, Loh KH, Cox KJ, Pedram K, Carr SA, Ting AY. (2016). Spatially resolved proteomic mapping in living cells with the engineered peroxidase APEX2. 10.1038/nprot.2016.018

148. Hammond GRV, Machner MP, Balla T. (2014). A novel probe for phosphatidylinositol 4-phosphate reveals multiple pools beyond the Golgi. 10.1083/jcb.201312072

149. Ragaz C, Pietsch H, Urwyler S, Tiaden A, Weber SS, Hilbi H. (2008). TheLegionella pneumophilaphosphatidylinositol-4 phosphate-binding type IV substrate SidC recruits endoplasmic reticulum vesicles to a replication-permissive vacuole. 10.1111/j.1462-5822.2008.01219.x

150. Glück IM, Mathias GP, Strauss S, Rat V, Gialdini I, Ebert TS, Stafford C, Agam G, Manley S, Hornung V, Jungmann R, Sieben C, Lamb DC. (2023). Nanoscale organization of the endogenous ASC speck. 10.1016/j.isci.2023.108382

151. Kohler D, Staniak M, Tsai T-H, Huang T, Shulman N, Bernhardt OM, MacLean BX, Nesvizhskii AI, Reiter L, Sabido E, Choi M, Vitek O. (2023). MSstats Version 4.0: Statistical Analyses of Quantitative Mass Spectrometry-Based Proteomic Experiments with Chromatography-Based Quantification at Scale. 10.1021/acs.jproteome.2c00834

152. Liang Z, Damianou A, Vendrell I, Jenkins E, Lassen FH, Washer SJ, Liu G, Yi G, Lou H, Cao F, Zheng X, Fernandes RA, Dong T, Tate EW, Daniel ED, Kessler BM. (2023). Proximity proteomics reveals UCH-L1 as an NLRP3 interactor that modulates IL-1β production in human macrophages and microglia. 10.1101/2023.10.09.561576

153. Fischer FA, Mies LFM, Nizami S, Pantazi E, Danielli S, Demarco B, Ohlmeyer M, Lee MSJ, Coban C, Kagan JC, Di Daniel E, Bezbradica JS. (2021). TBK1 and IKKε act like an OFF switch to limit NLRP3 inflammasome pathway activation. 10.1073/pnas.2009309118

154. Stutz A, Kolbe C-C, Stahl R, Horvath GL, Franklin BS, van Ray O, Brinkschulte R, Geyer M, Meissner F, Latz E. (2017). NLRP3 inflammasome assembly is regulated by phosphorylation of the pyrin domain. 10.1084/jem.20160933

155. Py BF, Kim M-S, Vakifahmetoglu-Norberg H, Yuan J. (2013). Deubiquitination of NLRP3 by BRCC3 Critically Regulates Inflammasome Activity. 10.1016/j.molcel.2012.11.009

156. Blander JM, Barbet G. (2018). Exploiting vita-PAMPs in vaccines. 10.1016/j.coph.2018.05.012

157. Kagan JC. (2023). Excess lipids on endosomes dictate NLRP3 localization and inflammasome activation. 10.1038/s41590-022-01364-2

158. Mateo-Tórtola M, Hochheiser IV, Grga J, Mueller JS, Geyer M, Weber ANR, Tapia-Abellán A. (2023). Non-decameric NLRP3 forms an MTOC-independent inflammasome. 10.1101/2023.07.07.548075

159. Hoffman HM, Mueller JL, Broide DH, Wanderer AA, Kolodner RD. (2001). Mutation of a new gene encoding a putative pyrin-like protein causes familial cold autoinflammatory syndrome and Muckle–Wells syndrome. 10.1038/ng756

160. Seoane PI, Beswick JA, Leach AG, Swanton T, Morris LV, Couper K, Lowe M, Freeman S, Brough D. (2023). Squaramides enhance NLRP3 inflammasome activation by lowering intracellular potassium. 10.1038/s41420-023-01756-9

161. Ran L, Ye T, Erbs E, Ehl S, Spassky N, Sumara I, Zhang Z, Ricci R. (2023). KCNN4 links PIEZO-dependent mechanotransduction to NLRP3 inflammasome activation. 10.1126/sciimmunol.adf4699

162. Ren G-M, Li J, Zhang X-C, Wang Y, Xiao Y, Zhang X-Y, Liu X, Zhang W, Ma W-B, Zhang J, Li Y-T, Tao S-S, Wang T, Liu K, Chen H, Zhan Y-Q, Yu M, Li C-Y, Ge C-H, Tian B-X, Dou G-F, Yang X-M, Yin R-H. (2021). Pharmacological targeting of NLRP3 deubiquitination for treatment of NLRP3-associated inflammatory diseases. 10.1126/sciimmunol.abe2933

163. Gregory GE, Munro KJ, Couper KN, Pathmanaban ON, Brough D. (2023). The NLRP3 inflammasome as a target for sensorineural hearing loss. 10.1016/j.clim.2023.109287

164. Youm Y-H, Grant RW, McCabe LR, Albarado DC, Nguyen KY, Ravussin A, Pistell P, Newman S, Carter R, Laque A, Münzberg H, Rosen CJ, Ingram DK, Salbaum JM, Dixit VD. (2013). Canonical Nlrp3 Inflammasome Links Systemic Low-Grade Inflammation to Functional Decline in Aging. 10.1016/j.cmet.2013.09.010

165. Bauernfeind F, Niepmann S, Knolle PA, Hornung V. (2016). Aging-Associated TNF Production Primes Inflammasome Activation and NLRP3-Related Metabolic Disturbances. 10.4049/jimmunol.1501336

166. Camell CD, Günther P, Lee A, Goldberg EL, Spadaro O, Youm Y-H, Bartke A, Hubbard GB, Ikeno Y, Ruddle NH, Schultze J, Dixit VD. (2019). Aging Induces an Nlrp3 Inflammasome-Dependent Expansion of Adipose B Cells That Impairs Metabolic Homeostasis. 10.1016/j.cmet.2019.10.006

167. He M, Chiang H-H, Luo H, Zheng Z, Qiao Q, Wang L, Tan M, Ohkubo R, Mu W-C, Zhao S, Wu H, Chen D. (2020). An Acetylation Switch of the NLRP3 Inflammasome Regulates Aging-Associated Chronic Inflammation and Insulin Resistance. 10.1016/j.cmet.2020.01.009

168. Reaves B, Horn M, Banting G. (1993). TGN38/41 recycles between the cell surface and the TGN: brefeldin A affects its rate of return to the TGN. 10.1091/mbc.4.1.93

169. MS L, KE H. (1992). The trans-Golgi network can be dissected structurally and functionally from the cisternae of the Golgi complex by brefeldin A

170. Tewari R, Bachert C, Linstedt AD. (2015). Induced oligomerization targets Golgi proteins for degradation in lysosomes. 10.1091/mbc.e15-04-0207

171. Wong CH, Wingett SW, Qian C, Hunter MR, Taliaferro JM, Ross-Thriepland D, Bullock SL. (2024). Genome-scale requirements for dynein-based transport revealed by a high-content arrayed CRISPR screen. 10.1083/jcb.202306048

172. Magupalli VG, Negro R, Tian Y, Hauenstein AV, Di Caprio G, Skillern W, Deng Q, Orning P, Alam HB, Maliga Z, Sharif H, Hu JJ, Evavold CL, Kagan JC, Schmidt FI, Fitzgerald KA, Kirchhausen T, Li Y, Wu H. (2020). HDAC6 mediates an aggresome-like mechanism for NLRP3 and pyrin inflammasome activation. 10.1126/science.aas8995

173. Wang L, Shi S, Unterreiner A, Kapetanovic R, Ghosh S, Sanchez J, Aslani S, Xiong Y, Hsu C-L, Donovan KA, Farady CJ, Fischer ES, Bornancin F, Matthias P. (2024). HDAC6/aggresome processing pathway importance for inflammasome formation is context-dependent. 10.1016/j.jbc.2024.105638

174. Xu L, Sowa ME, Chen J, Li X, Gygi SP, Harper JW. (2008). An FTS/Hook/p107^FHIP^Complex Interacts with and Promotes Endosomal Clustering by the Homotypic Vacuolar Protein Sorting Complex. 10.1091/mbc.e08-05-0473

175. Christensen JR, Kendrick AA, Truong JB, Aguilar-Maldonado A, Adani V, Dzieciatkowska M, Reck-Peterson SL. (2021). Cytoplasmic dynein-1 cargo diversity is mediated by the combinatorial assembly of FTS–Hook–FHIP complexes. 10.7554/elife.74538

176. Redwine WB, DeSantis ME, Hollyer I, Htet ZM, Tran PT, Swanson SK, Florens L, Washburn MP, Reck-Peterson SL. (2017). The human cytoplasmic dynein interactome reveals novel activators of motility. 10.7554/elife.28257

177. Szebenyi G, Wigley WC, Hall B, Didier A, Yu M, Thomas P, Krämer H. (2007). Hook2 contributes to aggresome formation. 10.1186/1471-2121-8-19

178. Venditti R, Rega LR, Masone MC, Santoro M, Polishchuk E, Sarnataro D, Paladino S, D’Auria S, Varriale A, Olkkonen VM, Di Tullio G, Polishchuk R, De Matteis MA. (2019). Molecular determinants of ER–Golgi contacts identified through a new FRET–FLIM system. 10.1083/jcb.201812020

179. D’Angelo G, Rega LR, De Matteis MA. (2012). Connecting vesicular transport with lipid synthesis: FAPP2. 10.1016/j.bbalip.2012.01.003

180. Cho NH, Cheveralls KC, Brunner A-D, Kim K, Michaelis AC, Raghavan P, Kobayashi H, Savy L, Li JY, Canaj H, Kim JYS, Stewart EM, Gnann C, McCarthy F, Cabrera JP, Brunetti RM, Chhun BB, Dingle G, Hein MY, Huang B, Mehta SB, Weissman JS, Gómez-Sjöberg R, Itzhak DN, Royer LA, Mann M, Leonetti MD. (2022). OpenCell: Endogenous tagging for the cartography of human cellular organization. 10.1126/science.abi6983

181. Foulquier F, Amyere M, Jaeken J, Zeevaert R, Schollen E, Race V, Bammens R, Morelle W, Rosnoblet C, Legrand D, Demaegd D, Buist N, Cheillan D, Guffon N, Morsomme P, Annaert W, Freeze HH, Van Schaftingen E, Vikkula M, Matthijs G. (2012). TMEM165 Deficiency Causes a Congenital Disorder of Glycosylation. 10.1016/j.ajhg.2012.05.002

182. Hsu H-Y, Wen M-H. (2002). Lipopolysaccharide-mediated Reactive Oxygen Species and Signal Transduction in the Regulation of Interleukin-1 Gene Expression. 10.1074/jbc.m111883200

183. Cameron AM, Castoldi A, Sanin DE, Flachsmann LJ, Field CS, Puleston DanielJ, Kyle RL, Patterson AE, Hässler F, Buescher JM, Kelly B, Pearce EL, Pearce EJ. (2019). Inflammatory macrophage dependence on NAD+ salvage is a consequence of reactive oxygen species–mediated DNA damage. 10.1038/s41590-019-0336-y

184. Branon TC, Bosch JA, Sanchez AD, Udeshi ND, Svinkina T, Carr SA, Feldman JL, Perrimon N, Ting AY. (2018). Efficient proximity labeling in living cells and organisms with TurboID. 10.1038/nbt.4201

185. Deutsch EW, Bandeira N, Perez-Riverol Y, Sharma V, Carver JJ, Mendoza L, Kundu DJ, Wang S, Bandla C, Kamatchinathan S, Hewapathirana S, Pullman BS, Wertz J, Sun Z, Kawano S, Okuda S, Watanabe Y, MacLean B, MacCoss MJ, Zhu Y, Ishihama Y, Vizcaíno JA. (2022). The ProteomeXchange consortium at 10 years: 2023 update. 10.1093/nar/gkac1040

186. Perez-Riverol Y, Csordas A, Bai J, Bernal-Llinares M, Hewapathirana S, Kundu DJ, Inuganti A, Griss J, Mayer G, Eisenacher M, Pérez E, Uszkoreit J, Pfeuffer J, Sachsenberg T, Yılmaz Ş, Tiwary S, Cox J, Audain E, Walzer M, Jarnuczak AF, Ternent T, Brazma A, Vizcaíno JA. (2018). The PRIDE database and related tools and resources in 2019: improving support for quantification data. 10.1093/nar/gky1106

187. Schmid-Burgk JL, Schmidt T, Gaidt MM, Pelka K, Latz E, Ebert TS, Hornung V. (2014). OutKnocker: a web tool for rapid and simple genotyping of designer nuclease edited cell lines. 10.1101/gr.176701.114

188. Schindelin J, Arganda-Carreras I, Frise E, Kaynig V, Longair M, Pietzsch T, Preibisch S, Rueden C, Saalfeld S, Schmid B, Tinevez J-Y, White DJ, Hartenstein V, Eliceiri K, Tomancak P, Cardona A. (2012). Fiji: an open-source platform for biological-image analysis. 10.1038/nmeth.2019

189. Stirling DR, Swain-Bowden MJ, Lucas AM, Carpenter AE, Cimini BA, Goodman A. (2021). CellProfiler 4: improvements in speed, utility and usability. 10.1186/s12859-021-04344-9

190. Schweppe DK, Eng JK, Yu Q, Bailey D, Rad R, Navarrete-Perea J, Huttlin EL, Erickson BK, Paulo JA, Gygi SP. (2020). Full-Featured, Real-Time Database Searching Platform Enables Fast and Accurate Multiplexed Quantitative Proteomics. 10.1021/acs.jproteome.9b00860

191. Eng JK, Jahan TA, Hoopmann MR. (2012). Comet: An open-source MSMS sequence database search tool. 10.1002/pmic.201200439

192. Beausoleil SA, Villén J, Gerber SA, Rush J, Gygi SP. (2006). A probability-based approach for high-throughput protein phosphorylation analysis and site localization. 10.1038/nbt1240

193. Huttlin EL, Jedrychowski MP, Elias JE, Goswami T, Rad R, Beausoleil SA, Villén J, Haas W, Sowa ME, Gygi SP. (2010). A Tissue-Specific Atlas of Mouse Protein Phosphorylation and Expression. 10.1016/j.cell.2010.12.001

194. Elias JE, Gygi SP. (2009). Target-Decoy Search Strategy for Mass Spectrometry-Based Proteomics. 10.1007/978-1-60761-444-9_5

195. Elias JE, Gygi SP. (2007). Target-decoy search strategy for increased confidence in large-scale protein identifications by mass spectrometry. 10.1038/nmeth1019

196. Savitski MM, Wilhelm M, Hahne H, Kuster B, Bantscheff M. (2015). A Scalable Approach for Protein False Discovery Rate Estimation in Large Proteomic Data Sets. 10.1074/mcp.m114.046995

